# Resistance to Atrial Fibrillation Domestication and Mitochondrial Dysfunction in Sheep: a potential key role of the TCA Cycle and mitochondrial redox state

**DOI:** 10.1101/2025.06.01.657249

**Authors:** Guido Caluori, Stanley Nattel, Benoît Pinson, Patrice Naud, Bertrand Beauvoit, Stéphane Claverol, Fanny Vaillant, Sabine Charron, Florent Guilloteau, Virginie Loyer, Marion Constantin, Miruna Popa, Andrei Belykh, Andreas Haeberlin, Sylvain Ploux, Hassan Adam Mahamat, Rémi Dubois, Bastien Guillot, Philipp Krisai, Tsukasa Kamakura, Olivier Bernus, Pierre Jaïs, Pierre Dos Santos, Philippe Pasdois

## Abstract

**Background:** Atrial fibrillation (AF) often progresses from paroxysmal to more stable forms. It is well-recognized that patients vary in their AF progression, but underlying mechanisms remain unclear. This work, performed in a sheep AF-model, aimed to identify atrial redox and energetic status differences between animals developing stable AF (AF-S) versus those resistant to AF-stabilization (AF-R).

**Methods:** AF was monitored with telemetry and maintained with bursts of atrial tachystimulation whenever sinus rhythm resumed. Electrophysiological remodeling was assessed *via* contact mapping. Structural remodeling was described by histology. Proteomic, metabolomic, enzymatic and bioenergetic remodeling were evaluated using frozen left atrial appendage (LAA) tissues and isolated LAA mitochondria. Healthy young rats were used to investigate if an induced metabolic challenge could stabilize AF episodes upon transesophageal atrial tachypacing challenge.

**Results:** AF-S sheep developed stable AF (>24-hours self-sustained) after 13 days on average, whereas AF-R sheep failed to develop self-sustained AF despite 120 days of electrically-maintained AF. Contact mapping and histological analysis revealed similar electro-structural remodeling in both groups. Metabolic analysis showed significant differences in tricarboxylic acid (TCA) cycle enzymes activities and a 45% increase in AF-S LAA succinate content versus AF-R. AF-S mitochondria showed abnormal mitochondrial succinate oxidation, associated with a significant 20% decrease in ATP synthesis rate, 22% increase in ROS emission and mitochondrial inner membrane hyperpolarization. The ratios of ATP to ADP, NAD^+^ to NADH, and Complex I/II were disturbed in AF-S compared to AF-R. Calculated mitochondrial NAD^+^ to NADH ratios suggest a reduced state of in-vivo AF-R mitochondria compared to the oxidized state of AF-S. Exogenous succinate was metabolized when incubated with rat atrial cardiomyocytes and altered redox balance, while intravenous succinate stabilized atrial arrhythmias induced by tachypacing *in vivo*.

**Conclusions:** Sheep resistant to AF-progression showed specific TCA cycle, energetic and redox adaptations compared to animals that developed self-sustained AF. In this animal model, mitochondrial TCA cycle remodeling and associated redox and energetic responses determined the resistance to AF domestication, with potential relevance to identify new mechanistic determinants of AF progression in humans.

## Introduction

Atrial fibrillation (AF) leads to significant population morbidity and mortality [1]. Allessie et al provided the paradigm-shifting observation that AF episodes produce progressive atrial remodeling with the eventual development of spontaneously persisting AF in goats [2]. In patients, the transition rate from paroxysmal to permanent AF has been reported to vary between 15% to more than 50% within a follow-up period ranging from 1 to 12 years [3–7]. Many patients with paroxysmal AF never progress to persistent AF. The biological mechanisms behind this AF resistance remain poorly understood, despite significant clinical importance.

Studies on human atria have revealed metabolic alterations with AF, including changes in the mitochondrial electron transport chain [8–10], increased reactive oxygen species (ROS) production [11,12], altered glucose and fat metabolism [9,13], and oxidative damage to mitochondrial DNA [14]. Whether differences in the metabolic response are linked to AF resistance is unknown.

To study mechanisms of differences in resistance to AF development in an animal model, we implemented a sheep model of AF induced by intermittent rapid atrial pacing (RAP) [15,16], similar to Allessie’s initial approach in goats [2]. We observed that a fraction of the instrumented sheep, despite 4 months of RAP exposure without loss of capture, never developed self-sustained AF ("AF resistant group", AF-R), while the majority did develop self-sustained AF ("AF sensitive group", AF-S), within three weeks of RAP onset. In this report, we will show how our findings suggest that the mitochondrial metabolic response plays a key role in determining resistance to AF.

## Methods

Experiments followed Directive 2010/63/EU (European Parliament) on animal-use for scientific research. Protocols were approved by the local ethical committee (CEEA50) at University of Bordeaux and by the French Government (APAFIS#20364 and APAFIS#29528, for sheep and rat, respectively).

### Sheep AF model

This study used a modified version of the atrial fibrillation (AF) model described by Jalife and colleagues [15]. Sheep were prepared for the surgical procedure after an intramuscular injection of ketamine (20 mg/Kg), acepromazine (0.1 mg/Kg) and buprenorphine (9 µg/Kg).

Anesthesia was induced by an intravenous bolus dose of propofol (1 mg/Kg) *via* a venous catheter in the foreleg, followed by endotracheal intubation and ventilation with a 1:1 mixture of room air and oxygen containing isoflurane (1.5 to 3%, as needed to maintain deep anesthesia). Thirty-one female sheep (Charmoise, mean age = 8.4±0.1 years, mean weight = 50.8±1.2 kg) were implanted with a dual chamber pacemaker (St. Jude Medical, Inc, St Paul, MN), two pacing leads (IS1 leads Tendril STS 2088TC, St. Jude Medical, Inc, St Paul, MN), and a telemetry device (EasyTEL+L – ETA, Emka Technologies, Paris, FR). After 7 days of recovery, the pacemaker was randomly activated to either sense and record the right atrial and ventricular signals (Sham group, N=11) or to sense, record and pace the right atria (Supplemental Figure 1A) to induce AF (AF-induction group, N=20). In the AF-induction group, bipolar square-wave atrial stimuli at 5-times voltage threshold and 10-Hz frequency were used to induce AF whenever sinus rhythm (SR) was detected. Tachypacing was delivered for 30 seconds, after which a 7-second pause was allowed to determine the spontaneous rhythm.

Among the animals in the AF-induction branch, two groups of sheep emerged, one developing stable AF (AF-S), and one resistant to AF progression (AF-R).

### Atrial endocardial electrical contact mapping

Electroanatomical mapping was performed with a CARTO™ 3 system using CARTO™ v6 software (Biosense Webster, Irvine, CA, USA), and a 20-pole steerable mapping catheter (PENTARAY™ NAV, Biosense Webster). Conduction velocity, (CV) and local effective refractory period (ERP) were calculated in the right and left atrium with programmed electrical pacing. Sinus rhythm was restored by direct-current cardioversion (1 shock, 50-70J) of AF-S animals prior to contact mapping.

### Sheep euthanasia and heart explantation

After anesthesia (see above) a median thoracotomy was performed and the heart was exposed. Mechanical activity was stopped by aortic retroperfusion with ice-cold Ringer-Lactate solution at a pressure of 80 mmHg, and by filling the chest cavity with ice-cold NaCl solution (9 g/L). After complete loss of mechanical activity, disappearance of a detectable arterial pressure and total absence of response to pain, the heart was explanted.

### Sheep atrial histology and structural characterization

The left atrial posterior wall was fixed in 4% paraformaldehyde in phosphate-buffered saline. Thin paraffin-embedded sections (5-µm) were stained with Masson trichrome. The slides were scanned with an Axio Scan.Z1 microscope (ZEISS, Jena, Germany) with 20x magnification in brightfield and the tiles automatically assembled to form the section overview. The images were imported into Fiji [17] for calculation of endomysial fibrosis, cardiomyocyte cross-section area and cell-to-cell distance.

### Proteomics

*nLC-MS/MS analysis and Label-Free Quantitative Data Analysis*. Following sample preparation and protein digestion (see Methods in Online Supplement), peptide mixture was analyzed on an Ultimate 3000 nanoLC system (Dionex, Amsterdam, Netherlands) coupled to an Electrospray Orbitrap Fusion™ Lumos™ Tribrid™ Mass Spectrometer (Thermo Fisher Scientific, San Jose, CA). Separation flow-rate was set at 300 nL/min. The mass spectrometer operated in positive-ion mode at a 2-kV needle voltage. Data were acquired using Xcalibur

### 4.4 software

#### Proteomic analysis

Data were searched by SEQUEST through Proteome Discoverer 2.5 (Thermo Fisher Scientific Inc.) against an *Ovis aries* uniprot database (21,195 entries in v2022-04). Spectra from peptides higher than 5000 Da or lower than 350 Da were rejected. Precursor Detector node was included. Peptide validation was performed using Percolator algorithm [18] and only “high confidence” peptides were retained corresponding to a 1% False Positive Rate at peptide level. Peaks were detected and integrated using the Minora algorithm embedded in Proteome Discoverer. Normalization was performed based on total *Ovis aries* protein amount. Quantitative data were considered for proteins quantified by a minimum of two peptides and a statistical *p*-value lower than 0.05 after correction for multiple comparisons with the Benjamini-Hochberg method.

#### Bioinformatics

All statistical analyses of proteomic data were conducted using RStudio (Version 2026,06,1, RStudio, PBC), the online platforms Reactome [19] and Metaboanalyst 6.0 [20]. We maintained an R Project to organize scripts, datasets, and outputs systematically, with R version 4.2.2. Batch effects were corrected using the ’limma’ package in R. Data wrangling was performed with the ’dplyr’ package to streamline the datasets into structured formats amenable to downstream analysis. Proteins were annotated with biological pathways using Wikipathway database. The original data set and analysis is available on the following link: https://doi.org/10.6084/m9.figshare.28269872.v1

#### Rat euthanasia and heart explant

Anesthesia was initially induced in male Wistar rats (225-300 g) with 3.5% isoflurane for 3 minutes. For heart explantation, rats were anticoagulated (heparin 1000 IU/Kg) and left in the induction chamber with 3.5% isoflurane for a further 3 minutes before euthanasia. After confirmation of the lack of pain reflex by pinching the lower paws and the tail, euthanasia was performed by cervical dislocation. The heart was removed after death confirmation by the absence of respiration.

### Rat atrial cardiomyocyte (RACM) isolation

Male Wistar rats (225-300g, N=10) were subjected to RACM isolated by collagenase digestion after cardiac explantation (see Online Supplement).

### Steady-state ROS production in RACM

Freshly-isolated RACMs were allowed to sediment in Tyrode solution on a 35 mm Flurodish (WPI, Sarasota, FL-US) at RT for 1h. They were then incubated for 30 min under 95% room-air/5% CO_2_ at 37°C in the presence of 5 µmol/L CellROX ™ Green (Thermo Fisher Scientific, MA-US). After incubation, brightfield and epifluorescence images were obtained with a Leica Thunder Imager system (Leica Camera, Wetzlar, Germany) and 488/530 excitation/emission filters. To observe the effect of exogenous succinate application on intracellular ROS production, RACMs were incubated with 5 mmol/L succinate (buffered with HEPES 25 mmol/L at pH 7.4 RT). RACMs were also incubated with Tyrode’s solution and H_2_O_2_ (50 µmol/L), as negative and positive control respectively.

### RACM incubation with exogenous succinate for metabolomic analysis

Freshly isolated RACM (N=4) were equally divided into two 1.5 ml conical vials each containing 1 ml of cell suspension. One vial was supplemented with succinate (buffered with HEPES 25 mmol/L at pH 7.4 RT) to reach a final concentration of 5 mmol/L. The paired vial was supplemented with Tyrode buffered with HEPES 25 mmol/L at pH 7.4 RT. Both vials were placed in an incubator at 37°C under gentle agitation for 30 minutes. Subsequently, a cell pellet was obtained by spinning the cell solution at 100g for 3 mins at 4°C. The supernatant was removed and the pellet was gently resuspended in fresh ice-cold Tyrode to dilute the residual succinate. This operation was repeated twice. The final dry pellet was snap frozen in liquid nitrogen and stored at -80°C for mass spectrometry analysis.

### Rat Langendorff perfusion and surface fluorescence measurements

Hearts from male Wistar Rats (225-300g, N=5) were cannulated and perfused in Langendorff mode and an optic-fiber cable, connected to a modified spectrofluorometer (Xenius, SAFAS Monaco), was placed in front of the LAA [21]. The NAD(P)H pool redox state was assessed on the beating LAA by recording autofluorescence (340/460nm). To observe mitochondrial H_2_O_2_ emission, hearts were loaded with Mitochondria Peroxy-Yellow 1 (3 µmol/L MitoPY1, 485/535nm, Tocris Bioscience, #4428) [22]. Succinate (pH 7.4) was then perfused at 5 mmol/L with a syringe pump for 10 min. A second washout was then performed before starting H_2_O_2_ perfusion (400 µmol/L) for 15 min. Distance and motion artefacts were diminished by dividing the emitted photon intensity by reflected signals measured at the respective excitation and emission wavelengths [23].

### In-vivo assessment of susceptibility to atrial arrhythmia in the rat

Anesthesia was induced in male Wistar rats (250-300g, N=20) with isoflurane (3.5%) and they were transferred supine to a heated mat equipped with an anesthesia mask (providing 80% oxygen/20% room-air with 2.5% isoflurane (Minerve, Esternay, France)). A CIB’ER Mouse® catheter (NuMED, Inc., Hopkinton, NY-USA) was introduced into the esophagus until an atrial electrogram was recorded. A 26G cannula inserted *via* the caudal vein was connected to a syringe pump for randomly assigned IV infusion of vehicle or succinate solution (80 mg/mL) at 15 mL/kg/h. In some experiments, 10 µmol/kg S1QEL1.1 was injected intraperitoneally at least 5 minutes prior to IV infusion onset. Atrial tachyarrhythmia (AF/AT) vulnerability was quantified 10 min after IV injection with 10 RAP bursts at 30 Hz for 30 sec, spaced 5-10 seconds to visually assess the presence of sinus rhythm or AF/AT. In a parallel set of experiments (N=6 Wistar rats), the same *in vivo* atrial tachypacing protocol was applied and serum collected in order to perform a targeted metabolomic analysis by mass spectrometry. Rat serum was obtained after cannulation of the ventral tail artery, from which 1-2 ml of blood were drawn in a non-treated conical vial. The blood was allowed to clot at RT for 20 mins, before being centrifuged twice at 2000g, 4°C, 10 mins. Serum aliquots were snap frozen in liquid nitrogen and stored at -80°C. For baseline paired serum collection 500µL of blood was taken before starting the atrial tachypacing sequence.

### Sheep LAA mitochondrial isolation and purification

Sheep LAA mitochondrial fraction was isolated and purified as previously published [24] with modifications. Following heart excision, the entire sheep LAA was dissected free and chopped into fine pieces before incubation at 4°C for 45 min while stirring in 30 mL of buffer containing (mmol/L): CaK_2_EGTA 2.77; K_2_EGTA 7.23 (pCa=7); MgCl_2_ 6.56; dithiothreitol (DTT) 0.5; MES 50; imidazole 20; taurine 20; Na_2_ATP 5.3; creatine phosphate 15, and 1 mg/mL collagenase type-2 (Worthington, Lakewood, NJ-US); pH 7.3 at 4°C with KOH. The resulting tissue suspension was used to isolate and purify mitochondria. Mitochondria were kept on ice at 80-100 mg/mL (Biuret assay) for <6 hours pre-study or frozen in liquid-N_2_ and stored at -80°C for later biochemical analysis

### Measurements of respiration, H_2_O_2_ emission, NAD(P)H autofluorescence, and ATP synthesis in isolated sheep LAA mitochondria

Mitochondrial function was studied in different bioenergetic states: basal (respiratory substrates only, state 2), state 3 (ADP 1.5 mmol/L) and after ATP-synthesis inhibition (state 4, carboxyatractlyloside, CAT, 5 µmol/L). Oxygen-consumption was measured by oxygraphy at 37°C (Digital Model 10, Rank Brothers and PowerLab, ADInstruments, Sidney, Australia). Mitochondria (0.25 mg/mL) were added to a buffer containing (mmol/L): mannitol 140, KCl 100, EGTA 1, MgCl_2_ 20, KH_2_PO_4_ 10, Amplex^®^ Red 15 µmol/L, horseradish peroxidase 12 µg/ml, and respiratory substrates. The respiratory chain was fueled with different substrate combinations, (i) GMS: glutamate (5 mmol/L) + malate (2 mmol/L) + succinate (5 mmol/L), (ii) GMS+S1QEL1.1 (210 nmol/L) (iii) PCM: palmitoyl-carnitine (15 µmol/L) + malate (2 mmol/L), or (iv) PM: pyruvate (5 mmol/L) + malate (2 mmol/L). Oxygen-consumption rates are expressed in natO/min/mg protein. The cap of the oxygraphic cell was equipped with an optic fiber coupled to a fluorometer (Xenius, Safas Monaco), and with a glass pH microelectrode (ThermoFisher Scientific). Autofluorescence of NAD(P)H (340/460 nm) and resorufin (570/600 nm) fluorescence were collected through the optic fiber. During ATP-synthesis, pH increase was converted into H^+^-consumption after HCl titration (Titripur Reagents, Merck). H^+^-consumption flux was expressed in nmol H^+^/min/mg protein and converted into ATP-synthesis flux, expressed in nmol of ATP/min/mg protein according to published methodology [25,26]. Oxidative phosphorylation yield, P to O ratio, was expressed as the ratio of maximal ATP-synthesis to oxygen consumption rates measured during ATP-synthesis. The respiratory control was expressed as the ratio of state 3 to state 2 respiration.

### Measurement of sheep LAA mitochondrial membrane potential

Mitochondrial membrane potential (ΔΨ_m_) was measured with a tetraphenylphosphonium (TPP^+^)-sensitive electrode (O2k, OROBOROS Instruments, Austria). Calibration was performed before each run with known quantity of TPP^+^ (2 µmol/L maximum according to published methodology [27]) in the respiration buffer supplemented with respiratory substrates (PM, GMS or PCM). ΔΨ_m_ was quantified according to published methodology [28] with respiratory substrates only, after ADP addition (1.5 mmol/L) followed by CAT (5 µmol/L) on a solution containing 0.2 mg/mL mitochondria. The mitochondrial matrix volume was set to 1.1 µL/mg [29]. Non-specific TPP^+^ mitochondrial binding was corrected for by adding 2 µmol/L carbonyl cyanide 4-(trifluoromethoxy)phenylhydrazone (FCCP) at the end of each run.

### Sample preparation for biochemical analysis

*Metabolite extraction:* Metabolites were either extracted with an acidic methodology or using an ethanol boiling methodology adapted from [30]. For the acidic approach, pieces of *in vivo* freeze-clamped LAA were pulverized under liquid-N_2_, and 100-150 mg were mixed with 500 µL cold perchloric acid (10%) solution supplemented with EDTA (10 mmol/L) and neutralized (pH 7.2-7.4 with KHCO_3_ and KOH-MOPS). For the ethanol-based methodology each rat plasma sample (100 µL) was mixed with 900 µL of extraction buffer (ethanol/HEPES 0.1 mM, pH 7.0, in a 4:1 ratio). For rat atrial cardiomyocyte pellets and sheep LAA frozen powder, samples were resuspended in 1 mL of the same extraction buffer and sheep LAA were homogenized using a 5 ml glass/PTFE Potter-Elvehjem grinder. For rat plasma and atrial cardiomyocytes, succinic acid-2,2,3,3-D4 (a deuterium-labeled succinic acid in which the four methylene hydrogens are replaced by deuterium; #293075, Sigma-Aldrich) was added to the biological extracts at a final concentration of 1 nmol/mL in the extraction buffer. This compound was utilized as an internal standard for normalization during mass spectrometry analyses. All biological materials were subsequently heated at 80°C for 3 minutes and dried using a rotavapor (140 rpm) at 65°C for 3 minutes. The resulting residues were reconstituted in either 1 mL of pure water (for plasma), 400 µL (for atrial cardiomyocytes), or 10 µL/mg tissue (for sheep LAA). Insoluble material was removed by centrifugation at 21,000g for 1 hour at 4°C. The clarified supernatant was subjected to ultrafiltration using Nanosep 10K Omega filters (#OD010C35, Pall).

To evaluate specific activities of cytosolic enzymes, 50-100 mg of frozen pulverized LAA were mixed with 200-400 µL of buffer containing 50 mmol/L KH_2_PO_4_, 50 µmol/L DTT, 0.3% w/v Tx-100, protease inhibitors (Roche cOmplete™), pH 7.2 at RT. For mitochondrial enzymes activity assessment, frozen isolated LAA mitochondria were solubilized in the same buffer at a concentration ranging from 0.2 to 4mg/ml depending on the enzymes studied.

### Sheep LAA metabolite content assessment

LAA metabolite content analysis was performed either by Liquid Chromatography Coupled to Conductimetry, UV absorbance or Mass Spectrometry Detectors or by enzymatic quantification from in vivo freeze-clamped LAA tissue. The detailed methodology is available in the online supplement.

### Assessment of mitochondrial and cytosolic enzyme specific activities

All specific activities were measured at 37°C by spectrophotometry (V-760, JASCO UK Limited) or fluorimetry (FP-8500, JASCO UK), unless otherwise stated. The detailed procedure for enzymatic assay is available in the online supplement. The following enzymes activities were assessed citrate synthase (CS), NADH:decylubiquinone oxidoreductase (Complex I), succinate:ubiquinone oxidoreductase (Complex II), pyruvate dehydrogenase (PDH), alpha-ketoglutarate dehydrogenase (𝛼-KGDH), succinyl-CoA synthetase (SCS), fumarase, malate dehydrogenase isoform 2 (MDH2), and D/L hydroxyglutarate dehydrogenase (2-HGDH).

### Measurement of carbonyl content in sheep LAA tissue and rat heart

*Spectrophotometric methodology in sheep atrial and rat heart samples:* 60-80 mg of sheep LAA or rat heart frozen tissue powder were solubilized in guanidine solution (6 mol/L). 200 µL of homogenate were mixed with 800µL of 12.5 mmol/L 2,4-Dinitrophenylhydrazine (DNPH) solution or with 800 µL of 2.5 mol/L HCl and shaken for 1 hour in the dark.

Following incubation, proteins were precipitated with 2 cycles of trichloroacetic acid treatment at 20 and 10%. Protein pellets (10,000g 10 min) were washed three times, with 50/50 ethyl-acetate-ethanol solution. The final pellets were resuspended in 300 µL guanidine solution (6 mol/L). Sample absorbance was assessed at 370 nm (Xenius, SAFAS Monaco). Protein concentration was determined using a bicinchoninic acid assay (ThermoFisher Scientific, #23227). Total protein carbonyl content was calculated considering the background absorbance and expressed in nmol/mg protein.

#### Western-Blot methodology on rat heart samples

Aliquots of 30 μg total protein previously extracted from the frozen rat heart tissue powder were diluted in a 15 μL volume of 6% (wt/v) SDS. An equal amount of 15 μL of 20 mM DNPH in 10% (v/v) trifluoroacetic acid was added into the protein solution and incubated at RT for 10 mins. The solution was then neutralized with 15 μL of 2 M Tris in 30% (v/v) glycerol, containing 7% (v/v) β-mercaptoethanol. The reacted protein solution was then processed for conventional Western Blot analysis, using goat anti-2,4-dinitrophenyl (DNP) as primary antibody (1:5000, A150117A, Bethyl Laboratories, Inc.), and Rabbit Anti-Goat IgG (H+L)-HRP Conjugate (1:5000 dilution, 1721034, Bio Rad Laboratories).

#### Metabolite compartmentation analysis at equilibrium

The [NAD^+^] to [NADH] ratio, oxaloacetate and alpha-ketoglutarate concentration within the cytosolic and mitochondrial compartments were calculated assuming that the system was at thermodynamic equilibrium. The same approach was used to calculate the proton motive force (Δp) and tissue [NAD^+^] to [NADH] ratio. The equations and values used for the ΔG°’ calculation are presented in the online supplement.

### Statistical Analysis

All statistics were performed using GraphPad Prism. Data are presented as mean±SD. Comparison were performed with non-parametric tests, because (i) the number of individual experiments varied from 3 to 13, (ii) these tests can be used in presence of outliers (the sheep model displaying important variability for some parameters, especially in the Sham group), and (iii) evaluation of normality and homoscedasticity of individual samples are not required. For comparison of multiple independent samples or two independent samples, the non-parametric Kruskal-Wallis test with Dunn’s correction for multiple comparisons and non- parametric multiple t-tests with the Holm-Sidak correction for multiple comparisons were used, respectively. For comparison of two groups of unpaired or paired samples, Mann-Whitney or Wilcoxon tests were used respectively. For survival curve analysis, we used the Kaplan-Meyer model. Differences were considered significant for a *p*-value <0.05; exact *p* values are shown in figures.

## Results

### LAA Electrical and structural remodeling

AF-S sheep developed spontaneously maintained AF (defined by episodes>24-hours) after an average of 13 days (range 5-22, Figures 1A and B and Supplemental Figure 1A), while AF-R sheep never did along the follow-up window (Figures 1A and C and supplemental Figure 1B). No adverse events were observed that required changes to the pacing protocol (see methods section for more details). Telemetry showed that 60% of AF-S sheep maintained sustained AF with no bursts required during the follow-up period, while the remaining 40% received between 1 and 21 bursts (Supplemental Figure 1C) during the entire follow-up period, representing less than 0.1% of daily time in sinus rhythm during the stable AF period. In contrast, AF-R sheep received on average 2300 bursts per 24 hours and never showed self-sustained AF (Supplemental Figure 1D).

**Figure 1.**
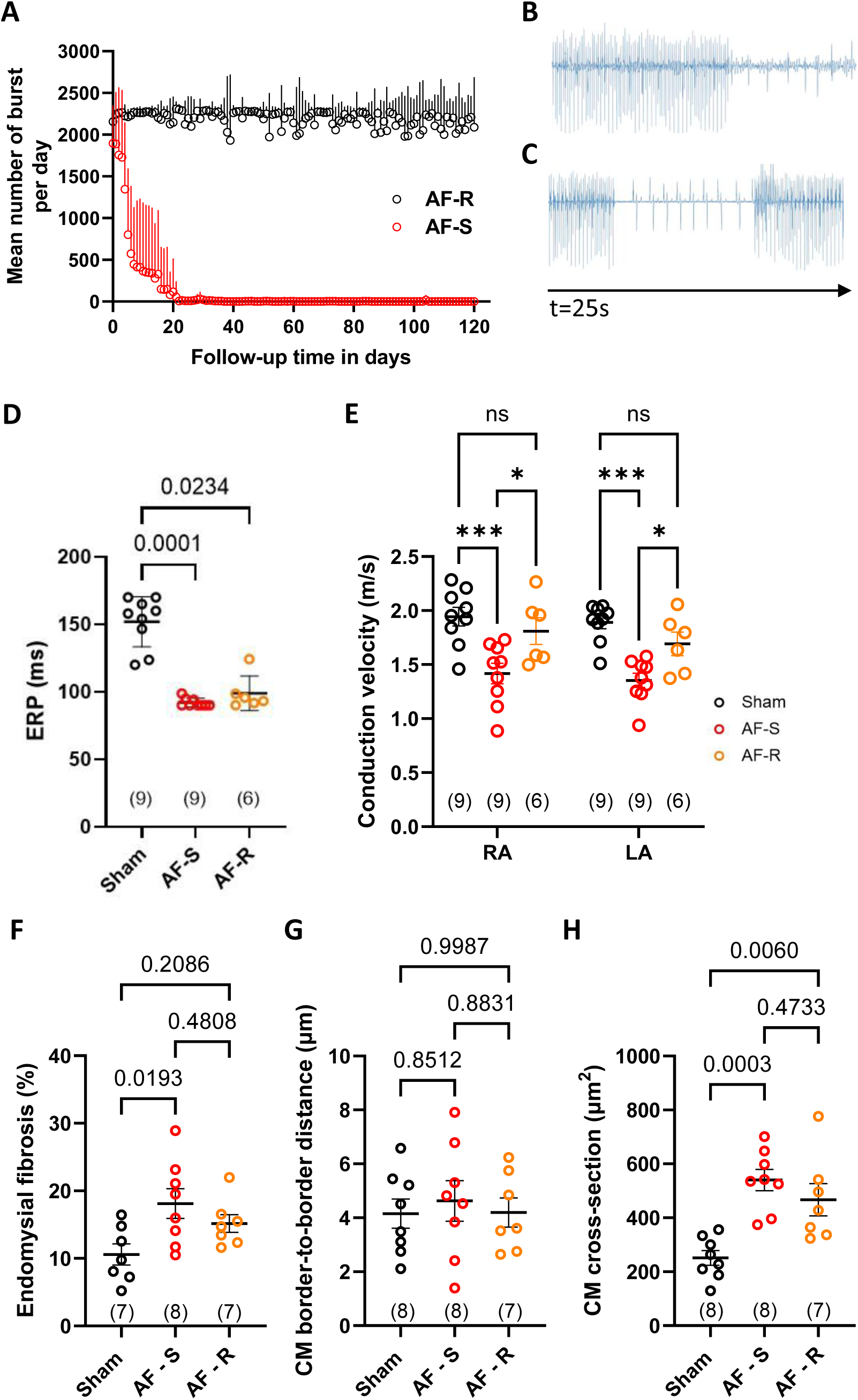
AF properties, electrophysiological and structural remodeling. **A**: Tachypacing bursts per 24-hours as a function of follow-up time. Results shown for AF-Sensitive (AF-S, N = 12) and AF-Resistant (AF-R, N = 8) sheep. **B-C**: Typical traces of the pseudo-ECG morphology recorded by the implanted telemetry. **B**: A burst of rapid atrial pacing followed by an episode of atrial fibrillation in the AF-S group. **C**: An episode of atrial fibrillation spontaneously converting to sinus rhythm observed in the AF-R group. Return to sinus rhythm triggered a burst of rapid atrial pacing. **D**: Effective refractory periods (ERPs) measured during electrophysiological contact mapping. **E**: Atrial conduction velocity calculated from the 3D maps obtained during contact mapping of the right (RA) and left (LA) atrium. **F**: Endomysial fibrosis measured from histological sections stained with Masson Trichrome. **G**: Cardiomyocyte (CM) border-to-border distance. **H**: CM cross-sectional area. Panels D to H show individual points and mean ± SD. Statistical analysis was performed with Kruskal-Wallis test and Dunn’s post-hoc test for multiple comparisons. Numbers in brackets correspond to individual experiments for each group. Complementary analysis is presented in Supplemental Figure 1.

Contact mapping and programmed stimulation, performed under isoflurane anesthesia (see Methods), indicated similar ERP reduction in both groups (Figure 1D and supplemental Figure 1E). Atrial CV calculated in the right and left atrium decreased significantly in AF-S when compared to both Sham and AF-R, which did not differ from each other (Figure 1E). Endomysial fibrosis was increased in AF-S versus Sham only (Figure 1F; supplemental Figure 1F). While extracellular space dimensions (Figure 1G) showed no differences between groups, AF-S and AF-R sheep both had increased cardiomyocyte cross-sectional area versus Sham (Figure 1H).

### Proteomic, enzymatic and metabolomic profiling

LAA proteomic profiling revealed significant differences between Sham, AF-S and AF-R sheep (Figure 2A). In general, AF-S versus Sham showed the greatest changes. Both AF-S and AF-R exhibited increased expression of proteins involved in ribosome assembly versus Sham, suggesting a common pattern of altered proteostasis. However, they also presented distinctive features: AF-R showed significant dysregulation of pathways involved in adrenergic signaling and cardiac contraction, while AF-S primarily exhibited modulation of metabolic pathways, including glycolysis/gluconeogenesis, fatty acid degradation, pyruvate and branched chain amino acid metabolism. Compared to AF-R, AF-S sheep showed consistent remodeling of the glycolytic pathway, with a notable overexpression of hexokinase-2 (HK2) and a trend towards an increase in proteins involved in glycogen synthesis (https://doi.org/10.6084/m9.figshare.28269872.v1). Based on these results we quantified LAA tissue hexokinase (Figure 2B) and L-lactate dehydrogenase (LDH, Supplemental Figure 2A) activities. We also quantified G6P (Figure 2C), pyruvate (Supplemental 2B), L-lactate (Supplemental Figure 2C), pyruvate to L-lactate ratio (Figure 2D), and glycogen (Figure 2E) content in LAA. These results show glycolysis activation in both AF-S and AF-R groups versus Sham, promoting G6P and associated glycogen accumulation in both groups. A schematic comparison of the glycolytic remodeling in the three groups is shown in Figure 2F. G6P is also linked to posttranslational modification (PTM) by O-GlcNAcylation, a PTM proposed to participate in AF-progression in the context of diabetes [31]. Assessment of protein O-GlcNAcylation in AF-S and AF-R (Supplemental Figure 3A) showed a net increase when compared to Sham, with no changes in the expression of the three enzymes regulating this pathway (Supplemental Figures 3B and 4). Glycolysis activation in AF-S and AF-R groups was not associated with increased activity of LDH (Supplemental Figure 2A) or accumulation of L-lactate (Supplemental Figure 2C), but was paralleled by both an accumulation of pyruvate (Supplemental Figure 2B) and an increased pyruvate to L-lactate ratio (Figure 2D) in AF-S LAA versus Sham. Finally, the AF-R group showed an increased expression of the monocarboxylate isoform 1 (MCT1) when compared to Sham (Supplemental Figure 2D).

**Figure 2.**
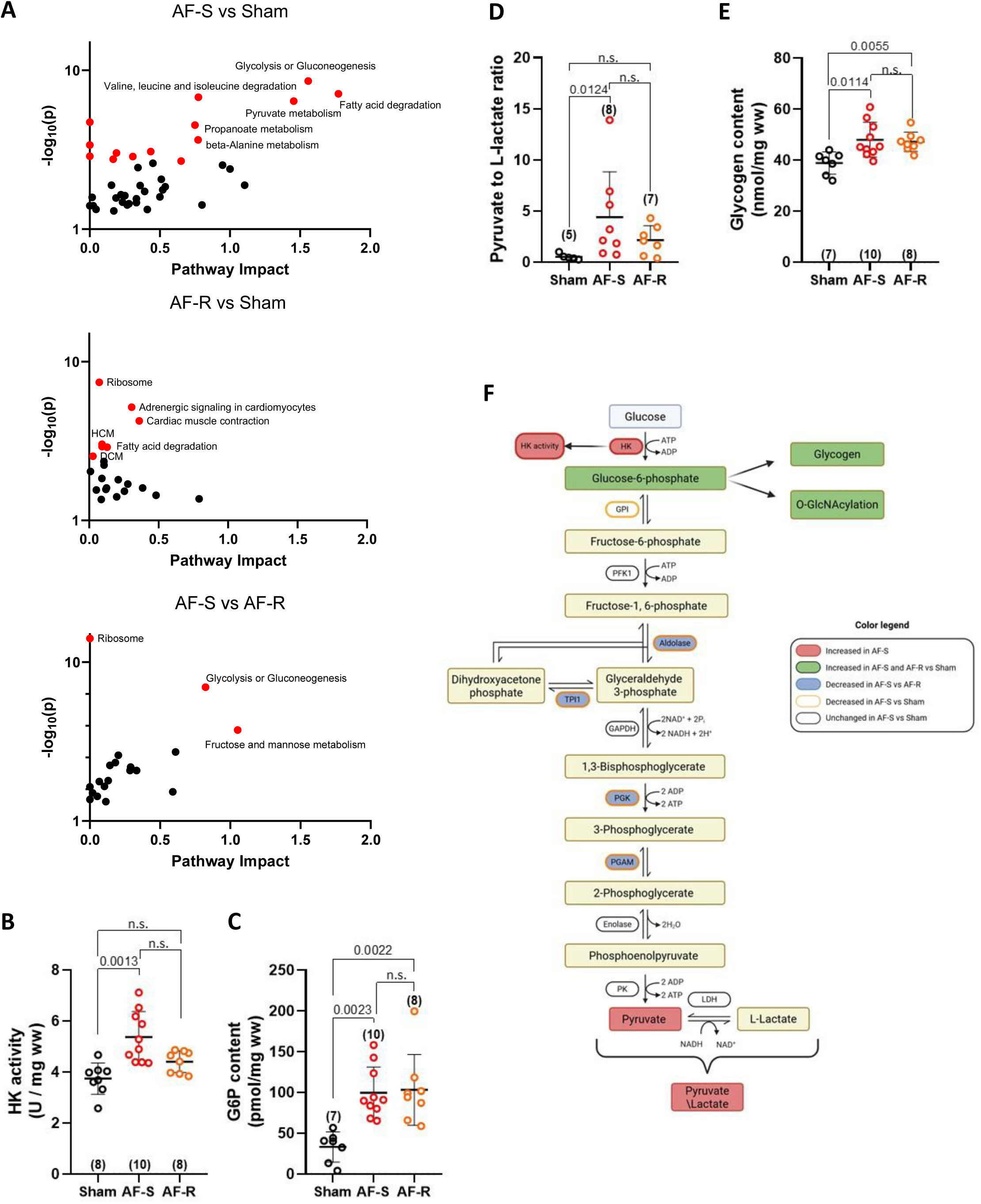
Proteomic, enzymatic and metabolomic analysis of LA-appendage. LAA samples freeze-clamped *in vivo* were analyzed by mass spectrometry for protein identification. The computational analysis suggested dysregulation of multiple proteins involved in mitochondrial function, metabolism and bioenergetics. **A**: Pathway enrichment analysis results for the comparisons between the 3 groups – red dots represent the pathways with false discovery rate <0.05. B to E: Hexokinase activity (HK, panel B), Glucose-6-phosphate (G6P, panel C), pyruvate to L-Lactate ratio (panel D), and Glycogen content (panel E) assessed in frozen LAA. Panel **F**: scheme summarizing the glycolytic remodeling assessed by proteomic, enzymatic and metabolomic studies. Complementary results are shown in supplemental figures 2, 3 and 4. B to E show individual points and mean ± SD. Statistical analysis was performed with Kruskal-Wallis test and Dunn’s post-hoc test for multiple comparisons. Numbers in brackets correspond to individual experiments for each group.

The proteomic profile AF-S LAA tissue was also characterized by a significant decrease in TCA cycle enzymes (FH, SUCLA2), cytosolic and mitochondrial adenylate kinases (AK1, AK2), mitochondrial malic enzyme (ME3), ROS detoxification (SOD2, PRDX3), cellular detoxification (ALDH3A2), and ATP synthase subunit-8 (MT-ATP8). AF-S LAA also showed increases in protein expression related to hypertrophic signaling (STAT3), acetyl- CoA production (ACSS1), glutathione synthesis (GCLC), outer membrane fusion (MFN1), complex I subunit (MT-ND4) and complex IV (COA1) assembly factor (Shown in raw data available at https://doi.org/10.6084/m9.figshare.28269872.v1). Given these findings, targeted metabolomic analysis for some TCA cycle metabolites and screening of the activity of TCA cycle enzyme were performed to further explore mitochondrial metabolic differences between the three groups. Compared to Sham and AF-S, AF-R showed a significant increase in isocitrate content (Figure 3A), while citrate and L-malate showed no differences between the three groups (Supplemental Figures 5A and B). Compared to Sham and AF-R, the AF-S group showed an accumulation of succinate, a substrate of respiratory chain complex II, while no differences in fumarate were observed between groups (Figures 3B and C). The TCA cycle metabolomic analysis suggests differential regulation of the TCA cycle between AF-R and AF-S, with potential impact on electron transport chain function through succinate accumulation, as well as redox modulation within the mitochondrial matrix. Regarding the TCA enzyme activity assessment, citrate synthase activity was evaluated at the tissue level and revealed increased activity in AF-R versus Sham, while AF-S sheep showed an intermediate phenotype (N.S. vs AF-R or Sham, Figure 3D). At the level of isolated mitochondria, the AF-S group was characterized by increased activity of pyruvate dehydrogenase (PDH, Figure 3E) and alpha-ketoglutarate dehydrogenase (𝛼-KGDH, Figure 3F), while fumarase activity was decreased (Figure 3G). Both AF-S and AF-R mitochondria showed decreased activity of L-malate dehydrogenase (MDH2, Figure 3H), and increased activity of 2-hydroxyglutarate dehydrogenase (Supplemental Figure 5C), an enzyme that produces alpha-ketoglutarate. In contrast, AF-R showed a decrease in succinyl-CoA synthase activity (SCS, Figure 3I) compared to both Sham and AF-S.

**Figure 3.**
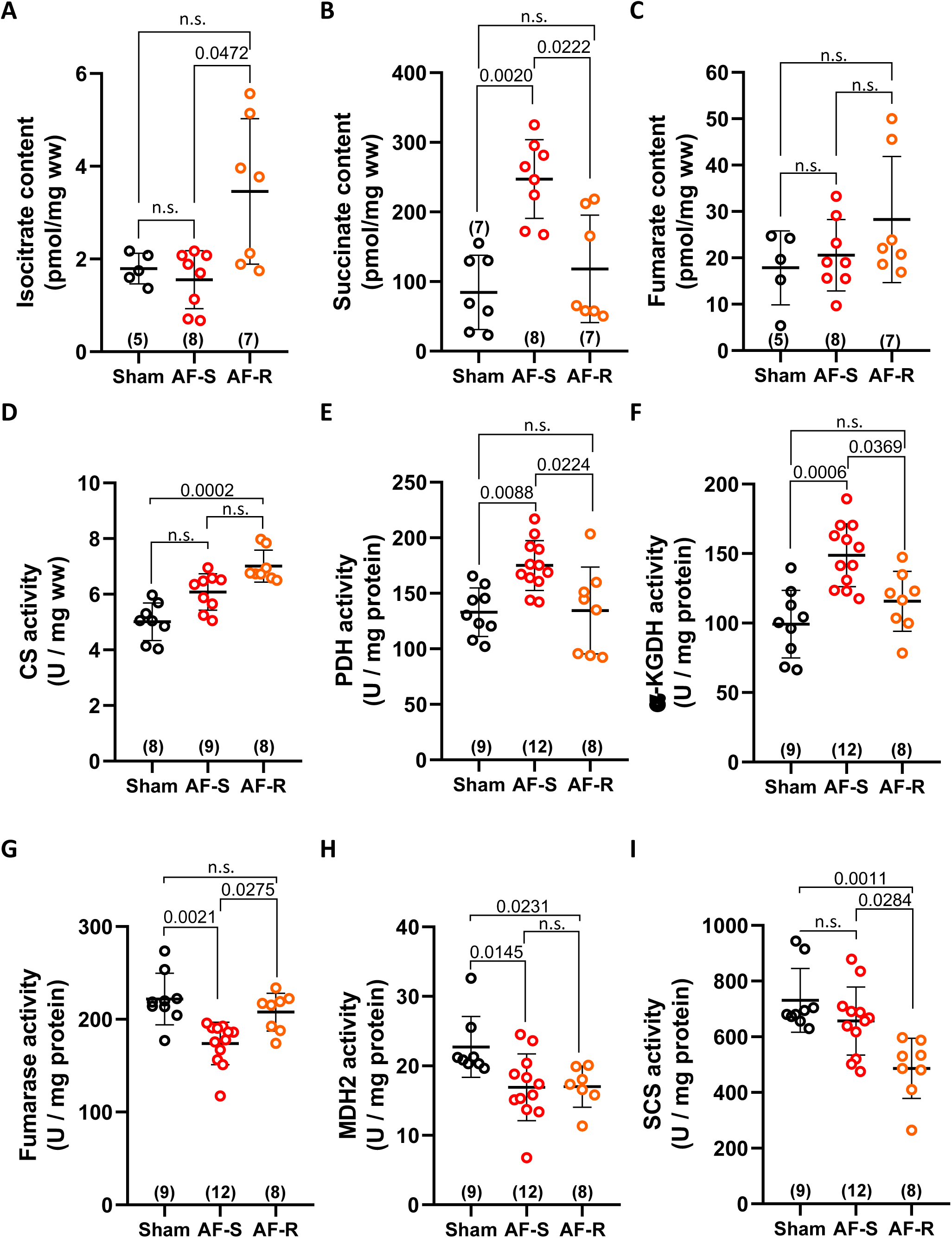
Metabolite content and enzyme activity in LA Appendage (LAA). Samples of LAA freeze-clamped *in vivo* were analyzed by high-pressure ion chromatography (HPIC) or enzymatic assay for metabolites belonging to the TCA cycle (see methods). Metabolite content is expressed in pmol/mg wet weight (ww). For TCA cycle, isocitrate (**A**), succinate (**B**), and fumarate (**C**) content were assessed. Enzymes’ activity was assessed on isolated mitochondria except for Citrate Synthase (CS, **D**) measured using LAA homogenate. Pyruvate dehydrogenase (PDH, **E**), alpha-ketoglutarate dehydrogenase (𝛼-KGDH, **F**), fumarase (**G**), L-malate dehydrogenase isoform 2 (MDH2, **H**), and succinyl-CoA synthetase (SCS, **I**). Panels show individual data points and mean ± SD. Statistical analysis was performed with Kruskal-Wallis test and Dunn’s post-hoc test for multiple comparisons. Numbers in brackets correspond to individual experiments. Complementary analysis is presented in Supplemental Figure 5.

By increasing pyruvate supply (secondary to increased glycolysis), acetyl-CoA production (increased PDH activity and ACSS1 content) and 𝛼-KG production (increased 𝛼-KGDH), while showing decreased activity of fumarase and MDH2, metabolic remodeling in AF-S sheep would promote succinate accumulation. In contrast, AF-R mitochondria showed a different remodeling pattern, with decreased activity of SCS and MDH2 despite increased CS activity potentially explaining isocitrate accumulation versus AF-S (NS versus Sham). Thus, the combined proteomic, metabolomic and enzymatic profile suggest a potential mitochondrial/metabolic contribution to AF resistance, prompting detailed investigations into mitochondrial function differences between AF-S and AF-R sheep.

### LAA mitochondrial function assessment

To gain insight into the metabolic behavior of mitochondria, we designed experiments to assess mitochondrial function using various respiratory substrates (state 2: non-phosphorylating respiration) and after maximal ATP synthesis induction (state 3: phosphorylating respiration), while mitochondria were oxidizing pyruvate + malate (PM), or glutamate + malate + succinate (GMS), or GMS along with the succinate-driven ROS emission inhibitor S1QEL1.1 [32] (S1QEL), or palmitoyl-carnitine + malate (PCM).

AF-S sheep exhibited quantitatively (but not significantly) greater state 2 respiration under PM conditions (Figure 4A), and lesser state 3 respiration (Figure 4B) associated with a significantly reduced state 3 to state 2 ratio compared to Sham in the presence of PM (Figure 4C). No differences were observed in other substrate combinations in state 2 or 3 respiration, nor between AF-R and Sham for any substrate condition (Figures 4A to C). Isolated mitochondria from AF-S sheep showed a decreased ATP-synthesis rate with PM or GMS compared to Sham and AF-R, but not with PCM (Figure 4D). The P to O ratio, representing the oxidative phosphorylation (OXPHOS) yield, was significantly decreased in AF-S mitochondria with GMS compared to Sham and AF-R (with a similar trend in PM), but remained unchanged with PCM between the three groups (Figure 4E). S1QEL decreased both state 3 respiration and ATP synthesis rate, and eliminated P to O differences observed in GMS among groups (Figure 4E). To assess whether the P to O ratio decrease in AF-S might influence the atrial energetic status, LAA content in PCr, ATP, ADP and AMP were quantified (Supplemental Figure 6). AF-S LAA, compared to AF-R and Sham, showed non- significant differences in ADP content and ATP to AMP ratio. In contrast, the ATP to ADP ratio was lower in AF-S compared to AF-R (Supplemental Figure 6E). AF-R LAA showed an increase in PCr content (*p*=0.05, Supplemental Figure 6D) compared to Sham.

**Figure 4.**
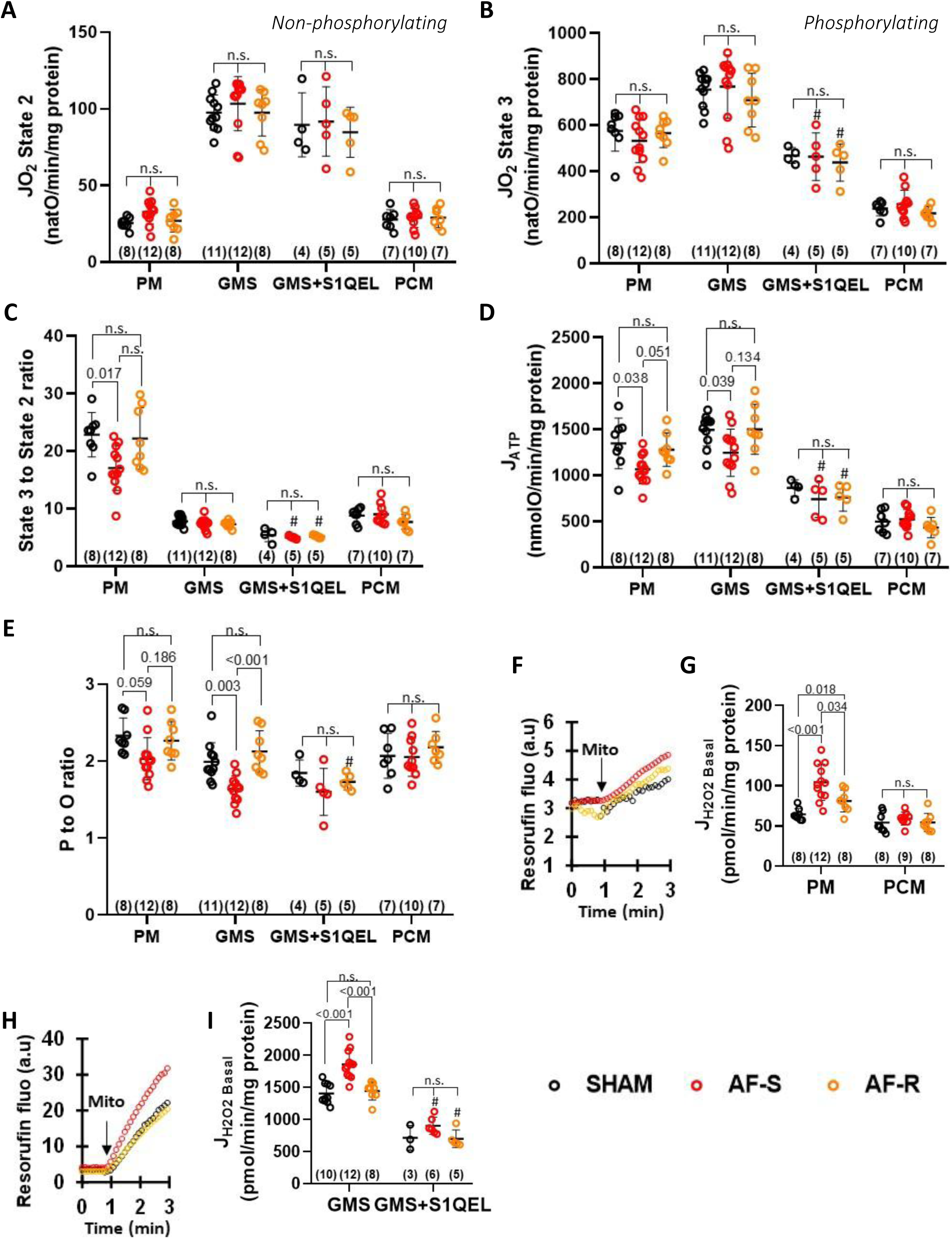
O_2_-consumption, respiratory coupling, rate of ATP synthesis, and H_2_O_2_ emission from mitochondria. Mitochondria were isolated and purified from LAA of Sham, AF-Sensitive (AF-S), and AF-Resistant (AF-R) sheep to evaluate (i) electron transport chain function, (panels **A:** State 2 non-phosphorylating respiration, **B:** respiration coupled to maximal ATP-synthesis (State 3, phosphorylating respiration) **C**: State 3 to State 2 ratio, **D**: rate of ATP synthesis, and **E:** P to O ratio reflecting OXPHOS yield. **F-I:** rate of hydrogen peroxide emission (H_2_O_2_ emission) assessed with pyruvate + malate or palmitoyl-carnitine + malate (PM or PCM, Panels **F** (raw traces) and **G**), or glutamate + malate + succinate in absence or presence of S1QEL (GMS or GMS+S1QEL, panels **H** (typical traces) and **I**). Panels show individual data points and mean ± SD. Statistical analysis was performed with t-test using the Holm-Sidak method to correct for multiple comparisons. The numbers in brackets represent individual experiments in each group. #*p*<0.05 GMS + S1QEL *vs.* GMS, by Wilcoxon test for paired samples. Complementary analysis is presented in Supplemental Figure 7.

Mitochondrial ROS emission was significantly increased during state 2 respiration under PM conditions in AF-S and AF-R sheep, compared to Sham (Figures 4F and G). This mitochondrial phenotype was not altered with PCM. In contrast, during succinate oxidation, only AF-S mitochondria showed increased ROS emission versus AF-R and Sham, which was prevented by S1QEL (Figures 4H and I). Mitochondrial ROS emission during state 3 respiration was identical among groups (Supplemental Figure 7A). Increased ROS emission in AF-S with succinate was associated with a more reduced NAD(P)H pool in state 2 respiration when compared to Sham and to both groups during state 3 respiration (Figures 5A and B).

**Figure 5.**
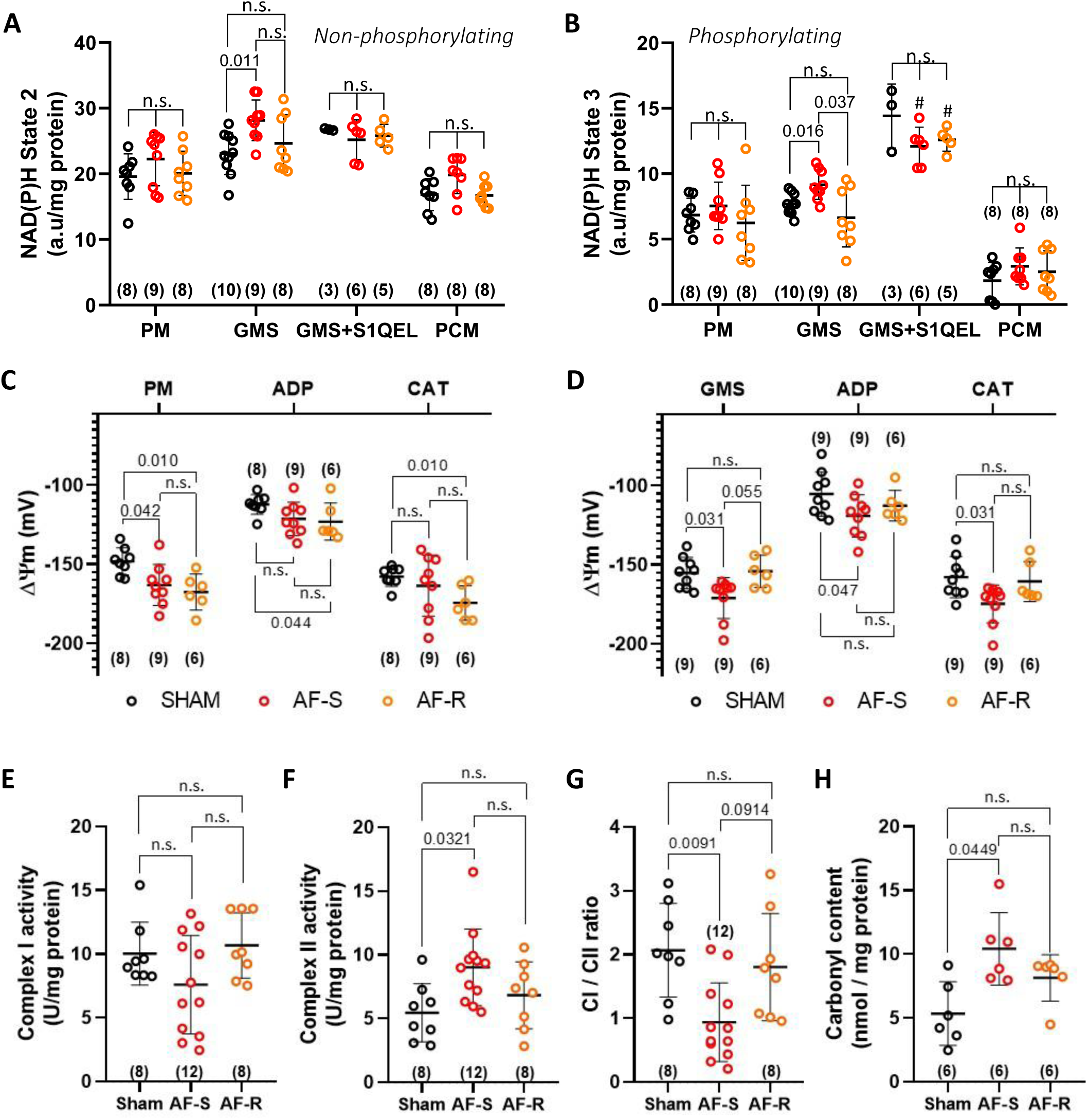
Mitochondrial membrane potential, complexes I and II specific activity in isolated mitochondria, carbonyl, NAD^+^ and NADH content in LAA. **A** and **B:** NAD(P)H autofluorescence measured on isolated mitochondria during state 2 (**A**) or state 3 (**B**) respiration while oxidizing different respiratory substrates: pyruvate + malate (PM), glutamate + malate + succinate (GMS), GMS + succinate-driven ROS inhibitor S1QEL (GMS+S1QEL), or palmitoyl-carnitine + malate (PCM). **C** and **D:** Mitochondrial inner membrane potential (ΔΨm) measured on isolated mitochondria oxidizing pyruvate + malate (**C**, PM) or glutamate + malate + succinate (**D**, GMS), and after sequential addition of 1.5 mmol/L ADP, and 5 µmol/L carboxyatractyloside (CAT) to stop ATP synthesis in Sham, AF-S, and AF-R groups. **E, F** and **G:** Mitochondrial complex I (**E**) and complex II (**F**) specific activity, and complex I/II activity ratio (**G**) in Sham, AF-S, and AF-R. **H:** Total carbonyl content measured by spectrophotometry. Panels show individual data-points and mean ± SD. Statistical analysis in panels A to D, was performed with t-tests using the Holm-Sidak method for multiple comparisons. Statistical analysis in panels E to H, was performed by Kruskal-Wallis test and Dunn’s post-hoc test for multiple comparisons. The numbers in brackets represent individual experiments.

An increase in inner mitochondrial membrane polarization (ΔΨm) was observed in AF-S and AF-R compared to Sham under state 2 respiration in PM conditions (Figure 5C, hyperpolarization of 14 and 20 mV for AF-S and AF-R versus Sham respectively), which was only significant during ATP synthesis and after its inhibition by CAT in AF-R mitochondria. Interestingly, under GMS conditions, AF-S mitochondria showed an apparent hyperpolarization compared to both Sham and AF-R (Figure 5D, 16 mV versus Sham in state 2). The inner membrane hyperpolarization in AF-S sheep was significant versus Sham during state 3 respiration (ADP, 14 mV versus Sham) and after its inhibition by CAT (17 mV vs Sham). No differences among groups were observed with PCM (Supplemental Figure 7B). It should be noted that mitochondrial potential was calculated with a fixed matrix volume for all groups and experimental conditions. Nonetheless, MFN1 expression was strongly increased in AF-S compared to Sham and AF-R (Supplemental Figure 7C), suggesting mitochondrial fragmentation and associated decreased matrix volume. Although ΔΨm hyperpolarizes when matrix volume decrease (Supplemental Figure 7D) the level of mitochondrial fragmentation could not explain on its own the 15 mV hyperpolarization observed in the AF-S group.

Since ΔΨm is the main driving force allowing calcium accumulation within the matrix, we measured the mitochondrial calcium uptake rate and maximal quantity of calcium needed to open the mitochondrial permeability transition pore (mPTP) in the presence of succinate. AF- S mitochondria exhibited a significant increase in calcium uptake compared to both Sham and AF-R (Supplemental Figures 7E and F). Compared to Sham, mPTP sensitivity to calcium was increased in AF-R, and AF-S showed an intermediate phenotype (NS vs Sham or AF-R Supplemental Figure 7G).

Thus, in-depth mitochondrial function analysis revealed common characteristics between AF- S and AF-R during PM oxidation, while several mitochondrial functions were only altered in AF-S during succinate oxidation.

Activity of electron transport chain complexes I and II was then investigated. No differences were observed at the level of complex I (Figure 5E), while complex II activity was significantly increased in AF-S versus Sham (Figure 5F, N.S. for AF-R versus Sham).

Complex II competes with complex I for ubiquinone reduction, and as a consequence, can modulate the level of NADH oxidation and ROS production. The complex I to complex II activity ratio decreased in AF-S compared to Sham (Figure 5G, AF-R N.S. vs Sham), consistent with a contribution of succinate to ROS emission (Figure 4I).

### LAA energetic/redox status

The overmentioned results suggesting ROS alterations in AF-S led us to evaluate the extent of oxidatively-modified protein accumulation in the atrial compartment, as indicated by carbonyl content. Carbonyl content increased significantly in AF-S versus Sham, with no significant difference between AF-R and Sham (Figure 5H).

To further understand the differential redox regulation within the atria of AF-S and AF-R sheep, NAD^+^ and NADH (cytosolic + mitochondrial) content were quantified. Compared to Sham, AF-S and AF-R mitochondria showed increased NAD^+^ total content (Figure 6A), but AF-S LAA had significantly decreased NADH total content compared to both Sham and AF- R (Figure 6B). Thus, the NAD^+^ to NADH ratio was significantly increased in AF-S LAA versus both AF-R and Sham, indicating a redox disturbance not observed in AF-R LAA (Figure 6C). Moreover, using the measured pyruvate to L-lactate ratio (Figure 2D) and the L-lactate dehydrogenase equilibrium constant (Supplemental Material), the cytosolic [NAD^+^] to [NADH] ratio was calculated and showed a more oxidized cytosolic compartment in both AF- S and AF-R compared to Sham LAA (Figure 6D).

**Figure 6.**
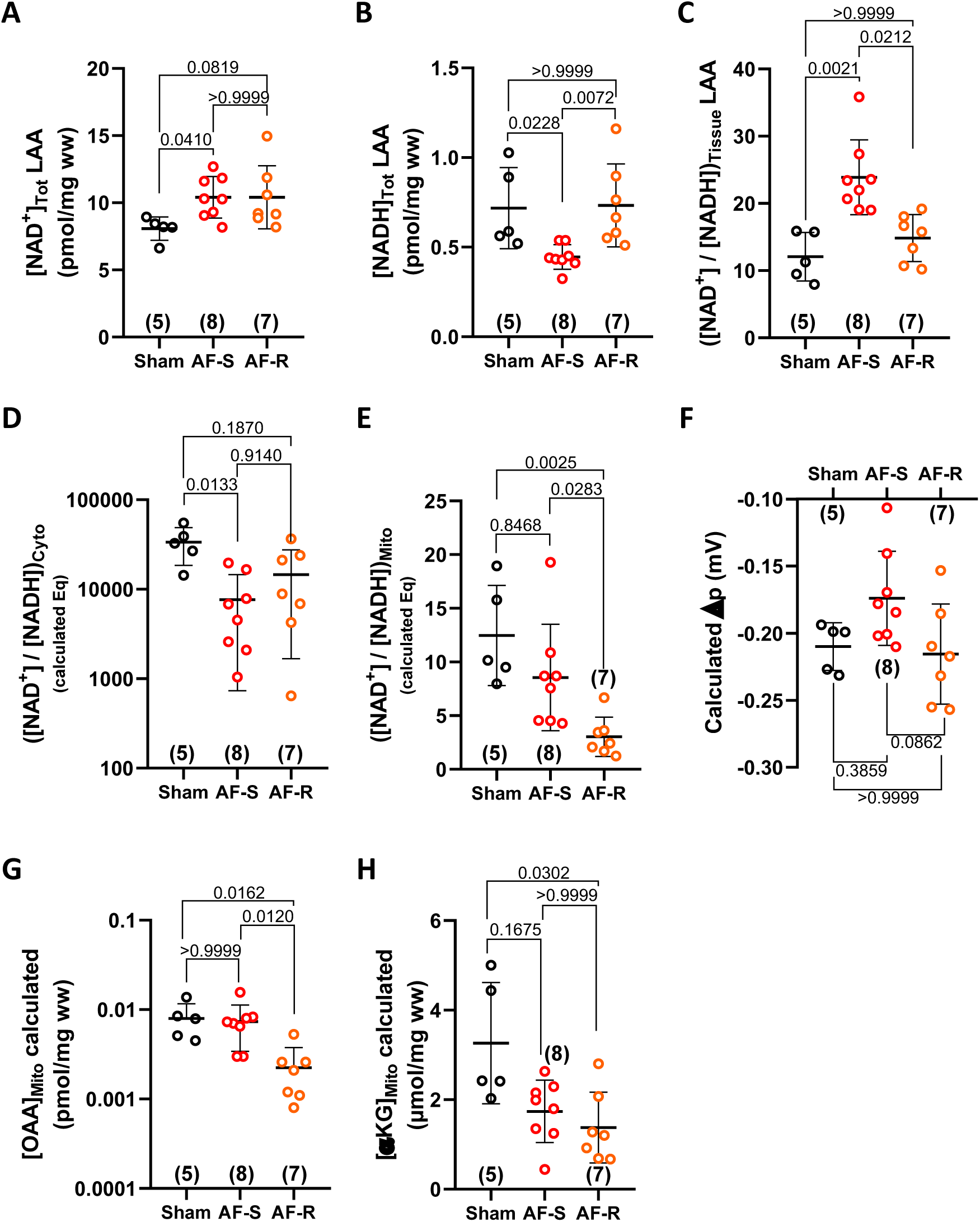
Differential mitochondrial redox regulation within AF-R and AF-S LAA. A, **B** and **C:** LAA content (HPIC approach see supplemental Material) in NAD^+^ (**A**) and NADH (**B**), and corresponding NAD^+^ to NADH ratio (**C**). **D** and **E:** cytosolic (**D**) and mitochondrial (**E**) NAD^+^ to NADH ratio calculated using the L-lactate to pyruvate and β-hydroxybutyrate to acetoacetate ratio under thermodynamic equilibrium postulate for the reaction catalyzed by lactate dehydrogenase and β-hydroxybutyrate dehydrogenase. **F:** calculated mitochondrial proton motive force (Δp) within the LAA tissue while considering the system at thermodynamic equilibrium. **G** and **H:** calculated mitochondrial oxaloacetate (**G**, [OAA]) and 𝛼-ketoglutarate (**H**, [𝛼-KG]) concentrations. Panels show individual data-points and mean ± SD. Statistical analysis was performed by Kruskal-Wallis test and Dunn’s post-hoc test for multiple comparisons. The numbers in brackets represent individual experiments.

The same approach was used to calculate the [NAD^+^] to [NADH] ratio within the mitochondrial compartment of the three groups. For this purpose, the β-hydroxybutyrate to acetoacetate ratio was quantified in LAA (Supplemental Figures 8A to C). AF-R LAA showed an increased matrix ratio of these ketone bodies (Supplemental Figure 8C) compared to both Sham and AF-S. As a consequence, the calculated mitochondrial [NAD^+^] to [NADH] ratio is lower (reduced environment) in AF-R versus both Sham and AF-S (Figure 6E).

Furthermore, from these local redox ratios, the overall tissue NAD^+^ to NADH ratio at equilibrium was calculated considering the NAD(H)^+^ content in the cytosolic and mitochondrial compartments (N_ratio_) as well as the ratio of the two compartments volumes (V_ratio_). Plotting the measured [NAD^+^] to [NADH] ratio as a function of the calculated value (Supplemental Figure 8D), showed that in Sham LAA there was no significant difference (Wilcoxon rank sum test) between the measured and calculated ratio. Thus, the redox system of Sham LAA works close to equilibrium. In contrast, the AF-S and AF-R groups are clearly far from the equilibrium hypothesis (*p*=0.0078 for both) and this should be considered as a potential secondary effect to the metabolic remodeling.

Owing to the fact that the calculated overall redox ratio of the tissue is mainly fixed by the mitochondrial NAD^+^ to NADH ratio (Supplemental Figure 8E), one can argue that β-HBDH may be working far from thermondynamic equilibrium in AF-R LAA and to some extent, in AF-S LAA, compared to Sham. Thus, it is likely that the decreased [NAD^+^] to [NADH] ratio measured within LAA tissue reflects oxidation of the mitochondrial compartment in AF-S compared to Sham and AF-R.

To further attempt to evaluate mitochondrial function within LAA tissue, the mitochondrial proton motive force (Δp) was also calculated at equilibrium for Sham, AF-S and AF-R LAA (Figure 6F). The results show that mitochondria showed a lower Δp in AF-S compared to Sham and AF-R. Thus, AF-S mitochondria *in vivo* could have a higher rate of oxygen consumption compared to Sham and AF-R, which is in line with the previously observed NAD^+^ oxidation state. This approach was used to calculate the relative [oxaloacetate] ([OAA]), [alpha-ketoglutarate] ([𝛼-KG]) and [L-malate] within the cytosolic and mitochondrial compartments (Figures 6G and H) and Supplemental Figures 8F to H). AF-R mitochondria showed a lower [OAA]_mito_ (Figure 6G) and [𝛼-KG]_mito_ (Figure 6H) than Sham, further suggesting that AF-R LAA have a differential remodeling of the TCA cycle versus Sham and AF-S.

### Succinate and atrial arrhythmia susceptibility in a rodent model

Our results suggest that in our ovine model, AF stabilization is paralleled by TCA cycle remodeling, mitochondrial succinate accumulation and oxidation of the mitochondrial compartment. Based on the data in figures 3B, 4E, 4I, 5A, 5D, 5H, and 6C, it is possible that succinate accumulation participates in metabolic disturbances that stabilize AF in AF-S sheep. To assess the potential role of succinate accumulation in AF, we first evaluated wether acute exposure of atrial cardiomyocytes to exogenous succinate can alter atrial metabolism, ROS emission and NAD(P)H fluorescence *ex vivo* in beating rat hearts and *in vitro* in the isolated rat atrial cardiomyocytes (RACMs). Succinate is a dicarboxylate, known (at least in ventricular cardiomyocytes or skeletal muscle) to be transported across the membrane only in acidic conditions (i.e. ischemia or intense exercise) [33,34]. RACMs were incubated with a high concentration of succinate (5 mmol/L) and its impact on TCA cycle metabolites level assessed by mass spectrometry. Following succinate incubation, the intracellular content of different mitochondrial and cytosolic metabolites was assessed by mass spectrometry using deuterium-labelled succinate (see Methods) and internal oxalate content (Figure 7A) used for signal correction and normalization. This analysis showed that the intracellular concentrations of succinate, fumarate and L-malate were increased with succinate exposure, while citrate and alpha-ketoglutarate showed a trend towards increase (Figures 7B to D and Supplemental Figures 9A and B). Three glycolytic intermediates (glucose-6-phosphate, fructose-1,6- bisphosphate and pyruvate) showed no changes following RACM incubation with exogenous succinate (Supplemental Figures 9C to E). Incubation of RACMs with exogenous succinate significantly increased the intracellular succinate concentration by 3.8 fold (Supplemental Figure 9F). At the functional level, RACM incubation with succinate significantly increased ROS emission (Figures 7E and F, and supplemental Figure 9G), and succinate perfusion of Langendorff-perfused beating rat hearts significantly increased both NAD(P)H and MitoPY1 LAA fluorescence (Figures 7G to J). H_2_O_2_ perfusion showed no impact on NAD(P)H fluorescence and produced a net increase in MitoPY1 fluorescence (Supplemental Figures 9H to K).

**Figure 7.**
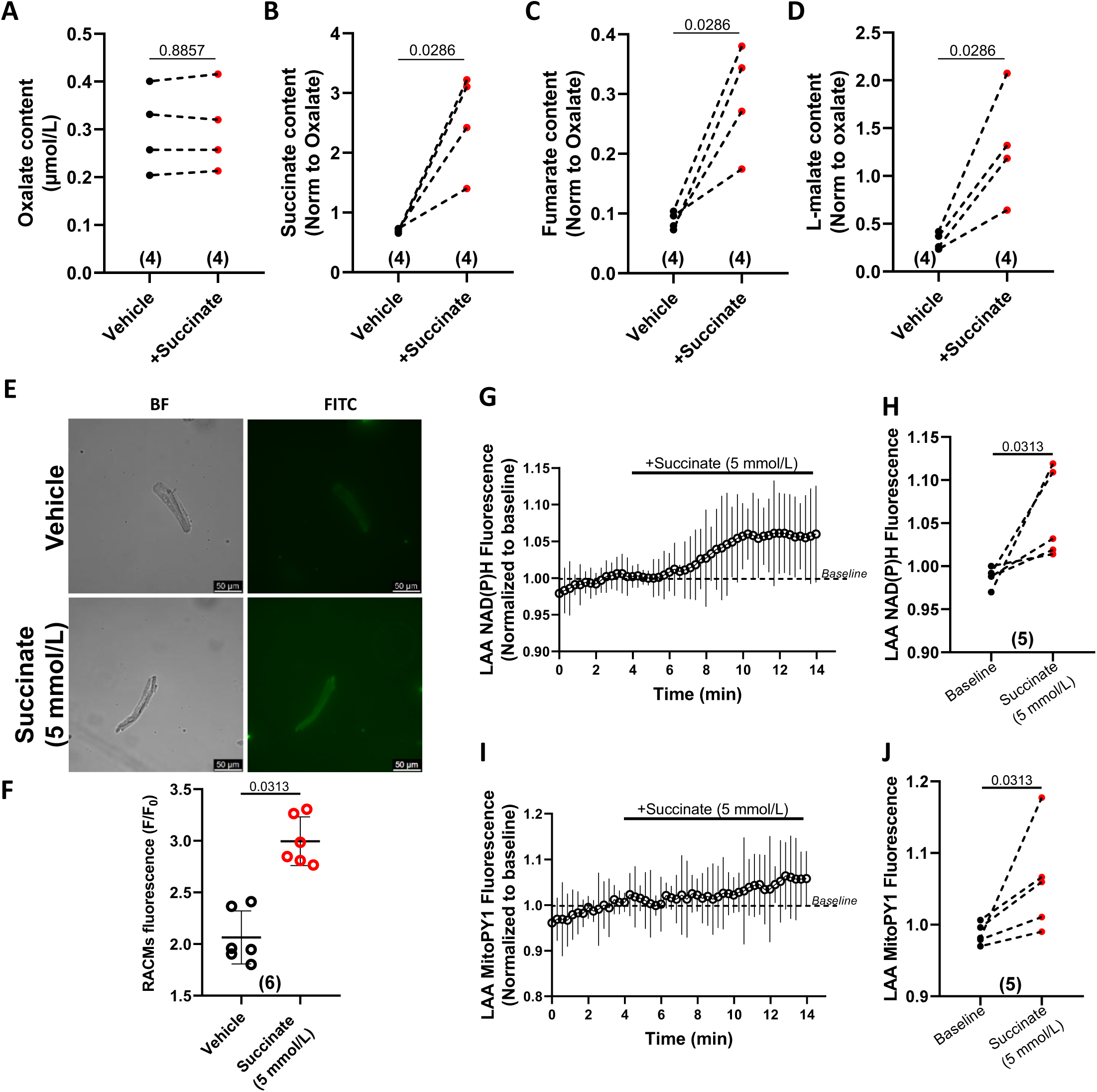
Effect of exogenous succinate application on atrial cardiomyocytes. **A-D:** Isolated rat atrial cardiomyocytes (RACMs) were incubated with 5 mmol/L of succinate and intracellular metabolites content was assessed by mass spectrometry. Signals correction-quantification and normalization were performed using both deuterium-labelled succinic acid and oxalic acid (**A**), respectively (see methods). Intracellular content of succinate (**B**), fumarate (**C**) and L-malate (**D**) is represented. **E:** Representative images illustrating ROS emission measurement by CellROX Green in RACM treated with either succinate at 5 mmol/L or vehicle (BF: bright field, FITC: Fluorescein). **F:** Quantification of intracellular ROS emission in presence of succinate measured with CellROX, with respect to the background. **G-J:** Evolution of LAA NAD(P)H (**G** (time course), **H** (quantification)) and MitoPY1 (H_2_O_2_ sensitive fluorescent probe, **I** (time course), **J** (quantification)) fluorescence measured in the *ex vivo* perfused rat heart, using an optical fiber positioned close to the LAA, upon perfusion with 5 mmol/L succinate. **A-D:** Statistical analysis *via* Mann-Whitney, N=4 in each group Test. **F, H and I:** Statistical analysis *via* Wilcoxon test for paired samples, N=6 panel F and N=5 panels G to J. Complementary analysis is presented in Supplemental Figure 10.

*In vivo* intravenous succinate administration increased atrial tachyarrhythmia vulnerability, susceptibility, and duration compared to values in both vehicle-infused and S1QEL pretreated animals (Figures 8A to C). Succinate-injected animals showed a significant increase in carbonyl content (Figure 8D, and supplemental Figures 10A and B). Succinate+S1QEL- treated rats had carbonyl content not significantly different from vehicle, but the reduction versus succinate-only was not statistically significant (Figure 8D).

**Figure 8.**
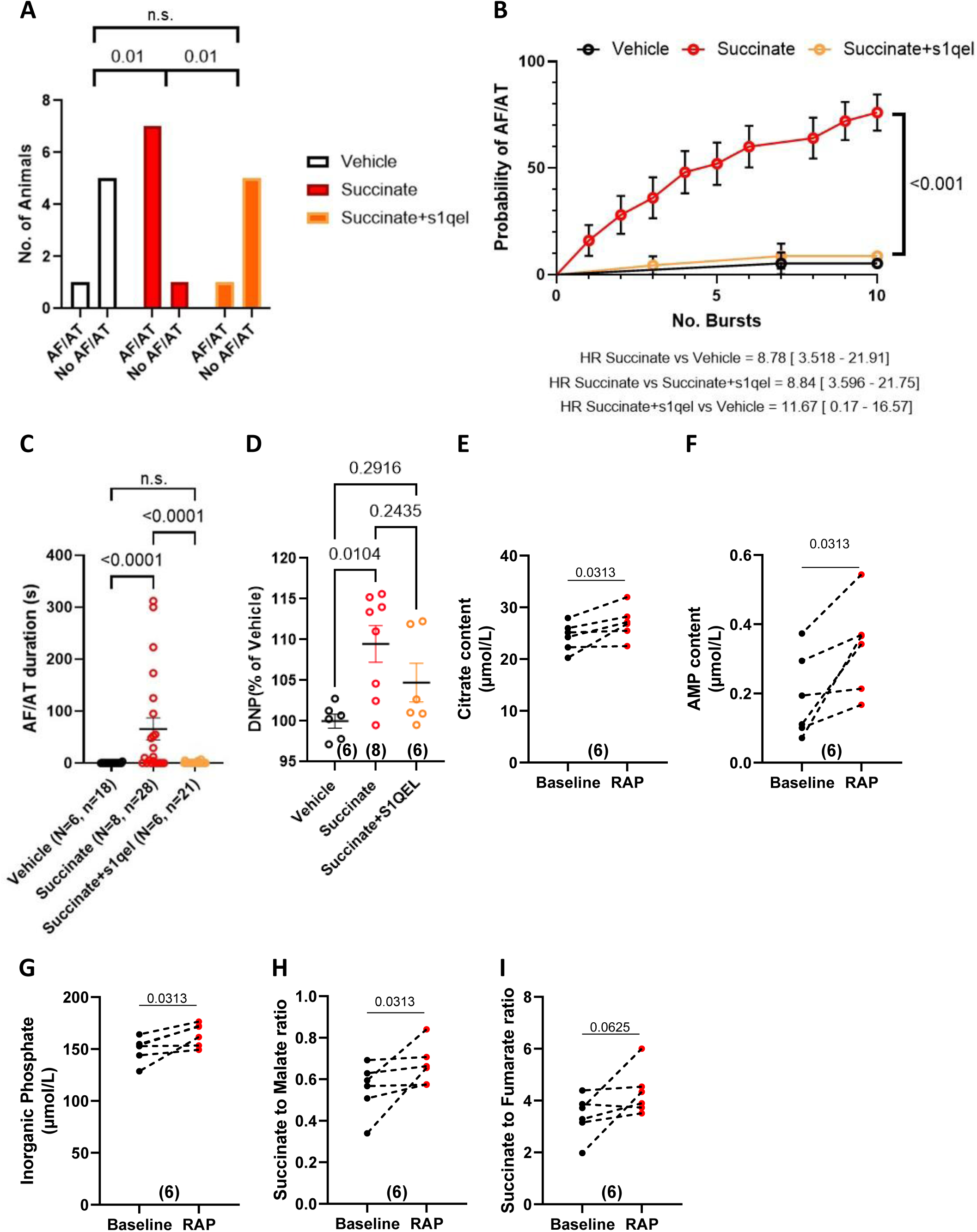
Effect of succinate on arrhythmia susceptibility and carbonylation *in vivo* and effect of RAP on blood metabolites. Atrial arrhythmia susceptibility was assessed *in vivo* by transesophageal burst-pacing following intravenous injection of vehicle, succinate or succinate + S1QEL. **A:** Number of animals with inducible AF/AT versus no arrhythmia. **B:** Cumulative probability of AF/AT as function of number of induction-bursts. **C:** AF/AT duration in response to a burst-pacing challenge. **D:** Carbonyl content of *in vivo* hearts measured by Oxyblot. **E-I:** Blood was harvested before (baseline) or after *in vivo* atrial tachystimulation by transesophageal burst-pacing (RAP), and citrate (**E**), adenosine monophosphate (AMP, **F**), inorganic phosphate (**G**) were assessed by mass spectrometry (N=6). **H**: Succinate to L-malate ratio. **I**: Succinate to fumarate ratio. Statistical analysis by Fischer exact test (**A**), Log-rank Mantel-Cox test (**B**), Kruskal-Wallis test and Dunn’s post-hoc test for multiple comparisons (**C, D**), and Wilcoxon test (**E**-**I**). Complementary analysis is presented in supplementary Figure 11.

In order to further explore the blood metabolic changes caused by atrial tachypacing *in vivo*, serum from rats subjected to RAP was analyzed by mass spectrometry. Acute atrial tachyarrhythmia was associated with the development of a specific blood metabolic profile compared to paired baseline control. RAP increased citrate, AMP, and inorganic phosphate, and caused a trend towards cis-aconitate increase, reflecting mitochondrial metabolic activation and increased energy consumption (Figures 8E to G, and Supplemental Figure 10C). A significant increase of the succinate to L-malate ratio and a trend towards increased succinate to fumarate ratio were observed (Figures 8H and I), although succinate, fumarate and L-malate were not changed (Supplemental Figures 10D to F).

## Discussion

### Major findings and novelty

We investigated the mechanisms underlying AF sensitivity and resistance in a sheep model, revealing involvement of metabolic, bioenergetic, redox and mitochondrial function differences. Notable findings in AF-S animals, compared to AF-R or Sham, included: i) altered proteomic, enzymatic and metabolomic profiles involving glycolysis, fatty acids utilization, and TCA cycle function/regulation; ii) increased succinate content and decreased ATP/ADP ratio, iii) increased activity of PDH and 𝛼-KGDH, paralleled by decreased fumarase activity likely participating to succinate tissue accumulation, iv) increased succinate-driven ROS emission; v) decreased OXPHOS efficiency during succinate oxidation, vi) altered equilibrium of the CI/CII activity ratio, and vii) increased mitochondrial [NAD^+^] / [NADH] ratio reflecting a redox crisis. Using the rat model, we show that atrial cardiomyocytes can metabolized exogenous succinate, and that this is paralleled by a redox balance alteration and an increased probability/stability of tachypacing-induced atrial arrhythmias. Moreover, we show that atrial tachypacing alone in the rodent leads to the development of a specific blood metabolic signature including an increased succinate to L-malate ratio. Finally, our analysis combining metabolite content assessment on *in vivo* freeze-clamped tissues with metabolic analysis by placing the system at the thermodynamic equilibrium suggests that resistant animal have more reduced mitochondrial matrix than AF-S sheep. This profile would be adaptative to the massive energetic burden caused by RAP, in terms of both OXPHOS yield and ROS detoxification efficiency when compared to the profile in AF-S sheep. Firstly, increased matrix NADH would promote electron fueling to complex I (push regulation), potentially improving the energetic status of AF-R atria compared to AF-S. Moreover, AF-R animals showed improved ketone body metabolism reflected by the increased β-OHB to AcAc (acetoacetate) ratio. Increased expression of BDH1 and OXCT1 (see https://doi.org/10.6084/m9.figshare.28269872.v1) would fuel acetyl-CoA towards the TAC cycle, further increasing NADH production. Secondly, a higher mitochondrial NADH level should favor ROS detoxification, protecting mitochondrial function. In contrast, the more oxidized matrix of AF-S mitochondria would limit NADH delivery to complex I, thus favoring entry of electrons at the level of complex II, especially in the presence of an accumulation of succinate, further decreasing the OXPHOS yield.

This is the first study, to our knowledge, to explore and identify metabolic factors differentiating AF-R animals from AF-S in a clinically relevant large animal model, highlighting the critical role of TCA cycle regulation and mitochondrial redox state in contributing to AF resistance. A schematic summary of our findings is provided in Figure 9.

**Figure 9.**
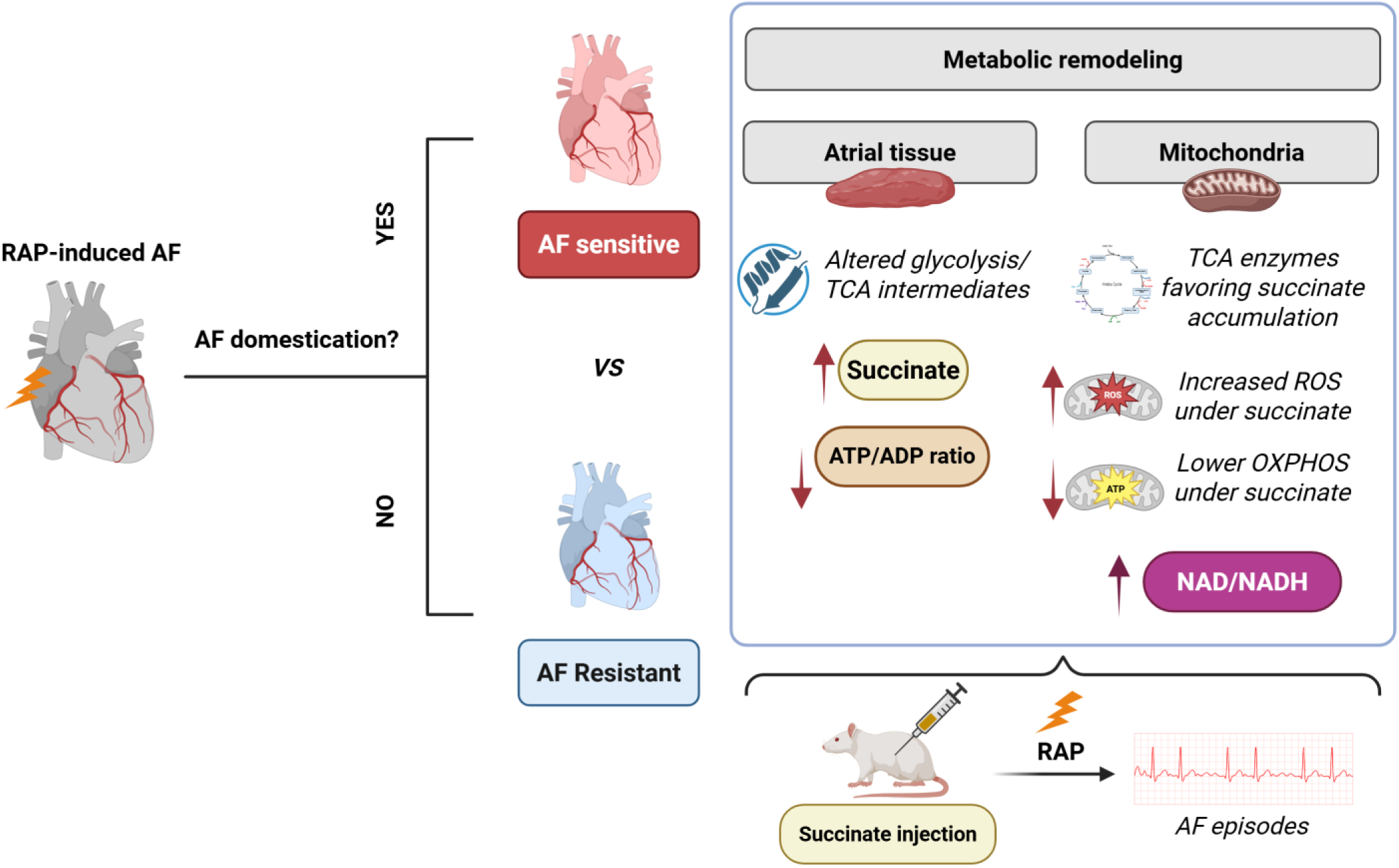
Scheme summarizing the study findings. Electrically-maintained AF led to domestication in AF-sensitive (AF-S) animals within 3 weeks, but some were AF resistant (AF-R) and failed to develop self-sustained AF despite 4 months of electrically maintained AF. Specific mitochondrial and metabolic remodeling were apparent only in AF-S. In presence of succinate, AF-S mitochondria showed increased ROS emission, decreased P to O ratio, increased complex II to complex I activity ratio. AF-R and AF-S mitochondria showed a differential regulation of the TCA cycle. At the whole-tissue level, AF-S LAA showed more oxidized proteins compared to Sham, AF-S alone showed an increased NAD^+^ to NADH ratio and an energetic impairment reflected by a decreased ATP to ADP ratio. AF-R LAA showed a higher PCr paralleled by an increased CKMT2 expression and a more reduced mitochondrial matrix secondary to an increased ketone body metabolism compared to AF-S. AF-S showed a differential TCA cycle enzymes regulation compared to AF-R, likely participating to succinate in the AF-S group. Created with BioRender.com.

### Involvement of LA energetic status and substrate metabolism in AF resistance

Increased energy demand caused by AF has different consequences in AF-S and AF-R sheep. AF-R sheep showed (versus AF-S) greater ATP to ADP ratio (Supplemental Figure 6E), increased mitochondrial content (Figure 3D) and larger PCr content than Sham (Supplemental Figure 6D), with a trend towards increased mitochondrial CK content (CKMT2 https://doi.org/10.6084/m9.figshare.28269872.v1 and supplemental Figure 8I), suggesting increased energy availability and indicating that stress-induced mitochondrial biogenesis adaptations contribute to AF-resistance. AF-S LAA showed specific mitochondrial function disturbances not observed in AF-R, particularly during succinate oxidation (Figures 4 and 5). Atrial energy metabolism is compromised by AF in patients and experimental models [8,16,35–38], characterized by decreased expression of assembling factors or core proteins of complexes I and II, reduced activity of these complexes [8,9,39], oxidative damage, and increased mitochondrial ROS production [8]. A recent study suggests that increased cardiomyocyte mitochondrial content and metabolic function are linked to AF-stabilization [40]. Prior studies also implicated mitochondrial dysfunction and energetic deficiencies in AF [36,41–44]. Our study goes beyond this previous work by evaluating metabolic changes associated with AF-resistance in an in vivo model and by identifying a potential key role of succinate accumulation in AF-sensitivity.

Reduced mitochondrial OXPHOS yield in AF-S mitochondria was observed with succinate only, but not with either fatty acid substrates (like palmitoyl-carnitine) or complex I linked respiratory substrate (Figure 4E, although a clear trend was observed). In our study, the shift towards glucose utilization previously reported in AF [45] was manifested in AF-S through increased (i) HK activity, (ii) pyruvate content, (iii) pyruvate to L-lactate ratio, and (iv) a lower cytosolic NAD^+^ to NADH ratio reflecting glycolysis activation. Moreover, both AF-S and AF-R groups showed increased atrial G6P and glycogen content, while L-lactate content or LDH activity were unchanged. Note that compared to Sham and AF-S, LAA AF-R showed a higher relative expression of MCT1 responsible for pyruvate and L-lactate exchange with the interstitial medium.

### Involvement of succinate buildup and TCA cycle remodeling in AF domestication

AF-S animals showed LA succinate accumulation likely secondary to TCA cycle remodeling leading to (i) a two step activation before complex II (PDH and 𝛼-KGDH) and (ii) a two step inhibition after complex II (Fumarase and MDH2). AF-S isolated mitochondria showed altered energetic output and increased ROS emission during succinate oxidation. Importantly, to get insights into the mitochondrial energy metabolism within the tissue, mitochondrial NAD^+^ to NADH ratio as well as Δp were calculated with the postulate of a system working close to the thermodynamic equilibrium, *i.e.* metabolic fluxes are rather small compared to enzyme capacities. As shown in supplemental Figure 8D, this hypothesis fully describes the physiological situation in the Sham group, while the AF-R group is more displaced from the equilibrium hypothesis. Considering this analysis and the evolution of Δp in the LA, we hypothesize that AF-S mitochondria in the LA are more depolarized than Sham or AF-R mitochondria. This is important when considering the impact of succinate accumulation on ROS emission because the lower the Δp the lower the succinate-driven ROS emission. Our results regarding carbonylation and the redox state of AF-S LA tissue indicate that the system is strongly oxidized in AF-S sheep. Thus, oxidative damage in AF-S LA tissue might be a consequence of both succinate buildup and failure of the detoxification system secondary to an alteration of the redox potential in the matrix compartment.

To our knowledge, this is the first report of a specific atrial accumulation of succinate and linked mitochondrial redox alteration in a relevant animal model. Succinate serves as a substrate for complex II in the respiratory chain and operates as a signaling molecule through the SUCNR1 receptor (formerly GPR91) [46,47]. Its accumulation has been implicated in various cardiovascular diseases [47–49], but its role remains relatively unexplored in AF. Elevated plasma succinate levels were associated with atrial dysfunction, stroke recurrence [48], and AF [50]. Succinate also accumulates in the ischemic heart [51], and AF has been suggested to be associated with supply-demand imbalance ischemia [52]. Examining the expression of both HIF1𝛼 and MCT4 (downstream to HIF1𝛼) by proteomic analysis we did not observe sign of hypoxia (https://doi.org/10.6084/m9.figshare.28269872.v1). In the AF atria, succinate accumulation is therefore not secondary to hypoxia in the sheep RAP model. Our detailed investigation of mitochondrial enzyme activity leads us to propose the following mechanisms contributing to succinate accumulation: AF-S mitochondria showed an increased activity of (i) HK, (ii) PDH, (iii) 𝛼-KGDH and (iv) 2-HGDH favoring pyruvate, acetyl-CoA and 𝛼-KG production which would favor succinate production at the level of the succinyl CoA synthetase (SCS). In parallel, fumarase and MDH2 activity were decreased, favoring succinate accumulation. In contrast, AF-R mitochondria showed a decrease in SCS and as well a lower level of 𝛼-KG, which might prevent succinate accumulation. Moreover, acute atrial tachypacing promoted both an increased succinate to L-malate ratio and citrate content in rat blood. AF-risk development in patients has been positively correlated with accumulation of D/L-2-hydroxyglutarate in the blood [50]. In the sheep this metabolite does not seem to play a role in AF stabilization but might be an interesting metabolite to follow to track atrial rhythm disturbances in patients. Future experiments evaluating the temporality of both TCA cycle remodeling and succinate buildup during AF progression, in relation to electrophysiological abnormalities, would help to decipher if and how succinate accumulation and mitochondrial redox alteration participate in AF progression and/or stabilization. It would also be interesting to compare TCA fluxes between AF-R and AF-S atria in order to precisely evaluate the redox state of the mitochondrial compartment between AF sensitive and resistant animals.

### The potential role of succinate accumulation and oxidation

In mouse permeabilized cardiac mitochondria, succinate oxidation by complex II is favored over NADH oxidation by complex I [53]. This phenomenon could explain the impaired NADH oxidation by complex I (Figures 5A and B), decreased OXPHOS efficiency (Figure 4E), and increased ROS emission (Figure 4I) observed in AF-S isolated mitochondria during succinate oxidation. The increased succinate in the matrix of AF-S mitochondria should participate to the reduced OXPHOS efficiency following complex I bypass. The competition between mitochondrial CI and CII to fuel the ETC is regulated by oxaloacetate [53], an endogenous natural metabolic inhibitor of complex II [54]. Moreover, the calculated [OAA]_Mito_ were equivalent in both the Sham and AF-S groups, thus in the AF-S group no metabolic competition could decrease succinate oxidation at the level of complex II.

A known effect of succinate accumulation is the stimulation of ROS overproduction at the level of complex I [32], although the global ROS emission is directly dependent on the Δp which experimentally is maintained really high in order to observe succinate-driven ROS emission. In patients, the principal source of ROS emission shifts from the cytosol to mitochondrial complex I during AF progression [11]. In this study [11], ROS emission was sensitive to rotenone, an inhibitor of complex I, which inhibits succinate-induced ROS emission. We found that S1QEL, a specific inhibitor of succinate-induced ROS emission at the level of complex I [32], blocked succinate-induced ROS overproduction in AF-S in non-phosphorylating conditions (i.e. high Δp). On isolated mitochondria, during succinate oxidation, mitochondrial inner membrane hyperpolarized, the NAD(P)H pool was more reduced, while *in vivo* it seems that despite succinate accumulation the mitochondrial [NAD^+^] to [NADH] ratio is increased and Δp decreased. Thus, although difficult to extrapolate the exact impact of succinate accumulation on mitochondrial function *in vivo* there is no doubt that its accumulation *in vivo* would lead to an altered function of the organelle.

In AF-S mitochondria, inner membrane hyperpolarization during succinate oxidation is paralleled by an increased rate of calcium uptake. At this stage the origin of such phenomenon is not known especially in condition where potentially mitochondrial heterogeneity is rather different in terms of mitochondrial size. So, further studies are needed to examine the effects of enhanced mitochondrial ROS emission in AF-S animals on sarcoplasmic reticulum calcium handling, which is involved in promoting AF-susceptibility [55]. Nonetheless, several studies have shown that AF is associated with an increased calcium leak from the RyR2 which ultimately end in an increased ATP consumption. AF-R atria did show adaptation that likely favor electron fueling at complex I level and PCr output while preserving a matrix redox state facilitating ROS. These adaptations might be part of the mechanisms contributing to AF resistance.

Exogenous application of succinate on RACM increased (i) intracellular concentrations of succinate, fumarate and L-malate, (2) ROS emission and NAD(P)H fluorescence (like in isolated mitochondria) in the left atrial appendage of the perfused heart. Thus, the rat atrial compartment metabolizes exogenous succinate when applied at a non-physiological dose of 5 mmol/L. Exogenous succinate administration *in vivo* heightened AF susceptibility and stability. The mitigation of succinate’s effects *in vivo* with S1QEL, along with the inhibitory effect of S1QEL on mitochondrial succinate-induced ROS emission, suggest a link between increased succinate, ROS emission and AF vulnerability. Succinate also has a paracrine effects in different tissues [56–58], but so far nothing is known about a putative paracrine effect on atrial electrophysiology.

Our work strongly suggests that TCA cycle remodeling, succinate overload, mitochondrial redox crisis and atrial electrophysiological instability are part of biological mechanisms favoring AF domestication.

### Potential Limitations

The timing of bioenergetic remodeling and AF domestication should be studied to evaluate the causality between TCA cycle remodeling and AF domestication. The rate of OXPHOS and the substrates preferentially oxidized by mitochondria within the fibrillating atrium *in vivo* being unknown, the mitochondrial dysfunction observed *in vitro*. Great caution should be taken when linking succinate buildup to mitochondrial ROS emission *in vivo*. Indeed, mitochondrial succinate-driven ROS emission is regulated by Δp, and its value is not known in vivo in the fibrillating atria. Our findings regarding acute AF induction in rats need to be extrapolated with caution to chronic AF in sheep or humans.

## Conclusions

This is the first study to our knowledge to investigate the bioenergetic mechanisms underlying resistance to AF domestication in a large-animal model. Our results suggest that disturbed mitochondrial succinate metabolism and altered mitochondrial redox balance may play an important role in sensitivity to AF domestication.

There appears to be a link between TCA cycle remodeling, succinate buildup, OXPHOS efficiency regulation, mitochondrial redox balance and AF progression. This work suggests that understanding the regulation of the TCA cycle during AF progression might help to develop new therapeutic strategies aiming to prevent AF progression in humans.

## Funding

This work was supported by the “Agence Nationale de la Recherche (ANR-17-CE14-0029-01 (PJ) and ANR-10-IAHU-04)”, the Canadian Institutes of Health Research (SN), the Heart and Stroke Foundation (SN), the Fondation Lefoulon-Delalande (GC), the Foundation Bordeaux University (GC), the French Federation of Cardiology (GC), and the Deutsche Forschungsgemeinschaft (DFG, German Research Foundation) - project number 514892030 (MP). The Integrion ionic chromatography station was acquired with the financial support of SIRIC BRIO2 (COMMUCAN) and Region Nouvelle-Aquitaine (AAPPF2021-2020- 12000110). The ICS600 chromatography station (ICS6000) and the high-resolution mass spectrometer (Exploris 120) were acquired with financial support from the “Contrat de Projet État Région” 2021-2027 (Biomarker OMICS to BP) and from the ITMO Cancer of Aviesan (I23007FS to BP) as part of the 2021-2030 Cancer Control Strategy, with funds administered by INSERM.

## Author Contribution Statement

GC: navigation catheter during electrophysiological contact mapping and analysis, histology analysis, *in vivo* and *ex vivo* experiments in the rodent models, telemetric and proteomic data analysis, manuscript writing and revision. SN: experimental design and data analysis, guided all aspects of manuscript preparation and provided extensive revision prior to submission of the final manuscript. FV and SC: Western-Blots. PN and FG: proteomic data bioinformatic analysis. SC: proteomic data production by mass spectrometry. BP: metabolomic study using HPIC and mass spectrometry. BB: system analysis at thermodynamic equilibrium. VL, AH, and SP: sheep surgery. MC and MP: histology. AB: sample preparation and biochemical analysis for *in vivo* rodent models. HAM and RD: MATLAB pipeline development used to analyze telemetric recordings. BG: heart tissue sampling. PK and TK: electrophysiological contact mapping. OB: funding and advise on study development. PJ: funding (P.I) and electrophysiological contact mapping supervision. PDS: funding, study supervision and revision prior to submission of the final manuscript. PP: funding, study design and supervision, work on isolated mitochondria, biochemistry, enzymatic, metabolomic, analysis at equilibrium, wrote the first draft of the paper and revised the manuscript.

## Acknowledgements

The authors thank Yohan Bommet, Endy Pierre, Stéphane Bloquet and Nicolas Carles from IHU LIRYC animal facility for their technical assistance and animal care. Authors thank St Jude Medical, now part of Abbott Laboratories, for the donation of the custom-programmed pacemaker. Authors thank Cyrille Casset from Abbott Laboratories and Valentin Meillet from Biosense Webster for their help in pacing device programming and electrophysiological 3D mapping training, respectively.

The authors thank the technical staff of the “Service d’Analyses Métaboliques” at TBMCore (Bordeaux University, CNRS UAR 3427, INSERM US05) for their assistance in metabolite analysis using liquid chromatography coupled with either conductimetry or mass spectrometry. None of the content presented in the present manuscript has been generated using AI. **Conflict of Interest**: The study was conducted with Research Academic funds only. None of the Authors have relationships with the industry relevant to the content of this paper. The only contribution from industry was the donation of the custom programmed pacemakers that we received from St Jude Medical, now part of Abbott Laboratories.

## Notes

https://doi.org/10.6084/m9.figshare.28269872.v1

### Competing Interest Statement

The authors have declared no competing interest.

## References

[1] J. Andrade, P. Khairy, D. Dobrev, S. Nattel, The clinical profile and pathophysiology of atrial fibrillation: relationships among clinical features, epidemiology, and mechanisms, Circ Res 114 (2014) 1453–1468. 10.1161/CIRCRESAHA.114.303211.

[2] M.C. Wijffels, C.J. Kirchhof, R. Dorland, M.A. Allessie, Atrial fibrillation begets atrial fibrillation. A study in awake chronically instrumented goats, Circulation 92 (1995) 1954–1968. 10.1161/01.cir.92.7.1954.

[3] R.R. De With, E.G. Marcos, E.A.M.P. Dudink, H.M. Spronk, H.J.G.M. Crijns, M. Rienstra, I.C. Van Gelder, Atrial fibrillation progression risk factors and associated cardiovascular outcome in well-phenotyped patients: data from the AF-RISK study, Europace 22 (2020) 352–360. 10.1093/europace/euz339.

[4] T. Ito, T. Noda, K. Nochioka, T. Shiroto, N. Yamamoto, H. Sato, T. Chiba, Y. Hasebe, M. Nakano, H. Takahama, J. Takahashi, S. Miyata, H. Shimokawa, S. Yasuda, Clinical impact of atrial fibrillation progression in patients with heart failure with preserved ejection fraction: A report from the CHART-2 Study, Europace 26 (2024) euae218. 10.1093/europace/euae218.

[5] R. Proietti, A. Hadjis, A. AlTurki, G. Thanassoulis, J.-F. Roux, A. Verma, J.S. Healey, M.L. Bernier, D. Birnie, S. Nattel, V. Essebag, A Systematic Review on the Progression of Paroxysmal to Persistent Atrial Fibrillation: Shedding New Light on the Effects of Catheter Ablation, JACC Clin Electrophysiol 1 (2015) 105–115. 10.1016/j.jacep.2015.04.010.

[6] Y. Koide, M. Yotsukura, H. Ando, S. Aoki, T. Suzuki, K. Sakata, E. Ootomo, H. Yoshino, Usefulness of P-wave dispersion in standard twelve-lead electrocardiography to predict transition from paroxysmal to persistent atrial fibrillation, Am J Cardiol 102 (2008) 573–577. 10.1016/j.amjcard.2008.04.065.

[7] G.J. Padfield, C. Steinberg, J. Swampillai, H. Qian, S.J. Connolly, P. Dorian, M.S. Green, K.H. Humphries, G.J. Klein, R. Sheldon, M. Talajic, C.R. Kerr, Progression of paroxysmal to persistent atrial fibrillation: 10-year follow-up in the Canadian Registry of Atrial Fibrillation, Heart Rhythm 14 (2017) 801–807. 10.1016/j.hrthm.2017.01.038.

[8] L. Emelyanova, Z. Ashary, M. Cosic, U. Negmadjanov, G. Ross, F. Rizvi, S. Olet, D. Kress, J. Sra, A.J. Tajik, E.L. Holmuhamedov, Y. Shi, A. Jahangir, Selective downregulation of mitochondrial electron transport chain activity and increased oxidative stress in human atrial fibrillation, Am J Physiol Heart Circ Physiol 311 (2016) H54–63. 10.1152/ajpheart.00699.2015.

[9] M. Mayr, S. Yusuf, G. Weir, Y.-L. Chung, U. Mayr, X. Yin, C. Ladroue, B. Madhu, N. Roberts, A. De Souza, S. Fredericks, M. Stubbs, J.R. Griffiths, M. Jahangiri, Q. Xu, A.J. Camm, Combined metabolomic and proteomic analysis of human atrial fibrillation, J Am Coll Cardiol 51 (2008) 585–594. 10.1016/j.jacc.2007.09.055.

[10] G.N. Kanaan, D.A. Patten, C.J. Redpath, M.-E. Harper, Atrial Fibrillation Is Associated With Impaired Atrial Mitochondrial Energetics and Supercomplex Formation in Adults With Type 2 Diabetes, Can J Diabetes 43 (2019) 67–75.e1. 10.1016/j.jcjd.2018.05.007.

[11] S.N. Reilly, R. Jayaram, K. Nahar, C. Antoniades, S. Verheule, K.M. Channon, N.J. Alp, U. Schotten, B. Casadei, Atrial sources of reactive oxygen species vary with the duration and substrate of atrial fibrillation: implications for the antiarrhythmic effect of statins, Circulation 124 (2011) 1107–1117. 10.1161/CIRCULATIONAHA.111.029223.

[12] Y.H. Kim, D.S. Lim, J.H. Lee, W.J. Shim, Y.M. Ro, G.H. Park, K.G. Becker, Y.S. Cho-Chung, M.-K. Kim, Gene expression profiling of oxidative stress on atrial fibrillation in humans, Exp Mol Med 35 (2003) 336–349. 10.1038/emm.2003.45.

[13] D. Montaigne, X. Marechal, P. Lefebvre, T. Modine, G. Fayad, H. Dehondt, C. Hurt, A. Coisne, M. Koussa, I. Remy-Jouet, F. Zerimech, E. Boulanger, D. Lacroix, B. Staels, R. Neviere, Mitochondrial dysfunction as an arrhythmogenic substrate: a translational proof-of-concept study in patients with metabolic syndrome in whom post-operative atrial fibrillation develops, J Am Coll Cardiol 62 (2013) 1466–1473. 10.1016/j.jacc.2013.03.061.

[14] P.H. Lin, S.H. Lee, C.P. Su, Y.H. Wei, Oxidative damage to mitochondrial DNA in atrial muscle of patients with atrial fibrillation, Free Radic Biol Med 35 (2003) 1310–1318. 10.1016/j.freeradbiomed.2003.07.002.

[15] R.P. Martins, K. Kaur, E. Hwang, R.J. Ramirez, D. Filgueiras-Rama, S.R. Ennis, Y. Takemoto, D. Ponce-Balbuena, Dominant frequency increase rate predicts transition from paroxysmal to long-term persistent atrial fibrillation, Circulation 129 (2014) 1472–1482. 10.1161/CIRCULATIONAHA.113.004742.

[16] A. Alvarez-Franco, R. Rouco, R.J. Ramirez, G. Guerrero-Serna, M. Tiana, S. Cogliati, K. Kaur, M. Saeed, R. Magni, J.A. Enriquez, F. Sanchez-Cabo, J. Jalife, M. Manzanares, Transcriptome and proteome mapping in the sheep atria reveal molecular featurets of atrial fibrillation progression, Cardiovasc Res 117 (2021) 1760–1775. 10.1093/cvr/cvaa307.

[17] J. Schindelin, I. Arganda-Carreras, E. Frise, V. Kaynig, M. Longair, T. Pietzsch, S. Preibisch, C. Rueden, S. Saalfeld, B. Schmid, J.-Y. Tinevez, D.J. White, V. Hartenstein, K. Eliceiri, P. Tomancak, A. Cardona, Fiji: an open-source platform for biological-image analysis, Nat Methods 9 (2012) 676–682. 10.1038/nmeth.2019.

[18] L. Käll, J.D. Canterbury, J. Weston, W.S. Noble, M.J. MacCoss, Semi-supervised learning for peptide identification from shotgun proteomics datasets, Nat Methods 4 (2007) 923–925. 10.1038/nmeth1113.

[19] M. Milacic, D. Beavers, P. Conley, C. Gong, M. Gillespie, J. Griss, R. Haw, B. Jassal, L. Matthews, B. May, R. Petryszak, E. Ragueneau, K. Rothfels, C. Sevilla, V. Shamovsky, R. Stephan, K. Tiwari, T. Varusai, J. Weiser, A. Wright, G. Wu, L. Stein, H. Hermjakob, P. D’Eustachio, The Reactome Pathway Knowledgebase 2024, Nucleic Acids Research 52 (2024) D672–D678. 10.1093/nar/gkad1025.

[20] Z. Pang, L. Xu, C. Viau, Y. Lu, R. Salavati, N. Basu, J. Xia, MetaboAnalystR 4.0: a unified LC-MS workflow for global metabolomics, Nat Commun 15 (2024) 3675. 10.1038/s41467-024-48009-6.

[21] P. Pasdois, B. Beauvoit, L. Tariosse, B. Vinassa, S. Bonoron-Adèle, P.D. Santos, Effect of diazoxide on flavoprotein oxidation and reactive oxygen species generation during ischemia-reperfusion: a study on Langendorff-perfused rat hearts using optic fibers, American Journal of Physiology-Heart and Circulatory Physiology 294 (2008) H2088–H2097. 10.1152/ajpheart.01345.2007.

[22] T. Andrienko, P. Pasdois, A. Rossbach, A.P. Halestrap, Real-Time Fluorescence Measurements of ROS and [Ca2+] in Ischemic / Reperfused Rat Hearts: Detectable Increases Occur only after Mitochondrial Pore Opening and Are Attenuated by Ischemic Preconditioning, PLoS ONE 11 (2016) e0167300. 10.1371/journal.pone.0167300.

[23] R. Brandes, V.M. Figueredo, S.A. Camacho, A.J. Baker, M.W. Weiner, Quantitation of cytosolic [Ca2+] in whole perfused rat hearts using Indo-1 fluorometry, Biophysical Journal 65 (1993) 1973–1982. 10.1016/S0006-3495(93)81274-8.

[24] P. Pasdois, J.E. Parker, A.P. Halestrap, Extent of Mitochondrial Hexokinase II Dissociation During Ischemia Correlates With Mitochondrial Cytochrome c Release, Reactive Oxygen Species Production, and Infarct Size on Reperfusion, JAHA 2 (2013) e005645. 10.1161/JAHA.112.005645.

[25] M. Nishimura, T. Ito, B. Chance, Studies on bacterial photophosphorylation. III. A sensitive and rapid method of determination of photophosphorylation, Biochim Biophys Acta 59 (1962) 177– 182.

[26] S. Dufour, N. Rousse, P. Canioni, P. Diolez, Top-down control analysis of temperature effect on oxidative phosphorylation, Biochemical Journal 314 (1996) 743–751. 10.1042/bj3140743.

[27] A.J. Moreno, D.L. Santos, S. Magalhães-Novais, P.J. Oliveira, Measuring Mitochondrial Membrane Potential with a Tetraphenylphosphonium-Selective Electrode, Current Protocols in Toxicology 65 (2015). 10.1002/0471140856.tx2505s65.

[28] N. Kamo, M. Muratsugu, R. Hongoh, Y. Kobatake, Membrane potential of mitochondria measured with an electrode sensitive to tetraphenyl phosphonium and relationship between proton electrochemical potential and phosphorylation potential in steady state, J Membr Biol 49 (1979) 105–121. 10.1007/BF01868720.

[29] A.P. Halestrap, The regulation of the oxidation of fatty acids and other substrates in rat heart mitochondria by changes in the matrix volume induced by osmotic strength, valinomycin and Ca2+, Biochemical Journal 244 (1987) 159–164. 10.1042/bj2440159.

[30] I. Ceballos-Picot, A. Le Dantec, A. Brassier, J.-P. Jaïs, M. Ledroit, J. Cahu, H.-K. Ea, B. Daignan-Fornier, B. Pinson, New biomarkers for early diagnosis of Lesch-Nyhan disease revealed by metabolic analysis on a large cohort of patients, Orphanet J Rare Dis 10 (2015) 7. 10.1186/s13023-014-0219-0.

[31] O.O. Mesubi, A.G. Rokita, N. Abrol, Y. Wu, B. Chen, Q. Wang, J.M. Granger, A. Tucker-Bartley, E.D. Luczak, K.R. Murphy, P. Umapathi, P.S. Banerjee, T.N. Boronina, R.N. Cole, L.S. Maier, X.H. Wehrens, J.L. Pomerantz, L.-S. Song, R.S. Ahima, G.W. Hart, N.E. Zachara, M.E. Anderson, Oxidized CaMKII and O-GlcNAcylation cause increased atrial fibrillation in diabetic mice by distinct mechanisms, J Clin Invest 131 (2021) e95747, 95747. 10.1172/JCI95747.

[32] H.-S. Wong, P.-A. Monternier, M.D. Brand, S1QELs suppress mitochondrial superoxide/hydrogen peroxide production from site IQ without inhibiting reverse electron flow through Complex I, Free Radical Biology and Medicine 143 (2019) 545–559. 10.1016/j.freeradbiomed.2019.09.006.

[33] H.A. Prag, A.V. Gruszczyk, M.M. Huang, T.E. Beach, T. Young, L. Tronci, E. Nikitopoulou, J.F. Mulvey, R. Ascione, A. Hadjihambi, M.J. Shattock, L. Pellerin, K. Saeb-Parsy, C. Frezza, A.M. James, T. Krieg, M.P. Murphy, D. Aksentijević, Mechanism of succinate efflux upon reperfusion of the ischaemic heart, Cardiovasc Res 117 (2021) 1188–1201. 10.1093/cvr/cvaa148.

[34] A. Reddy, L.H.M. Bozi, O.K. Yaghi, E.L. Mills, H. Xiao, H.E. Nicholson, M. Paschini, J.A. Paulo, R. Garrity, D. Laznik-Bogoslavski, J.C.B. Ferreira, C.S. Carl, K.A. Sjøberg, J.F.P. Wojtaszewski, J.F. Jeppesen, B. Kiens, S.P. Gygi, E.A. Richter, D. Mathis, E.T. Chouchani, pH-Gated Succinate Secretion Regulates Muscle Remodeling in Response to Exercise, Cell 183 (2020) 62–75.e17. 10.1016/j.cell.2020.08.039.

[35] F.E. Mason, J.R.D. Pronto, K. Alhussini, C. Maack, N. Voigt, Cellular and mitochondrial mechanisms of atrial fibrillation, Basic Res Cardiol 115 (2020) 72. 10.1007/s00395-020-00827-7.

[36] Y.-M. Cha, P.P. Dzeja, W.K. Shen, A. Jahangir, C.Y.T. Hart, A. Terzic, M.M. Redfield, Failing atrial myocardium: energetic deficits accompany structural remodeling and electrical instability, Am J Physiol Heart Circ Physiol 284 (2003) H1313–1320. 10.1152/ajpheart.00337.2002.

[37] M. Tsuboi, I. Hisatome, T. Morisaki, M. Tanaka, Y. Tomikura, S. Takeda, M. Shimoyama, A. Ohtahara, K. Ogino, O. Igawa, C. Shigemasa, S. Ohgi, E. Nanba, Mitochondrial DNA deletion associated with the reduction of adenine nucleotides in human atrium and atrial fibrillation, Eur J Clin Invest 31 (2001) 489–496. 10.1046/j.1365-2362.2001.00844.x.

[38] L. Pool, L.F.J.M. Wijdeveld, N.M.S. de Groot, B.J.J.M. Brundel, The Role of Mitochondrial Dysfunction in Atrial Fibrillation: Translation to Druggable Target and Biomarker Discovery, Int J Mol Sci 22 (2021) 8463. 10.3390/ijms22168463.

[39] J.H. Rennison, L. Li, C.R. Lin, B.S. Lovano, L. Castel, S.Y. Wass, C.C. Cantlay, M. McHale, A.M. Gillinov, R. Mehra, B.B. Willard, J.D. Smith, M.K. Chung, J. Barnard, D.R. Van Wagoner, Atrial fibrillation rhythm is associated with marked changes in metabolic and myofibrillar protein expression in left atrial appendage, Pflugers Arch 473 (2021) 461–475. 10.1007/s00424-021-02514-5.

[40] A. Alvarez-Franco, R. Rouco, R.J. Ramirez, G. Guerrero-Serna, M. Tiana, S. Cogliati, K. Kaur, M. Saeed, R. Magni, J.A. Enriquez, F. Sanchez-Cabo, J. Jalife, M. Manzanares, Transcriptome and proteome mapping in the sheep atria reveal molecular featurets of atrial fibrillation progression, Cardiovasc Res 117 (2021) 1760–1775. 10.1093/cvr/cvaa307.

[41] C. Ozcan, Z. Li, G. Kim, V. Jeevanandam, N. Uriel, Molecular Mechanism of the Association Between Atrial Fibrillation and Heart Failure Includes Energy Metabolic Dysregulation Due to Mitochondrial Dysfunction, J Card Fail 25 (2019) 911–920. 10.1016/j.cardfail.2019.08.005.

[42] M. Wiersma, D.M.S. van Marion, R.C.I. Wüst, R.H. Houtkooper, D. Zhang, N.M.S. de Groot, R.H. Henning, B.J.J.M. Brundel, Mitochondrial Dysfunction Underlies Cardiomyocyte Remodeling in Experimental and Clinical Atrial Fibrillation, Cells 8 (2019) 1202. 10.3390/cells8101202.

[43] M.J. Mihm, F. Yu, C.A. Carnes, P.J. Reiser, P.M. McCarthy, D.R. Van Wagoner, J.A. Bauer, Impaired myofibrillar energetics and oxidative injury during human atrial fibrillation, Circulation 104 (2001) 174–180. 10.1161/01.cir.104.2.174.

[44] J. Ausma, W.A. Coumans, H. Duimel, G.J. Van der Vusse, M.A. Allessie, M. Borgers, Atrial high energy phosphate content and mitochondrial enzyme activity during chronic atrial fibrillation, Cardiovasc Res 47 (2000) 788–796. 10.1016/s0008-6363(00)00139-5.

[45] M. Harada, J. Melka, Y. Sobue, S. Nattel, Metabolic Considerations in Atrial Fibrillation - Mechanistic Insights and Therapeutic Opportunities, Circ J 81 (2017) 1749–1757. 10.1253/circj.CJ-17-1058.

[46] M. de Castro Fonseca, C.J. Aguiar, J.A. da Rocha Franco, R.N. Gingold, M.F. Leite, GPR91: expanding the frontiers of Krebs cycle intermediates, Cell Commun Signal 14 (2016) 3. 10.1186/s12964-016-0126-1.

[47] W. Shan, H. Cui, Y. Xu, J. Xue, L. Zheng, Succinate metabolism in cardiovascular diseases, Global Translational Medicine 1 (2022) 160. 10.36922/gtm.v1i2.160.

[48] S.E. Nelson, Z. Ament, Z. Wolcott, R.E. Gerszten, W.T. Kimberly, Succinate links atrial dysfunction and cardioembolic stroke, Neurology 92 (2019) e802–e810. 10.1212/WNL.0000000000006957.

[49] F.J. Osuna-Prieto, B. Martinez-Tellez, L. Ortiz-Alvarez, X. Di, L. Jurado-Fasoli, H. Xu, V. Ceperuelo-Mallafré, C. Núñez-Roa, I. Kohler, A. Segura-Carretero, J.V. García-Lario, A. Gil, C.M. Aguilera, J.M. Llamas-Elvira, P.C.N. Rensen, J. Vendrell, J.R. Ruiz, S. Fernández-Veledo, Elevated plasma succinate levels are linked to higher cardiovascular disease risk factors in young adults, Cardiovascular Diabetology 20 (2021) 151. 10.1186/s12933-021-01333-3.

[50] M. Bulló, C. Papandreou, J. García-Gavilán, M. Ruiz-Canela, J. Li, M. Guasch-Ferré, E. Toledo, C. Clish, D. Corella, R. Estruch, E. Ros, M. Fitó, C.-H. Lee, K. Pierce, C. Razquin, F. Arós, L. Serra-Majem, L. Liang, M.A. Martínez-González, F.B. Hu, J. Salas-Salvadó, Tricarboxylic acid cycle related-metabolites and risk of atrial fibrillation and heart failure, Metabolism 125 (2021) 154915. 10.1016/j.metabol.2021.154915.

[51] E.T. Chouchani, V.R. Pell, E. Gaude, D. Aksentijević, S.Y. Sundier, E.L. Robb, A. Logan, S.M. Nadtochiy, E.N.J. Ord, A.C. Smith, F. Eyassu, R. Shirley, C.-H. Hu, A.J. Dare, A.M. James, S. Rogatti, R.C. Hartley, S. Eaton, A.S.H. Costa, P.S. Brookes, S.M. Davidson, M.R. Duchen, K. Saeb-Parsy, M.J. Shattock, A.J. Robinson, L.M. Work, C. Frezza, T. Krieg, M.P. Murphy, Ischaemic accumulation of succinate controls reperfusion injury through mitochondrial ROS, Nature 515 (2014) 431–435. 10.1038/nature13909.

[52] K.A. van Bragt, H.M. Nasrallah, M. Kuiper, J.J. Luiken, U. Schotten, S. Verheule, Atrial supply-demand balance in healthy adult pigs: coronary blood flow, oxygen extraction, and lactate production during acute atrial fibrillation, Cardiovasc Res 101 (2014) 9–19. 10.1093/cvr/cvt239.

[53] T. Molinié, E. Cougouilles, C. David, E. Cahoreau, J.-C. Portais, A. Mourier, MDH2 produced OAA is a metabolic switch rewiring the fuelling of respiratory chain and TCA cycle, Biochimica et Biophysica Acta (BBA) – Bioenergetics 1863 (2022) 148532. 10.1016/j.bbabio.2022.148532.

[54] L. Wojtczak, A.B. Wojtczak, L. Ernster, The inhibition of succinate dehydrogenase by oxalacetate, Biochim Biophys Acta 191 (1969) 10–21. 10.1016/0005-2744(69)90310-6.

[55] H. Dridi, A. Kushnir, R. Zalk, Q. Yuan, Z. Melville, A.R. Marks, Intracellular calcium leak in heart failure and atrial fibrillation: a unifying mechanism and therapeutic target, Nat Rev Cardiol 17 (2020) 732–747. 10.1038/s41569-020-0394-8.

[56] M.P. Murphy, L.A.J. O’Neill, Krebs Cycle Reimagined: The Emerging Roles of Succinate and Itaconate as Signal Transducers, Cell 174 (2018) 780–784. 10.1016/j.cell.2018.07.030.

[57] M.P. Murphy, E.T. Chouchani, Why succinate? Physiological regulation by a mitochondrial coenzyme Q sentinel, Nat Chem Biol 18 (2022) 461–469. 10.1038/s41589-022-01004-8.

[58] D.G. Ryan, M.P. Murphy, C. Frezza, H.A. Prag, E.T. Chouchani, L.A. O’Neill, E.L. Mills, Coupling Krebs cycle metabolites to signalling in immunity and cancer, Nat Metab 1 (2019) 16–33. 10.1038/s42255-018-0014-7.

