## Supplemental Materials for "Resistance to Atrial Fibrillation Domestication and Mitochondrial Dysfunction in Sheep: a potential key role of the TCA Cycle and mitochondrial redox state"

**Short title: AF resistance and mitochondrial adaptation**

**Author's' list:** Guido Caluori PhD [a], Stanley Nattel MD [c], Benoît Pinson PhD [d], Patrice Naud PhD [c], Bertrand Beauvoit [e], Stéphane Claverol PhD [f], Fanny Vaillant PhD [a], Sabine Charron MsC [a], Florent Guilloteau BSc [g], , Virginie Loyer [a], Marion Constantin MsC [a], Miruna Popa MD [a], Andrei Belykh PhD [a], Andreas Haeberlin MD, PhD [a], Sylvain Ploux MD, PhD [a,b], Hassan-Adam Mahamat PhD [a], Rémi Dubois PhD [a], Bastien Guillot PhD [a], Philipp Krisai MD [a], Tsukasa Kamakura MD [a], Olivier Bernus PhD [a], Pierre Jaïs MD [a,b] , Pierre Dos Santos MD, PhD [a,b], Philippe Pasdois PhD [a, h]

**Departments and Institutions:**

a. Univ. Bordeaux, INSERM, CRCTB, U 1045, IHU Liryc, F-33000 Bordeaux, France

b. CHU de Bordeaux, INSERM, U 1045, F-33000 Bordeaux, France

c. Department of Medicine and Research Center, Montreal Heart Institute and Université de Montréal, Montréal, Québec, Canada; Department of Pharmacology and Therapeutics, McGill University, Montréal, Québec, Canada; Institute of Pharmacology, West German Heart and Vascular Center, University of Duisburg-Essen, Essen, Germany.

d. Service d'Analyses Métaboliques- TBMCore, CNRS UAR 3427, Inserm US 005, Univ. Bordeaux, France

e. Univ. Bordeaux, INRAE, BFP, UMR 1332, F-33140 Villenave d'Ornon, France

f. Plateforme Protéome, Univ. Bordeaux, Bordeaux, France

g. Laboratoire de diagnostic moléculaire, hôpital Maisonneuve-Rosemont, Montréal, Québec, Canada

h. Current address: Univ. Bordeaux, CNRS, IBGC, UMR 5095, F-33000 Bordeaux, France

**Address for Correspondence:**

Philippe Pasdois. Université de Bordeaux, Institut de Biochimie et Génétique Cellulaires CNRS-UMR5095, 1, rue Camille Saint-Saëns, F-33077 Bordeaux, France. Tel : +33556999064.

**Article category**: Original article

### Supplemental Methods

***Antibodies and chemicals***.

The antibodies used in this study are listed in supplementary table 1. All chemicals used in this study were purchased from Sigma unless otherwise stated.

***Ethics***.

All experiments were performed following the guidelines from Directive 2010/63/EU of the European Parliament on the protection of animals used for scientific purposes. Protocols applied to sheep and rats were approved by the local ethical committee (CEEA50) at the University of Bordeaux and by the French Government (authorization references APAFIS#20364 and APAFIS#29528, for sheep and rat, respectively).

***AF sheep model***.

In this study, we used a modified version of the atrial fibrillation (AF) model extensively described by Jalife and colleagues.[1] Thirty-two female sheep (Charmoise, mean age = 8.4±0.1 years, mean weight = 50.8+/-1.2 kg) were implanted with a sterile dual chamber pacemaker (St. Jude Medical, Inc, St Paul, MN), two pacing leads (IS1 leads Tendril STS 2088TC, St. Jude Medical, Inc, St Paul, MN), and a telemetric device (EasyTEL+L – ETA, Emka Technologies, Paris, FR). We used older sheep because in preliminary studies we found that there was a high incidence of AF-resistance in younger sheep.

Sheep were prepared for the surgical procedure after an intramuscular injection of ketamine (20 mg/Kg), acepromazine (0.1 mg/Kg) and buprenorphine (9 µg/Kg). After installation of a venous line in the left front leg, anesthesia was induced by an intravenous bolus of propofol (1 mg/Kg). Sheep were then rapidly intubated and maintained under anesthesia using an equal mixture of air and oxygen supplemented by isoflurane (1.5 to 3%). Temperature, arterial pressure, $PO_2$, $PCO_2$, and ECG were continuously monitored throughout the surgical procedure (Care Station 650, GE HealthCare).

The pacing leads were positioned in the right atrial appendage and at the apex of the right ventricle through the left jugular vein using 8 Fr peelable introducers. Measurements of impedance at the tip of the electrodes, electric signal amplitude and capture threshold were used to correctly position the leads during the surgical procedure. Once optimal lead positions were found, their distal ends were screwed into the myocardial walls and their proximal ends secured to the atrial and ventricular ports of the pacemaker. The pacing device was then positioned and secured at the base of the neck within a submuscular pocket. The right ventricular lead was exclusively used to monitor ventricular rhythm during the protocol and the ventricular pacing option was disabled in the pacemaker.

Sheep were also equipped with a telemetric device (EasyTEL+L – ETA, Emka Technologies, Paris, France) composed of one solid-tip lead positioned just before the entrance of the right atria through the left jugular vein, and one subcutaneous spring-tip lead positioned opposite the solid tip at the top of the thorax. This relative position of the two biopotential probes allowed us to obtain similar amplitudes of P waves and QRS complexes on the measured pseudo-ECG. The telemetric equipment was used to record sheep-heart electrical activity and temperature continuously over an average follow-up period of 127±5 days using a signal resolution of 10 bits, a range of ±5 mV, and sampling rate of 500Hz.

After 7 to 10 days of recovery post-surgery, the pacemaker was activated to either sense and record the right atrial and ventricular signals (Sham group, N=11) or to sense, record and pace the right atria to induce AF (Electrostimulation group, N=21). The pacing voltage was set to 5V, while typical capture threshold at 0.5-ms duration were below 1V. The pacing rate was set to 10Hz and burst stimulation duration to 30 seconds, after which the pacemaker algorithm sensed the atrial rate for 7 seconds. If, during this non-stimulated period, the atrial rate was below 180 bpm, a new 30-second burst was delivered, otherwise an episode of AF had been triggered and the device stopped stimulating until the atrial rate dropped below 180 bpm (restoration of sinus rhythm). This experimental approach is meant to lead to the development of self-sustained AF due to AF-related remodeling. AF progression rate and stability were continuously monitored using the telemetric platform coupled to a MATLAB script (MathWorks, Natick, Massachusetts, USA) developed in the laboratory, analyzing the telemetric signals to extract the number of electrical bursts received per 24 hours.

***Experimental design.***

Three groups of animals were analyzed, the Sham-operated group (Sham), the AF-tachypaced group composed of all animals who developed stable AF (AF-sensitive, AF-S), and the resistant tachypaced group considered resistant because the animal did not develop stable AF despite an average of 120±7 days of tachypacing (AF-resistant, AF-R). Most animals underwent atrial endocardial electro-anatomical mapping before surgical implantation and at the end of the follow-up period to evaluate the impact of tachypacing and AF on atrial electrophysiological remodeling. Five days after the terminal *in-vivo* electrophysiological study, sheep hearts were explanted to perform an extended biochemical and bioenergetic analysis of the left atrial compartment.

***Atrial endocardial electrical contact mapping.***

The electroanatomical mapping experiments were performed on a CARTO™ 3 PIU using CARTO™ v6 software (both from Biosense Webster, Irvine, CA, USA). Female charmoise sheep were sedated and kept under general anesthesia by continuous ventilation with 1-2% isoflurane. Two introducers of 11 F and 8 F were inserted in the femoral veins to allow the introduction of a steerable 8.5 F sheath (Agilis NxT, Abbott Laboratories, Chicago, IL-US) and a 6 F steerable diagnostic catheter (EP XT™, Boston Scientific, Marlborough, MA-US). The sheath was used to maneuver a 20-pole steerable mapping catheter in the atrial chambers (PENTARAY™ NAV, Biosense Webster); the diagnostic catheter was introduced via the coronary sinus of the animals for pacing and monitoring. Heparin was administered as an i.v. bolus before the insertion of the devices and continuously infused *via* the flushing saline bags of the sheath and mapping catheter to avoid thrombus formation. Total activation time (TAT) of the right and the left atrium were calculated as the difference between the average latest and average earliest point of electrograms (EGM) annotation, during an S1 stimulation train at a cycle-length (CL) of 500 ms from the coronary sinus. Local ERP was calculated by pacing the atrial tissue with the mapping catheter, using one extrastimulus with a progressively decreasing S2 interval, following an S1 train at a 500 ms CL. The ERP was calculated as the longest S1-S2 interval at which electrical activation was not observed on the coronary sinus catheter. This parameter was calculated in the following areas: atrial appendage, lateral wall, interatrial septum, posterior wall, anterior wall, inferior/superior vena cava (for the right atrium), and superior/inferior pulmonary vein ostium (for the left atrium).

***Left atrial appendage freeze-clamping and heart explant.***

A thoracotomy was performed under general anesthesia and analgesia. The sheep heart was exposed after opening the pericardium. Before euthanasia and heart excision, a piece of the left atrial appendage (LAA) was freeze-clamped *in-vivo* using liquid-nitrogen cooled tongs and stored at -80°C for latter biochemical analysis (see below). AF-sensitive, AF-resistant, and Sham animals had their tissue clamped during AF, tachypacing and sinus rhythm, respectively. Heart mechanical activity was stopped by aortic retroperfusion of an ice-cold solution of Ringer-Lactate, at a constant pressure of 80 mmHg, and by filling the chest cavity with an ice-cold sodium chloride solution at 9 g/L. After complete cardiac arrest and subsequent disappearance of a detectable arterial pressure, heart was explanted and immersed in ice-cold Tyrode solution containing (in mmol/L): NaCl 137, KCl 5, $MgCl_2$ 1, HEPES 20, glucose 10,

$CaCl_2$ 1.2. The heart was then rapidly cannulated and retroperfused with the cold Tyrode solution to wash all the cavities and coronary circulation of residual blood.

***Atrial histology and structural characterization.***

The left atrial posterior wall was fixed in 4% paraformaldehyde in phosphate buffered saline and further prepared for histological characterization. Fixed tissue slices were dehydrated in their cassette with a HistoCore PEARL (Leica Biosystems, Wetzlar, Germany) using the overnight program 2. The dehydrated sections were embedded in paraffin blocks using a HistoCore Arcardia C (Leica) and kept at -20 °C until use. Thin sections (5-µm) were obtained with an RM2255 microtome (Leica) and stained with Masson trichrome dyes using an Autostainer XL workstation (Leica). The slides were scanned with an Axio Scan.Z1 microscope (ZEISS, Jena, Germany) with 20x magnification in brightfield and the tiles automatically assembled to form the section overview. The images were imported in Fiji[2] for calculation of the parameters of interest. Endomysial fibrosis was estimated as the area fraction of the blue-stained connective tissue in the midmyocardial region, excluding perivascular areas. Cardiomyocyte cross-section area was estimated by measuring and averaging the area of 10-20 regions of interest in transversally-cut fibers, taken at a minimum of three locations. Cell-to-cell distance (CM border-to-border distance) was estimated by measuring the center-to-center distance of the same regions of interest and subtracting the median cross-section diameter.

***LAA tissue preparation for biochemical analysis.***

Pieces of frozen LAA were pulverized under liquid nitrogen, and 100 to 150 mg were mixed with 500 µL of cold perchloric acid (10%) solution supplemented with EDTA (10 mmol/L) to precipitate proteins. Following centrifugation for 7 min at 17,000g, 450µL of supernatant were collected and neutralized using 230µL of $KHCO_3$ (2.4mol/L) supplemented by MOPS (10% w/v) and KOH (10% w/v) leading to an extracted solution at pH 7.2 - 7.4 at room temperature (RT). A final centrifugation for 7 min at 17,000g was performed and the collected supernatant frozen in liquid nitrogen before being stored at -80°C for later biochemical analysis of metabolite content (see main manuscript).

***Measurement of mitochondrial calcium retention capacity assay.***

Mitochondria were incubated at 37°C in the following buffer, KCl 125 mol/L, MOPS 20 mmol/L, Tris 10 mmol/L, EGTA 10 µmol/L, $KH_2PO_4$ 2 mmol/L, pH 7.3 adjusted at RT with KOH. Extra-mitochondrial calcium was monitored using the fluorescent dye calcium green™-5N (Thermo Fisher Scientific) at a final concentration of 1 µmol/L. Calcium spikes (35 µmol/L) were added successively until mitochondria could not buffer calcium in the medium, thus reflecting the maximum calcium retention capacity (CRC) before opening of the mitochondrial permeability transition pore (mPTP). A calibration curve with known calcium concentration was performed to quantify the total extra-mitochondrial calcium. CRC was expressed as µmol of free calcium per mg protein. The free calcium concentration in the buffer was calculated considering the dissociation constant of the different calcium ligand, the quantity of metals and the pH of the assay buffer used. To assess the maximal mitochondrial calcium uptake velocity as a function of calcium quantity in the buffer, the decay slope of each peak was measured. The maximal mitochondrial calcium uptake velocity was then expressed as µmole of free calcium per min per mg protein.

***Rat atrial cardiomyocyte (RACM) isolation***

Male Wistar rats (225-300g, N=10) were anaesthetized using 3% isoflurane and anticoagulated (heparin 10000 UI/Kg) before euthanasia by cervical dislocation. Hearts were rapidly excised and placed into ice-cold Tyrode buffer containing (mmol/L) NaCl 130, KCl 5.4, $MgCl_2$ 1.4, $NaH_2PO_4$ 0.4, HEPES 5, glucose 10, creatine 10, taurine 10, and $CaCl_2$ 0.75 pH 7.4 at RT. Contraction was blocked by adding 0.1 mmol/L EGTA with the same solution gassed with 100% $O_2$ at 37°C retroperfused from the aorta. Collagenase II (1 mg/mL, Worthington Biochemical, Lakewood, NJ-US), protease XIV (0.1 mg/mL) and $CaCl_2$ 20 µmol/L were added to enzymatically digest the extracellular matrix. Atrial tissue was then cut from the heart, chopped and digested in fresh enzyme-supplemented isolation solution for 5-10 mins at 37°C under agitation. Once pelleted, the cells were slowly reintroduced to calcium and then resuspended in Tyrode. containing (mmol/L) NaCl 130, KCl 5.4, $MgCl_2$ 1, HEPES 5, glucose 10, $CaCl_2$ 1.

***Steady-state ROS production in atrial cardiomyocytes***

RACM were allowed to sediment in Tyrode solution on a 35 mm Flurodish (WPI, Sarasota, FL-US) at RT for 1h. They were then incubated for 30 min at 37°C and 5% $CO_2$ in the presence of CellROX ™ Green (Thermo Fisher Scientific, MA-US) at the recommended concentration of 5 µmol/L. To estimate the effect of extracellular succinate on intracellular ROS production, the following conditions were tested:

- Succinate (buffered with HEPES 25 mmol/L at pH 7.4 RT) at 5 mmol/L
- $H_2O_2$ at 50 µmol/L as technical positive control
- Tyrode as negative control

After incubation, brightfield and epifluorescence images were obtained with a Leica Thunder Imager system (Leica Camera, Wetzlar, Germany) with the standard 488/530 excitation/emission filters. At least 10 cells per dish were imaged and results averaged under the same conditions of exposure time and light intensity. Cell-body fluorescence relative to background (F/F0) was calculated using Fiji[2] by dividing the average mean intensity of the region of interest enclosing a cardiomyocyte by the average mean intensity of the cell-free dish surface.

***Atrial surface fluorescence measurements on perfused rat hearts.***

The distal end of a bifurcated optic-fiber cable (5 mm diameter, 6.8 $mm^2$ per bundle) was placed at ~ 0.5 cm of the LAA wall. The two proximal ends of the optic-fiber cable were connected to a modified spectrofluorometer (Xenius, SAFAS Monaco) like previously published.[3] The

NAD(P)H pool redox state was assessed on the beating LAA by recording the autofluorescence collected at $\lambda_{em}$ 460nm ($\lambda_{ex}$ 340nm). Like previously reported the majority of the fluorescent signal recorded at 460nm reflects the pool of NADH and NADPH of the mitochondrial compartment.[4] To evaluate the mitochondrial $H_2O_2$ emission hearts were loaded with the fluorescent probe Mitochondria Peroxy-Yellow 1 (MitoPY1, ($\lambda_{ex}$ 485nm & $\lambda_{em}$ 535nm, Tocris Bioscience, #4428) at a final concentration of 3 µmol/L like previously published[5] with minor modifications. Fluorescent dye loading was performed using a syringe pump, connected to the perfusion cannula, delivering the dye loading solution containing MitoPY1(360 µmol/L and pluronic F127 1.5mg/mL, solubilized in DMSO) at a rate of 100 µL/min. MitoPY1 loading was stopped once the fluorescent signal was ~ 3 to 4 times higher than the initial background signal (~ 30 min loading period). A 20 to 30 min washout was performed until NAD(P)H and MitoPY1 fluorescent signals stabilized. Succinate was then perfused at 5 mmol/L using a syringe pump connected to the perfusion cannula by a three-way stopcock for 10 min. A second washout period was then performed before starting the perfusion of $H_2O_2$ at a final concentration of 400 µmol/L for 15 min. Distance and motion artefact were diminished by dividing the emitted photon intensity by the reflected signals measured at the respective excitation and emission wavelengths.[6]

***In-vivo assessment of susceptibility to atrial arrhythmia in the rat.***

Male Wistar rats (250-300g, N=20) were put under general anesthesia using an induction chamber saturated with 3.5% isoflurane for 3 min (Minerve, Esternay, France). Upon confirming slower and deeper respiration, animals were transferred supine to a heated mat equipped with an anesthesia mask providing a mix of 80% oxygen and 20% air supplemented with 2.5% isoflurane. The animal’s body temperature was measured with a rectal probe and kept at 37-37.5 °C by adjusting the set point of the heating mat. A CIB'ER Mouse® catheter (NuMED, Inc., Hopkinton, NY-USA) was introduced in the animal’s esophagus until a clear atrial electrogram was recorded. Needle electrodes were placed subcutaneously in the front and rear paws to measure lead 1 of the surface ECG. The signals were acquired through a PowerLab 26 Series recorder, visualized and recorded with LabChart proprietary software. A 26G cannula was inserted in the caudal vein and connected to a syringe pump for IV infusions, at rate of 15 mL/kg/h of :

- A 3% w/v solution of NaCl buffered with 5mM HEPES at pH 7.4 – vehicle group (N=6)
- An 80 mg/mL solution of succinic acid buffered with 5mM HEPES to pH 7.4 – succinate group (N=8)
- An 80 mg/mL solution of succinic acid buffered with 5mM HEPES at pH 7.4 preceded by an i.p. injection of 10 µmol/kg of S1QEL1.1 in PBS – S1QEL group (N=6)

The concentration of the vehicle solution was chosen to match the osmolarity of the succinic acid solution, measured between 900-1000 mOsm with an automatic osmometer (Löser Messtechnik, Berlin, Germany). Atrial fibrillation/tachycardia (AF/AT) vulnerability was estimated after 10 min of IV injection by challenging the hearts with up to 10 rapid atrial stimulation trains at 30 Hz for 30 sec. The challenge was repeated up to 4 times/heart after 3 to 5 min of recovery.

***Content of OGA, GFAT2, and OGT by western blotting***.

For immunoblotting, 80 mg of freeze-clamped LAA grounded powder were solubilized in 500 µL of ice-cold RIPA lysis buffer (Merck, Germany, #R0278), containing protease inhibitor cocktail (Sigma-Aldrich, Missouri, United States, #P8340), phosphatase inhibitor cocktail (Sigma-Aldrich, #A32957), 2 mmol/L PMSF (Sigma-Aldrich, 93482) and 50 µmol/L PUGNAc (O-(2-acetamido-2-deoxy-D-glucopyrano-sylidene)amino-N-phenyl-carbamate, Sigma-Aldrich, #A7229). Samples were homogenized using a small Dounce tissue grinder (Sigma-Aldrich, #D8938) and incubated for 30 min on ice. Tissue homogenates were centrifuged for

15 min at 16,000*g* and 4°C, and the resulting supernatants were used as the total protein fraction. The antibodies used in this study are listed in Supplementary table 1.

For measurement of OGA expression, cytosolic proteins were extracted using the Mem-PER plus membrane protein extraction kit (Thermo Fisher Scientific, Massachusetts, US, #89842), and following the manufacturer protocol. Briefly, 80 mg of LAA powder were solubilized into 500 µL of the permeabilization buffer supplemented with protease and phosphatase inhibitor cocktails (as described above) and PUGNAc 50 µM, homogenized using a small Dounce tissue grinder and incubated for 10 min at 4°C with agitation. Tissue homogenates were centrifuged for 15 min at 16,000g and 4°C, and the resulting supernatants were used as the cytosolic protein fraction. Protein concentrations were measured based on bicinchoninic acid assay (ThermoFisher Scientific, #23227). All samples were mixed with Laemmli buffer (Bio-Rad, California, US, #1610737) complemented with DTT 100 µmol/L (Sigma-Aldrich, #646563) and heated at 95°C for 5 min, before use for immunoblotting. 20 µg of proteins were separated on 1 mm TGX acrylamide SDS-PAGE and transferred onto PVDF membrane (Bio-Rad) with Transblot Turbo system (Bio-Rad) for 7 min. Analysis was performed using Image Lab software (Bio-Rad). Each target was normalized to the corresponding stain-free intensity (total protein), except for OGA, which was performed on cytosolic protein fraction (antibody did not work on total fraction). Supplementary Table 1 presents the primary and secondary antibodies as well as the experimental strategy used to evaluate the impact of AF on the O-GlcNAcylation posttranslational modification pathway.

| **Target** | **Primary antibody** | | | **Secondary antibody** | | | **Gel** |
|---|---|---|---|---|---|---|---|
| | **Reference** | | **Dilution** | **Reference** | | **Dilution** | |
| O-GlcNAc | Anti-O-GlcNAc (RL2)-HRP mouse monoclonal IgG | ab201995 Abcam | 1/10000 in BSA 2% | none | | | 7.5% |
| OGA | Anti-MGEA/OGA rabbit polyclonal IgG | ab105217 Abcam | 1/5000 in everyBlot blocking buffer (Bio-Rad) | Goat anti-rabbit IgG | 170-6515 Bio-Rad | 1/10000 in everyBlot blocking buffer (Bio-Rad) | 7.5% |

| | | | | | | | |
|---|---|---|---|---|---|---|---|
| OGT | Anti-OGT rabbit polyclonal IgG | ab96718 Abcam | 1/1000 in everyBlot blocking buffer (Bio-Rad) | Goat anti-rabbit IgG | 170-6515 Bio-Rad | 1/10000 in everyBlot blocking buffer (Bio-Rad) | 7.5% |
| GFAT2 | Anti-GFPT2 rabbit monoclonal IgG | ab190966 Abcam | 1/5000 in everyBlot blocking buffer (Bio-Rad) | Goat anti-rabbit IgG | 170-6515 Bio-Rad | 1/10000 in everyBlot blocking buffer (Bio-Rad) | 7.5% |

**Supplementary Table 1.** GFAT2: glutamine: fructose-6-phosphate aminotransferase; O-GlcNAc: O-linked N-acetylglucosamine; OGA: O-GlcNAcase; OGT: O-GlcNAc transferase;

***Assessment of mitochondrial and cytosolic enzymes specific activity.***

Citrate Synthase (CS) was assessed as previously published [7] using 5mg of LAA homogenate. One unit of CS was defined as equal to the reduction of 1 µmole DTNB/ min, and specific activity was expressed in U/mg wet weight. Complex I activity (NADH:decylubiquinone oxidoreductase) was measured using 40µg of LAA mitochondria following Rustin et al [8]. NADH consumption was measured at 340 nm with or without rotenone (1 µmol/L) to correct for rotenone-insensitive NADH consumption. One unit of complex I was taken to be equal to the oxidation of 1 µmole of NADH per min, and specific activity was expressed in U/mg protein. Complex II activity (succinate:ubiquinone oxidoreductase) was measured as previously published [9] using 3µg of LAA mitochondria. One unit of complex II was taken to be equal to the reduction of 1 µmole DCIP per min, and specific activity was expressed in U/mg of protein. Pyruvate dehydrogenase (PDH) activity was measured according to [10] using 20 µg of LAA mitochondria. One unit of PDH was taken to be equal to the production of 1 µmole NADH per min, and specific activity was expressed in U/mg of protein. Alpha-Ketoglutarate dehydrogenase ($\alpha$-KGDH) activity was measured using 20µg of LAA mitochondria in a buffer containing (mmol/L): Hepes 50, $NAD^+$ 0.4, CoASH 0.4, Rotenone 0.005, $MgCl_2$ 10, BSA 0.04%, pH 7 (RT with KOH). Following baseline collection, reaction was started by adding 10 mmol/L final of alpha-Ketoglutarate and $\alpha$-KGDH activity was monitored through NADH production measured at 340nm. One unit of $\alpha$-KGDH was taken to be equal to the production of 1 µmole NADH per min, and specific activity was expressed in U/mg of protein. Succinyl-CoA synthetase (SCS) activity was measured using 12µg of LAA mitochondria in a buffer containing (mmol/L): Hepes 50, $MgCl_2$ 10, BSA 0.04%, ADP 1, GDP 1, $H_2PO_4$ 1, DTNB 0.15, decylubiquinone 0.05, 2.5µg/mL myxothiazol, pH 7 (RT with KOH). Following baseline collection, reaction was started by adding 250 µmol/L final of Succinyl-CoA and SCS activity was monitored through DTNB reduction measured at 412nm. One unit of SCS was taken to be equal to the reduction of 1 µmole DTNB per min, and specific activity was expressed in U/mg of protein. Fumarase activity was measured with 20 µg of LAA mitochondria added to the following buffer: Tris 50 mmol/L, KCl 150 mmol/L, pH 8, supplemented with Acetyl-CoA (100 µmol/L), $NAD^+$ (300 µmol/L), Rotenone (0.5 µmol/L), Antimycin A (0.5 µmol/L). The reaction was started by Fumarate addition (5 mmol/L) and NADH production was followed at 340 nm. One unit of Fumarase activity was taken to be equal to 1 µmole of NADH produced per min, and specific activity was expressed in U/mg protein. D and L-2-hydroxyglutarate dehydrogenases activity was measured with 20 µg of LAA mitochondria added to the following buffer: Hepes 50 mmol/L (pH 7.5), $ZnCl_2$ 0.6 µmol/L and DCIP 33 µmol/L. Following baseline recording the reaction was started by addition of 2.5 mmol/L of D/L-alpha-hydroxyglutaric acid (DL-HG, Sigma #94577). DCIP reduction was followed at 600 nm. Parallel experiments were performed in absence of D/L-HG to correct from non-specific DCIP reduction at 600 nm. One unit of D/L-2-hydroxyglutarate dehydrogenase activity was taken to be equal to 1 µmole of DCIP reduced per min, and specific activity was expressed in U/mg protein. Malate dehydrogenase isoform 2 (MDH2) activity was measured with 2µg of LAA mitochondria in a buffer containing (mmol/L): Tris-HCL 50 (pH 8), KCl 150, citrate synthase 0.5 U/mL, rotenone 0.003, acetyl-CoA 0.1, $NAD^+$ 0.4. Following baseline collection reaction was started by addition of 10 mmol/L L-malate, and MDH2 activity was recorded through NADH production by fluorescence (under stirring at 37°C, 340/460 nm, FP-8500, JASCO UK Limited). A calibration curve was generated using known quantity of malate dehydrogenase, and MDH2 activity in the sample expressed in U/mg protein.

Total creatine kinase (CK) activity was measured using 5mg LAA homogenate added to a buffer containing (mmol/L): Tris-HCl 100 (pH 7.4), $NADP^+$ 0.4, $MgCl_2$ 10, ADP 5, glucose 10, TX-100 0.3% (w/v), hexokinase 0.125 U/mL, glucose-6-phosphate dehydrogenase 0.25 U/mL. The reaction was started by PCr addition (20-mmol/L) and NADPH production was followed at 340 nm. Parallel experiments were performed without PCr addition to correct for non-specific NADPH production. One unit of CK activity was taken to be equal to 1 µmole of NADPH produced/min, and specific activity was expressed in U/mg wet weight. L-Lactate dehydrogenase (LDH) activity was measured using 130 µg of LAA homogenate in a buffer containing (mmol/L): Tris-HCl 100 (pH 7.1 at 37°C), NADH 300 µmol/L, Rotenone 1 µmol/L. Following baseline collection, reaction was started by adding 10 mmol/L pyruvate and LDH activity monitored by following NADH consumption at 340 nm.

***Samples preparation and protein digestion before Mass Spectrometry analysis.***

Protein-samples were solubilized in Laemmli buffer, and 10 µg were deposited onto SDS-PAGE gel. After colloidal blue staining, each lane was cut out from the gel and sectioned into 1 mm x 1 mm gel-pieces. Gel-pieces were destained in 25 mmol/L ammonium bicarbonate/50% ACN, rinsed twice in ultrapure water and shrunk in ACN for 10 min. After ACN removal, gel-pieces were dried at room temperature, covered with trypsin solution (10 ng/µL in 50 mM $NH_4HCO_3$), rehydrated at 4°C for 10 min, and finally incubated overnight at 37°C. Spots were then incubated for 15 min in 50 mmol/L $NH_4HCO_3$ at room temperature with rotary shaking. The supernatant was collected, and an $H_2O$/ACN/HCOOH (47.5:47.5:5) extraction solution was added onto gel-slices for 15 min. The extraction step was repeated twice. Supernatants were pooled and dried in a vacuum centrifuge. Digests were finally solubilized in 0.1% HCOOH.

***Sheep LAA metabolite content assessment.***

*Quantification of Metabolites by Liquid Chromatography Coupled to Conductimetry, UV absorbance or Mass Spectrometry Detectors:* Metabolites extracted from rat and sheep samples (see above) were separated using high-performance ion chromatography (HPIC) systems (Integrion for sheep LAA or ICS6000 for rat extracts; Thermo Fisher Scientific), both equipped with an AS11-HC-4 µm analytical column (250 × 2 mm), eluent generators, and eluant suppressors (Thermo Fisher Scientific). Separation was achieved with a discontinuous KOH gradient, as described in [11], at a flow rate of 0.38 mL/min. For sheep LAA, metabolite detection was performed by conductimetry for citrate, isocitrate, phosphocreatine, and pyruvate, or by UV absorbance at 225 nm for fumarate, 260 nm for adenylnucleotides and $NAD^+$, and 340 nm for NADH using a diode-array detector (Thermo Fisher Scientific). Metabolites were identified based on their retention times, co-injection with pure standards, and/or their UV spectral signatures. Quantification of metabolite levels was achieved using calibration curves generated with pure compounds. For rat extracts, metabolite detection was carried out using a high-resolution Orbitrap mass spectrometer (Exploris 120; Thermo Fisher Scientific) coupled to an EASY-IC ion source operating in negative ion mode at a needle voltage of -2.5 kV, with scan-to-scan lock-mass correction. Nitrogen was employed as the sheath gas, auxiliary gas, and sweep gas, set to 50, 10, and 1, respectively. Data acquisition was performed using Xcalibur 4.7 software (Thermo Fisher Scientific). Succinic acid-2,2,3,3-D4, used as an internal standard, and other metabolites of interest were quantified from full MS scans (m/z range: 70–1000) acquired at a resolution of 120,000 (at m/z = 200), using TraceFinder 5.2 software (Thermo Fisher Scientific). Metabolites were identified based on retention time, exact mass, and natural isotopic distribution, and their levels were normalized to the internal standard (succinic acid-2,2,3,3-D4). Metabolite concentrations in plasma fractions (µM) were determined using standard curves generated with pure compounds. For atrial cardiomyocytes, metabolite levels were normalized to oxalate content, which remained constant in both vehicle- and succinate-treated cells.

*Enzymatic quantification of sheep LAA metabolites:* To quantify L-malate, 80 µL of ethanol/HEPES metabolic extracts (see above) were added to a reaction buffer containing Tris (50 mM), KCl (150 mM, pH 8.0), $NAD^+$ (300 µM), acetyl-CoA (100 µM), and citrate synthase (1 U/mL). The reaction was carried out at 37°C, and NADH production was monitored by fluorescence (340/460 nm, FP-8500, JASCO UK Limited). A calibration curve was generated using known concentrations of L-malate, and the results were expressed as nmol/mg wet weight. For oxaloacetate quantification, 80 µL of ethanol/HEPES metabolic extracts were mixed with a reaction buffer containing Tris (50 mM), KCl (150 mM, pH 8.0), NADH (100 µmol/L), and fumarase (1 U/mL). After recording a baseline, the reaction was initiated by adding malate dehydrogenase (0.5 U/mL). Internal calibration was performed using known quantities of oxaloacetate (0.5–5 nmol). NADH consumption was monitored by fluorimetry (340/460 nm, FP-8500, JASCO UK Limited). Oxaloacetate content was corrected for the external oxaloacetate added after malate dehydrogenase addition and was expressed as nmol/mg wet weight. Succinate levels were measured in PCA/$KHCO_3$ extracts. A total of 80 µL of extract was incubated at 37°C in a reaction buffer containing $KH_2PO_4$ (50 mM), EDTA (1 mM), bovine serum albumin (BSA; 0.1%), and KCN (0.5 mM) (pH 7.2 at room temperature). Sequential additions of ventricular sheep mitochondria (20 µg), rotenone (1 µM), decylubiquinone (85 µM), and 2,6-dichlorophenolindophenol (DCIP, 48 µM) were performed. The reduction of DCIP was monitored spectrophotometrically at 600 nm in the presence or absence of 3-nitropropionic acid (3-NPA; 1 mM), an irreversible inhibitor of succinate dehydrogenase. Calibration curves were generated using known succinate concentrations, both with and without 3-NPA, and the succinate content was expressed as nmol/mg wet weight. Glucose-6-phosphate (G6-P) glycogen contents were assessed as described in [7] and adapted from [12]. For G6-P assay 80µL of PCA extract (see above) and for glycogen assay 400µL of

LAA homogenate were used (70mg/ml in 0.3mol/L PCA). L-Lactate content was measured using L-Lactate oxidase [13] with slight modifications. The final concentrations of the different compounds used were: Amplex red 1 µmol/L, Horseradish Peroxidase 1 U/mL, L-Lactate Oxidase 0.33 mU/mL. For the assay 10 µL of LAA metabolite extract (PCA methodology) were used and the reaction followed at 37°C under continuous stirring by fluorimetry (570/583nm, FP-8500, JASCO UK Limited). A calibration curve was performed in the same experimental condition with known quantity of L-lactate and content was expressed as nmol/mg wet weight. B-Hydroxybutyrate (β-OHB) and Acetoacetate (AcAc) content were measured according to publish methodology[14] using concentrated (350 mg/ml) and fresh metabolite extracts (PCA acidic extraction + neutralization). For β-OHB and AcAc assessment 100µL of metabolite extract were used and study were performed using fluorimetry (β-OHB, NADH production 340/460 nm) and spectrophotometry (AcAc, NADH consumption 340nm) and respective content was expressed as nmol/mg wet weight.

***Subcellular redox poise and metabolites compartmentation inferred from near-equilibrium assumption.***

The NAD-to-NADH ratios within the cytoplasm and the mitochondria were calculated assuming that lactate (LDH) and beta-hydroxybutyrate (HBDH) dehydrogenases were working close to thermodynamic equilibrium[15] and using the measured lactate-to-pyruvate and beta-hydroxybutyrate-to-acetoacetate ratios, according to the following equations:

$$\frac{NAD_{cyto}}{NADH_{cyto}} = EXP\left(-\frac{\Delta G^{\circ\prime\,LDH}}{RT}\right) \times \frac{[L - lactate]}{[Pyruvate]}$$

$$\frac{NAD_{mito}}{NADH_{mito}} = EXP\left(-\frac{\Delta G^{\circ\prime\,HBDH}}{RT}\right) \times \frac{[\beta - Hydroxybutyrate]}{[Acetoacetate]}$$

From these local redox ratios, the overall tissue redox ratio can be calculated using the following equation:

$$(NAD/NADH)tissue = \frac{\left(\frac{redox_{cyto}}{1+redox_{cyto}}\right)\cdot\left(\frac{Vol_{ratio}\times N_{ratio}}{1+Vol_{ratio}\times N_{ratio}}\right)+\left(\frac{redox_{mito}}{1+redox_{mito}}\right)\cdot\left(\frac{1}{1+Vol_{ratio}\times N_{ratio}}\right)}{\left(\frac{1}{1+redox_{cyto}}\right)\cdot\left(\frac{Vol_{ratio}\times N_{ratio}}{1+Vol_{ratio}\times N_{ratio}}\right)+\left(\frac{1}{1+redox_{mito}}\right)\cdot\left(\frac{1}{1+Vol_{ratio}\times N_{ratio}}\right)}$$

which can be simplified as the following equation:

$$(NAD/NADH)tissue = \frac{\left(\frac{redox_{cyto}\times Vol_{ratio}\times N_{ratio}}{1+redox_{cyto}}\right)+\left(\frac{redox_{mito}}{1+redox_{mito}}\right)}{\left(\frac{Vol_{ratio}\times N_{ratio}}{1+redox_{cyto}}\right)+\left(\frac{1}{1+redox_{mito}}\right)}$$

where $redox_{mito}$ and $redox_{cyto}$ are the NAD-to-NADH ratios within the mitochondria and the cytoplasm, respectively ($redox_{mito} = \frac{NAD_{mito}}{NADH_{mito}}$ and $redox_{cyt} = \frac{NAD_{cyto}}{NADH_{cyto}}$), $Vol_{ratio}$ is the cytoplasmic to mitochondrial volume ratio ($Vol_{ratio} = \frac{Vol_{cyto}}{Vol_{mito}}$) and $N_{ratio}$, the cytoplasmic to mitochondrial NAD(H) content ratio ($N_{ratio} = \frac{(NAD+NADH)_{cyto}}{(NAD+NADH)_{mito}}$). Moreover, the protomotive force was calculated assuming that the malate-aspartate shuttle was working close to thermodynamic equilibrium resulting in the following expression:

$$\Delta p = \frac{\Delta\mu H^{+}}{F} = -\frac{RT}{F}Ln\left(\frac{NAD_{cyto}}{NADH_{cyto}}\times\frac{NADH_{mito}}{NAD_{mito}}\right)$$

The distribution of the L-malate across the mitochondrial inner membrane was calculated assuming that the $malate^{2-}/Pi^{2-}$ and the $Pi^{-}$-$H^{+}$ electroneutral carriers were working close to the thermodynamic equilibrium. Under such conditions, the malate concentration ratio is related to the transmembrane pH difference, as shown in the following equation:

$$\frac{[L-malate]_{cyto}}{[L-malate]_{mito}} = e^{-2\times 2.3\times\Delta pH} \qquad \text{eq. 1}$$

According to the mass conservation law, the measured malate content of the tissue is a function of the local malate concentrations and of the subcellular volumes, as shown in the following equation:

$$L - malate_{tissue} = Vol_{cyto} \times [L - malate]_{cyto} + Vol_{mito} \times [L - malate]_{mito} \qquad \text{eq. 2}$$

Combining equations 1 and 2, one can calculate the cytoplasmic and mitochondrial malate contents, expressed as in pmol/mg ww, as shown in the following equations:

$$L - malate_{mito} = \frac{L - malate_{tissue}}{1 + \frac{Vol_{cyto}}{Vol_{mito}} . e^{2\times 2.3\times \Delta pH}}$$

$$L - malate_{cyto} = \frac{L - malate_{tissue}}{1 + \frac{Vol_{mito}}{Vol_{cyto}} . e^{-2\times 2.3\times \Delta pH}}$$

Finally, the cytoplasmic and mitochondrial oxaloacetate contents as well as the mitochondrial alpha-ketoglutarate content were calculated assuming that the cytosolic and mitochondrial malate dehydrogenases (MDH) and isocitrate dehydrogenase (IDH) were working close to the thermodynamic equilibrium resulting in the following expressions:

$$OAA_{cyto} = L - malate_{cyto} \times \frac{NAD_{cyto}}{NADH_{cyto}} \times e\left(-\frac{\Delta G^{\circ\prime\,MDH}}{RT}\right)$$

$$OAA_{mito} = L - malate_{mito} \times \frac{NAD_{mito}}{NADH_{mito}} \times e\left(-\frac{\Delta G^{\circ\prime\,MDH}}{RT}\right)$$

$$\alpha - KG_{mito} = \frac{isocitrate_{mito}}{\mathrm{CO2}_{tissue}} \times \frac{NAD_{mito}}{NADH_{mito}} \times e\left(-\frac{\Delta G^{\circ\prime\,IDH}}{RT}\right)$$

Values of parameters used in the calculations are listed in supplemental Table 1.

**Supplemental Tables**

**<u>Supplemental Table 1</u>: Values of parameters used in the calculations of redox potentials and metabolite contents.**

| | |
|---|---|
| ΔG°' LDH (kJ/mol) | -24.7 |
| ΔG°' MDH (kJ/mol) | 27.8 |
| ΔG°' HBDH (kJ/mol) | 5.2 |
| ΔG°' IDH (kJ/mol) | -11.6 |
| $CO_{2\ Tissue}$ (nmol/mg ww) | 0.617 |
| ΔpH (Sham group) | 0.40 [16] |
| ΔpH (AF-S group) | 0.35[§] |
| ΔpH (AF-S group) | 0.41[§] |
| F (C/mol) | 96845 |
| T (°K) | 310 |
| R (J/mol/K) | 8.314 |
| $N_{ratio}$ | 0.429 [17,18] |
| $V_{ratio}$ | 0.266 [19] |

Footnotes:

§ : ΔpH in AF-S and AF-R values were adjusted to the variation compared to the Sham group

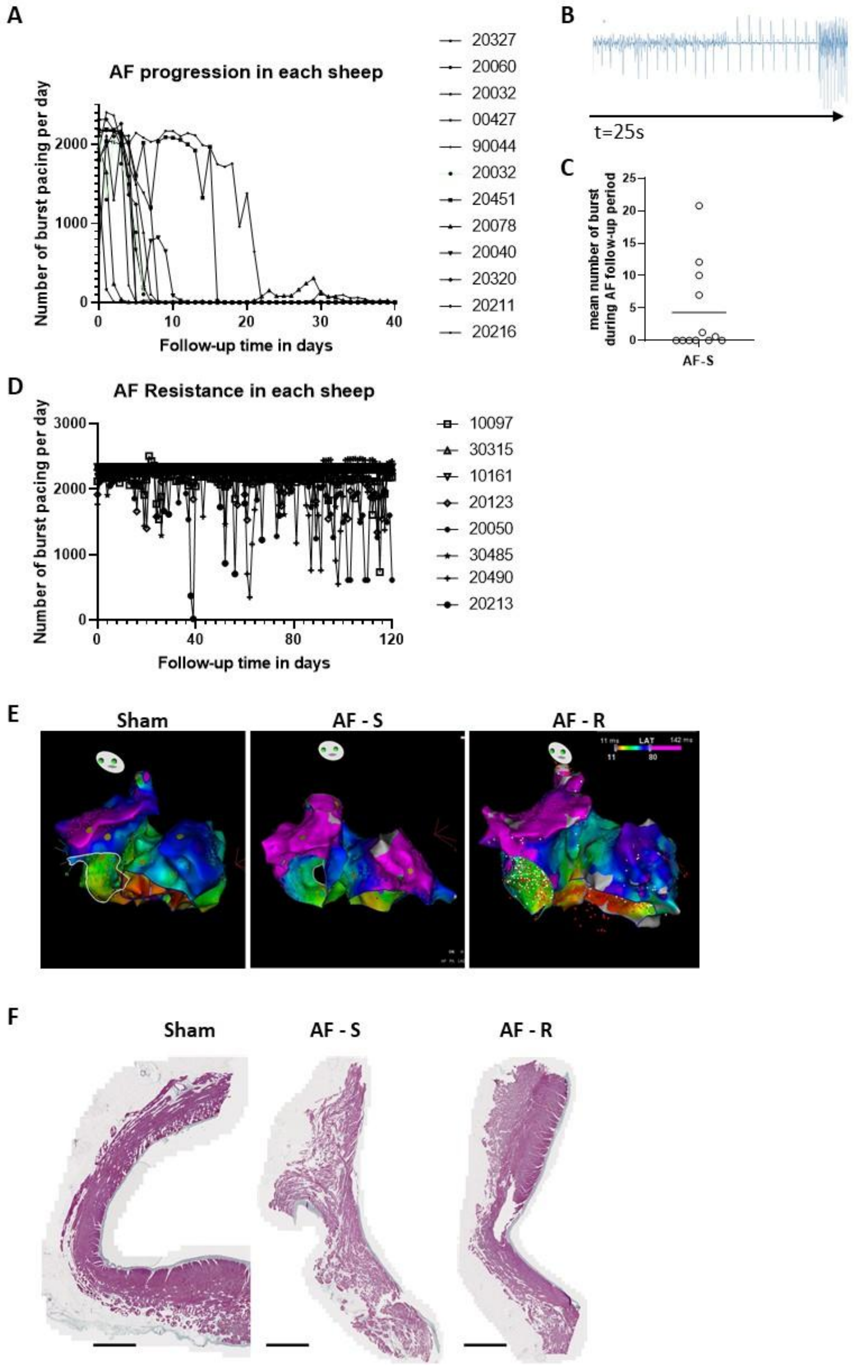


**Supplemental Figure 1. AF properties, electrophysiological and structural remodeling.**
**A**: AF progression during the first 40 days of follow-up, reflected by the number of burst pacing episodes per day, in each sheep included in the AF-S group. **B:** Typical traces of the pseudo-ECG morphology recorded by the implanted telemetry showing an episode of atrial fibrillation spontaneously converting to sinus rhythm and triggering a burst of rapid atrial pacing typically observed in sheep resistant to AF stabilization (AF-R). **C, D:** Mean number of bursts in individual AF-S sheep (**C**) and AF-R sheep (**D**) over the observation period. **E**: Representative electro-anatomical maps of the bi-atrial local activation time, for the three groups, in left anterior oblique view **F**: Representative images of left atria (LA) posterior wall histological preparations, for the three groups, after Masson trichrome stain and scanning. Scale bar = 2 mm.

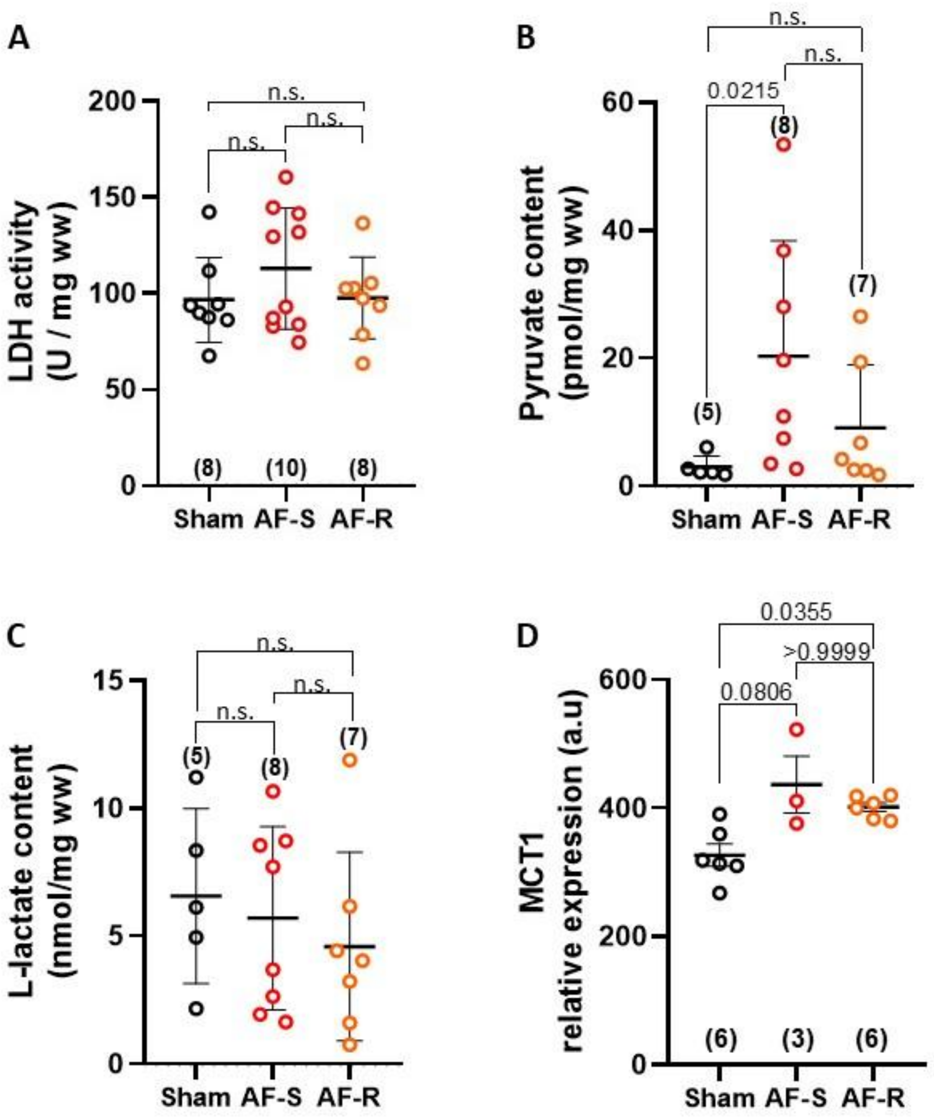


**<u>Supplemental Figure 2.</u> Schematic representing glycolytic and branched metabolic pathways remodeling**

Left Atrial Appendage (LAA) samples freeze-clamped *in vivo* were analyzed by mass spectrometry for protein identification and enzymatic assay to assess proteins activity. **A** to **D**: L-lactate dehydrogenase activity activity (LDH, panel A), Pyruvate content (G6P, panel B), L-Lactate content (panel C), and relative protein expression of MCT1 (panel D). A to D show individual points and mean ± SD. Statistical analysis was performed with Kruskal-Wallis test and Dunn’s post-hoc test for multiple comparisons. The Number in brackets represents the number of individual experiments for Sham, AF-S and AF-R groups.

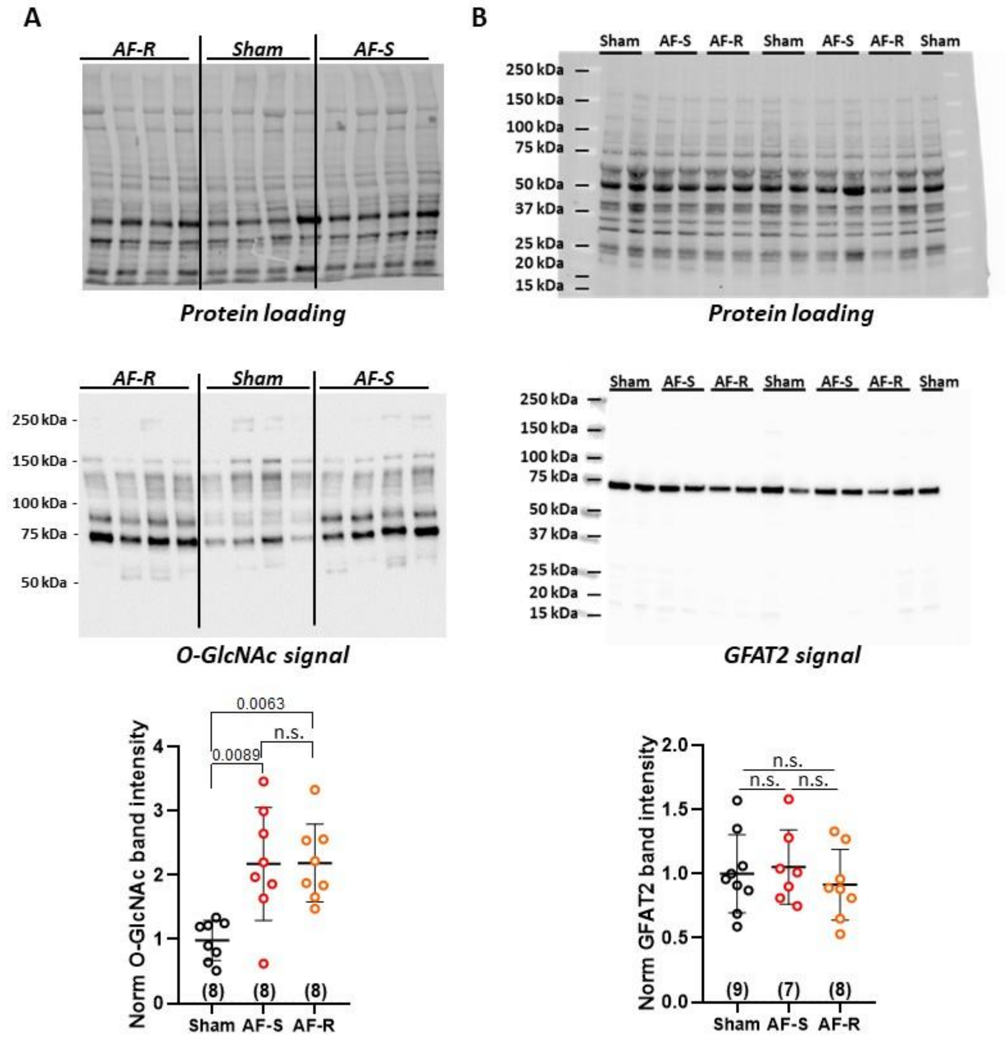


<u>**Supplemental Figure 3.**</u> **O-GlcNAcylated proteins accumulation and GFAT2 content.**
**A**: O-GlcNAcylation level of left atrial appendage proteins from Sham, AF-Sensitive (AF-S), and AF-Resistant (AF-R) sheep. Top panel: representative image showing protein loading on the gel and immunoblot of O-GlucNAcylated proteins (O-GlcNAc). Bottom panel: Individual data-points and mean±SD bars, normalized to stain free and then to mean of Sham group. **B**: Expression levels of the glutamine-fructose-6-phosphate aminotransferase (GFAT2) in the left atrial appendage of Sham, AF-S, and AF-R. Top panel: representative image showing protein loading on the gel and immunoblot of GFAT2 signals. Bottom panel: Individual data-points and mean±SD bars, normalized to stain free and then to mean of Sham group. Statistical analysis was performed by Kruskal-Wallis test and Dunn's post-hoc test for multiple comparisons. The number in brackets represents the number of individual experiments for Sham, AF-S and AF-R groups.

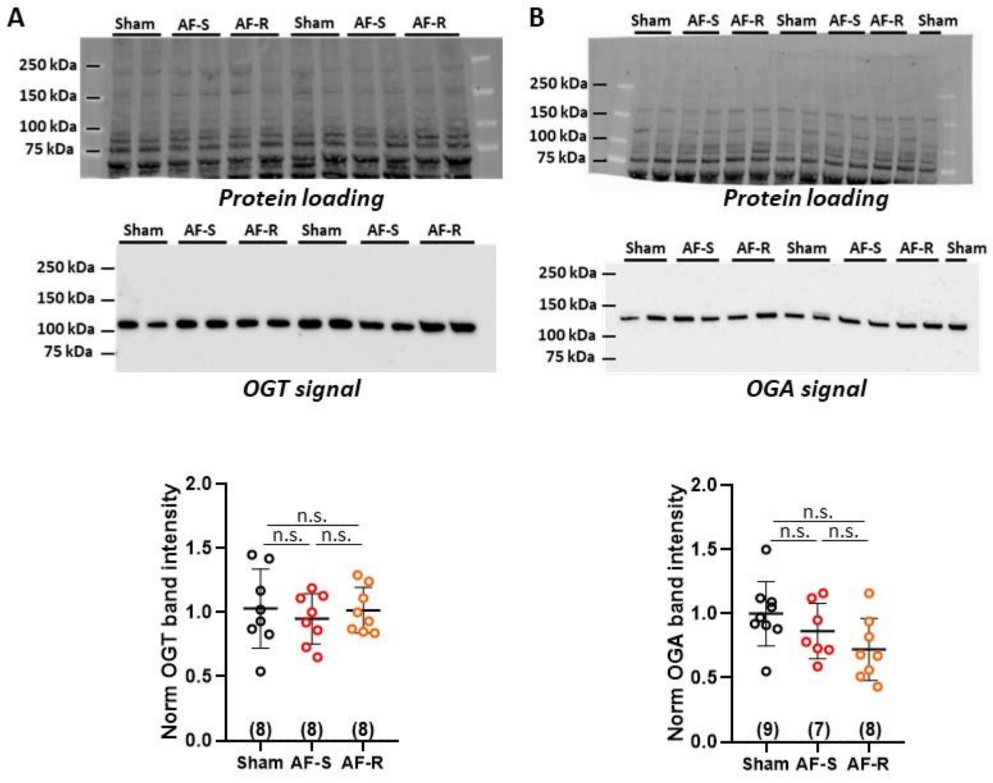


**<u>Supplemental Figure 4.</u> Expression levels of OGT and OGA.**

**A** and **B**: Expression levels of the O-GlcNAc transferase (OGT, panel **A**) and O-GlcNAcase (OGA, panel **B**), in the left atrial appendage of Sham, AF-Sensitive (AF-S), and AF-Resistant (AF-R) sheep. Top panels protein loading on the gels, middle panels OGT or OGA specific signals, and bottom panels quantification of OGT and OGA levels normalized to stain free and then Sham group mean. In the quantification plots, individual band intensity and the corresponding mean ± SD are shown for each group. Statistical analysis was performed by Kruskal-Wallis test and Dunn's post-hoc test for multiple comparisons.

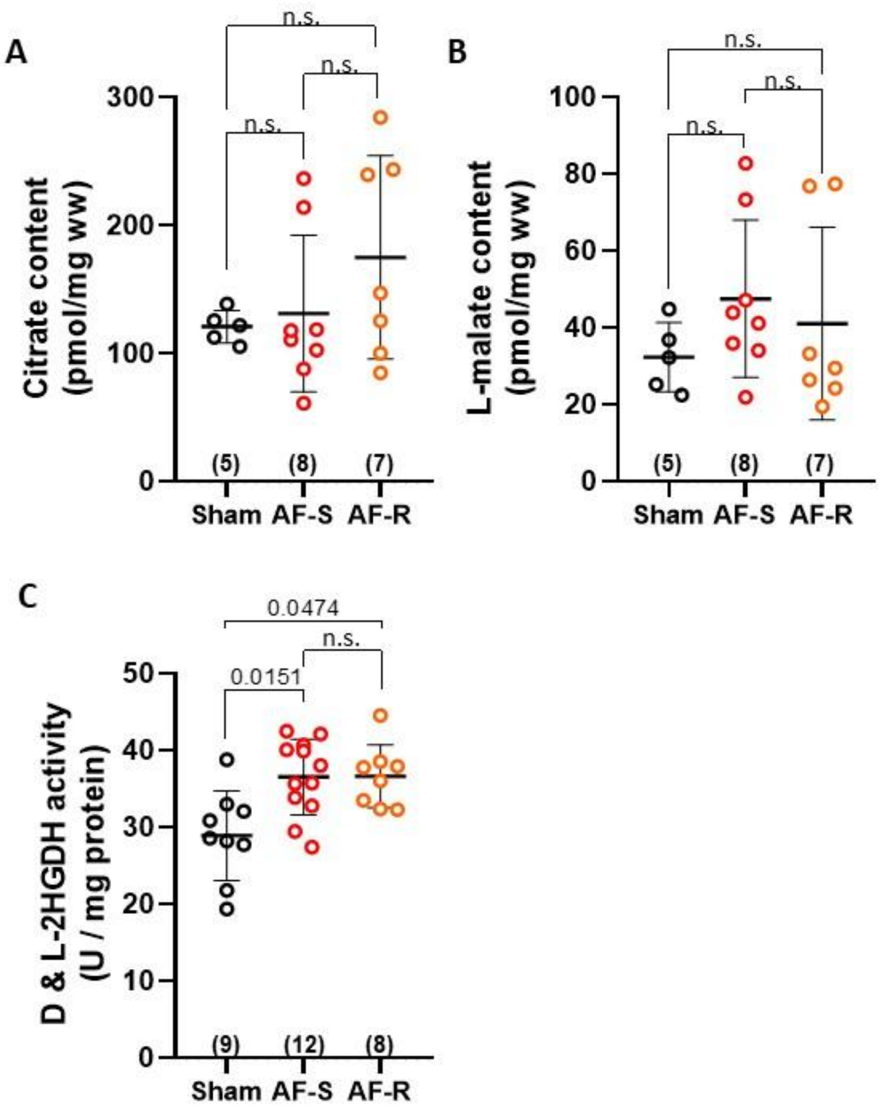


**<u>Supplemental Figure 5.</u> TCA metabolites and D,L-2HGDH activity**

Pieces of LAA freeze-clamped *in vivo* were analyzed by high-pressure ion chromatography (HPIC) for metabolites belonging to the TCA cycle. Metabolite content is expressed in pmol/mg wet weight (ww). For TCA cycle, citrate (**A**), L-malate (**B**), and fumarate (**C**) content were assessed. **D:** D,L-2-hydroxyglutarate dehydrogenase (D & L-2HGDH) activity was assessed on isolated mitochondria. Panels show individual data points and mean±SD. Statistical analysis was performed with Kruskal-Wallis test and Dunn's post-hoc test for multiple comparisons. The number in brackets represents the number of individual experiments for Sham, AF-S and AF-R groups.

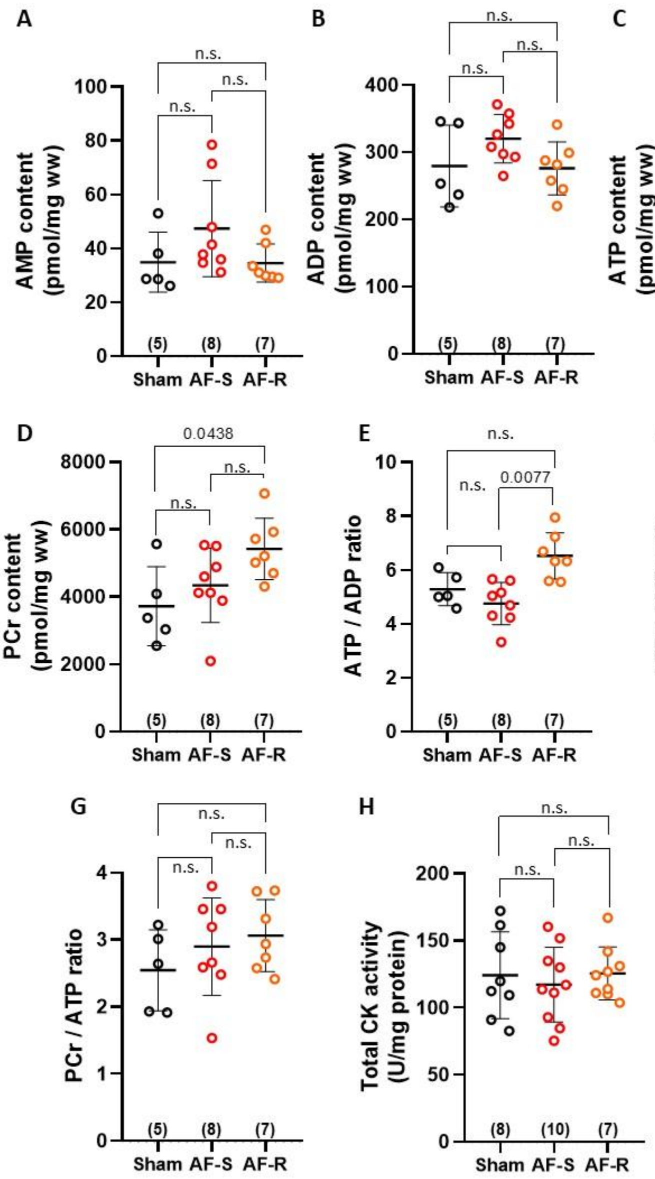


**<u>Supplemental Figure 6.</u> LAA energetic status**

Pieces of Left Atrial Appendage (LAA) freeze-clamped *in vivo* were analyzed by HPIC following ethanol/Hepes extraction for their content in AMP (**A**), ADP (**B**), ATP (**C**), and PCr (**D**). The corresponding ATP to ADP ratio (**E**), ATP to AMP ratio (**F**) and PCr to ATP ratio (**G**) were calculated to assess the energetic status of the LAA in the different groups. Creatine Kinase (CK) total activity (**H**) was measured in LAA homogenates using a spectrophotometric approach. Panels show individual points and mean ± SD. Statistical analysis was performed by Kruskal-Wallis test and Dunn's post-hoc test for multiple comparisons. The number in brackets represents the number of individual experiments for Sham, AF-S and AF-R groups.

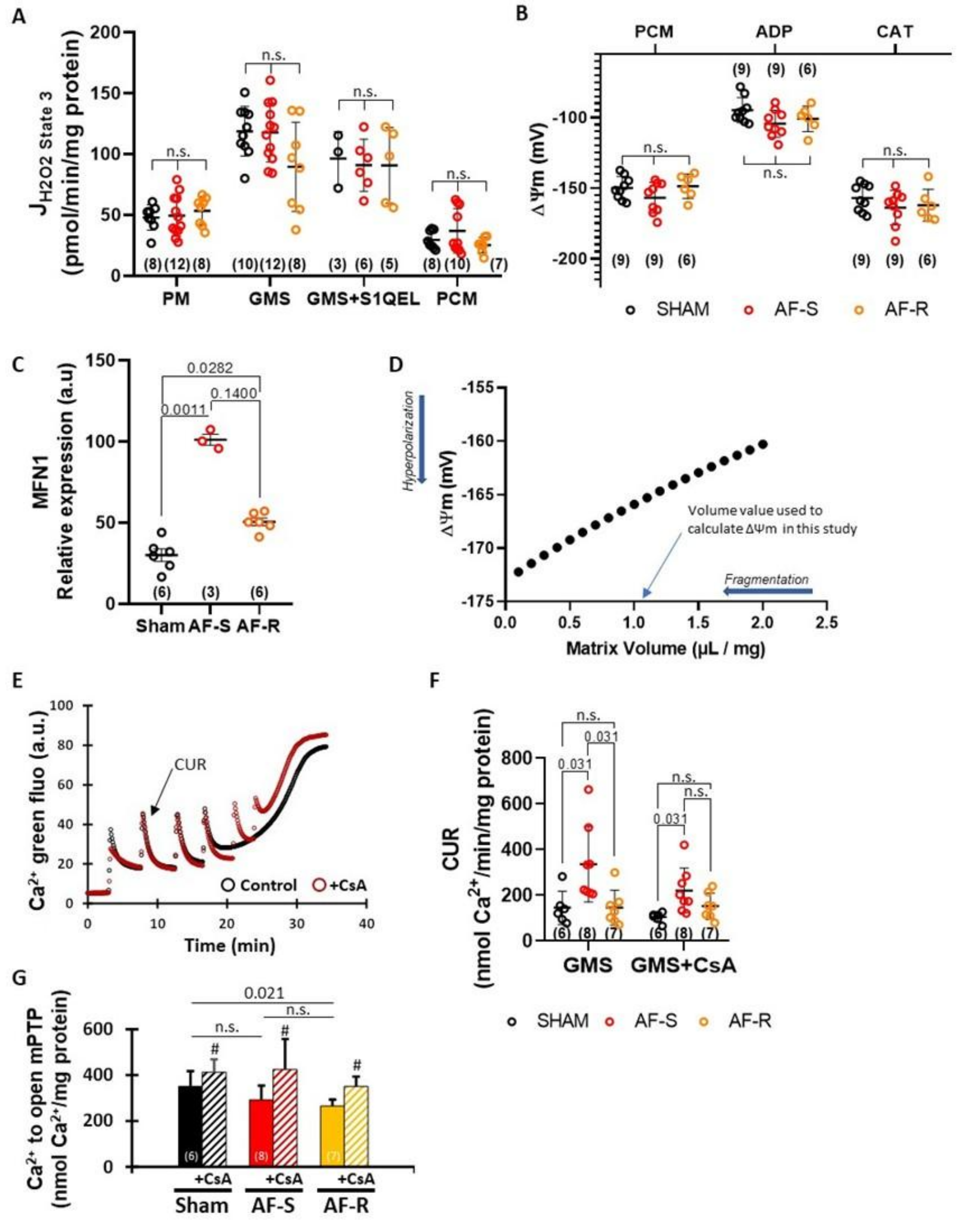


**<u>Supplemental Figure 7.</u> Mitochondrial function and calcium retention capacity assay**

Mitochondria were isolated and purified from LAA of Sham (black), AF Sensitive (red, AF-S), and AF Resistant (orange, AF-R) in order to evaluate (i) the function of the electron transport chain, (panels **A:** hydrogen peroxide emission during ATP synthesis (state 3), **B**: mitochondrial inner membrane potential (ΔΨm) during palmitoyl-carnitine + malate (PCM) oxidation alone and after ADP addition (induction of state 3) followed by inhibition by carboxyatractyloside (CAT). Panel **C**: relative expression level of the mitofusin isoform 1 studied by Mass spectrometry. Panel **D:** evolution of ΔΨm as a function of mitochondrial volume.

Panels **E** to **G**. For calcium retention capacity assay electron transport chain was fueled with glutamate + succinate + malate (GMS) with or without 200 nmol/L cyclosporin A (GMS+CsA) to inhibit mitochondrial permeability transition pore (mPTP) opening. Serial $CaCl_2$ additions of 35 µmol/L were applied until mPTP opened. **E:** Typical original recordings during the calcium retention capacity assay. **F:** Maximal calcium uptake rate (CUR). **G:** Total quantity of calcium required to open mPTP. Data points and mean ± SD are shown. Statistical analysis was performed using the Holm-Sidak method for multiple comparisons. The number in brackets represents the number of individual experiments for each experimental condition and group. # $p<0.05$ GMS+CsA *vs*. GMS paired data analyzed by Wilcoxon test. Panel C. Analysis Kruskal-Wallis test and Dunn's post-hoc test for multiple comparisons.

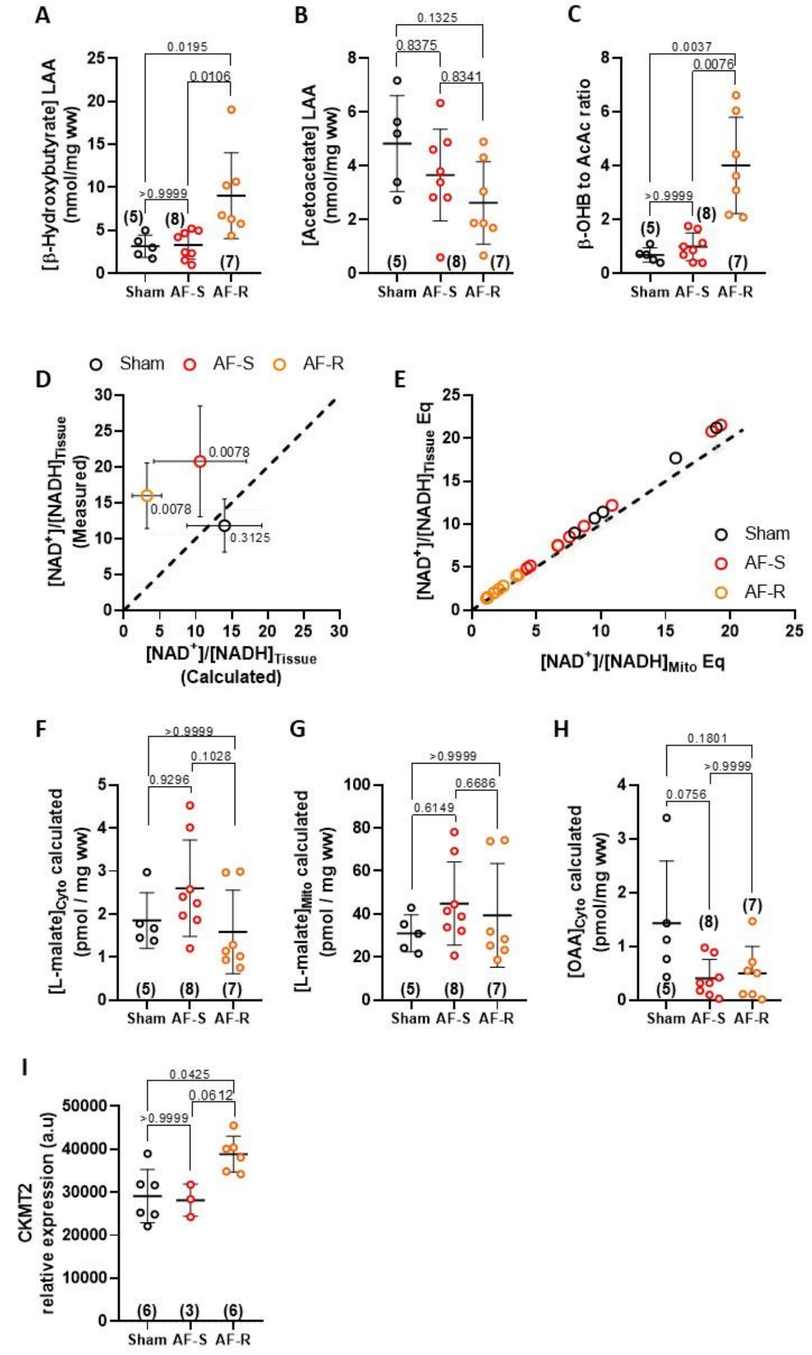


**Supplemental Figure 8. LAA redox state, calculation of the tissular NAD to NADH, and relative expression mitochondrial creatine kinase.**

Panels **A, B** and **C:** content of β-hydroxybutyrate (**A**), acetoacetate (**B**) β-hydroxybutyrate to acetoacetate ratio (**C**) measured in Left Atrial Appendage (LAA) freeze-clamped tissue *in vivo*. **D** and **E**: Graphs representing the mean measured LAA tissue $NAD^+$ to NADH ratio as a function of the mean calculated $NAD^+$ to NADH ratio (**D**), and linear correlation between the calculated LAA tissue $NAD^+$ to NADH ratio and the calculated mitochondrial $NAD^+$ to NADH ratio (**E**). Panels **F** to **H**: calculated L-malate content within the cytosolic (**F**) and mitochondrial (**G**) compartments, and calculated cytosolic oxaloacetate content (**H**). Panel **I**. LAA relative expression level of the mitochondrial creatine kinase studied by Mass spectrometry. Panels show individual data-points and mean ± SD. Statistical analysis in panels A to C and F to I, was performed by Kruskal-Wallis test and Dunn's post-hoc test for multiple comparisons. The number in brackets represents the number of individual experiments for Sham, AF-S and AF-R groups.

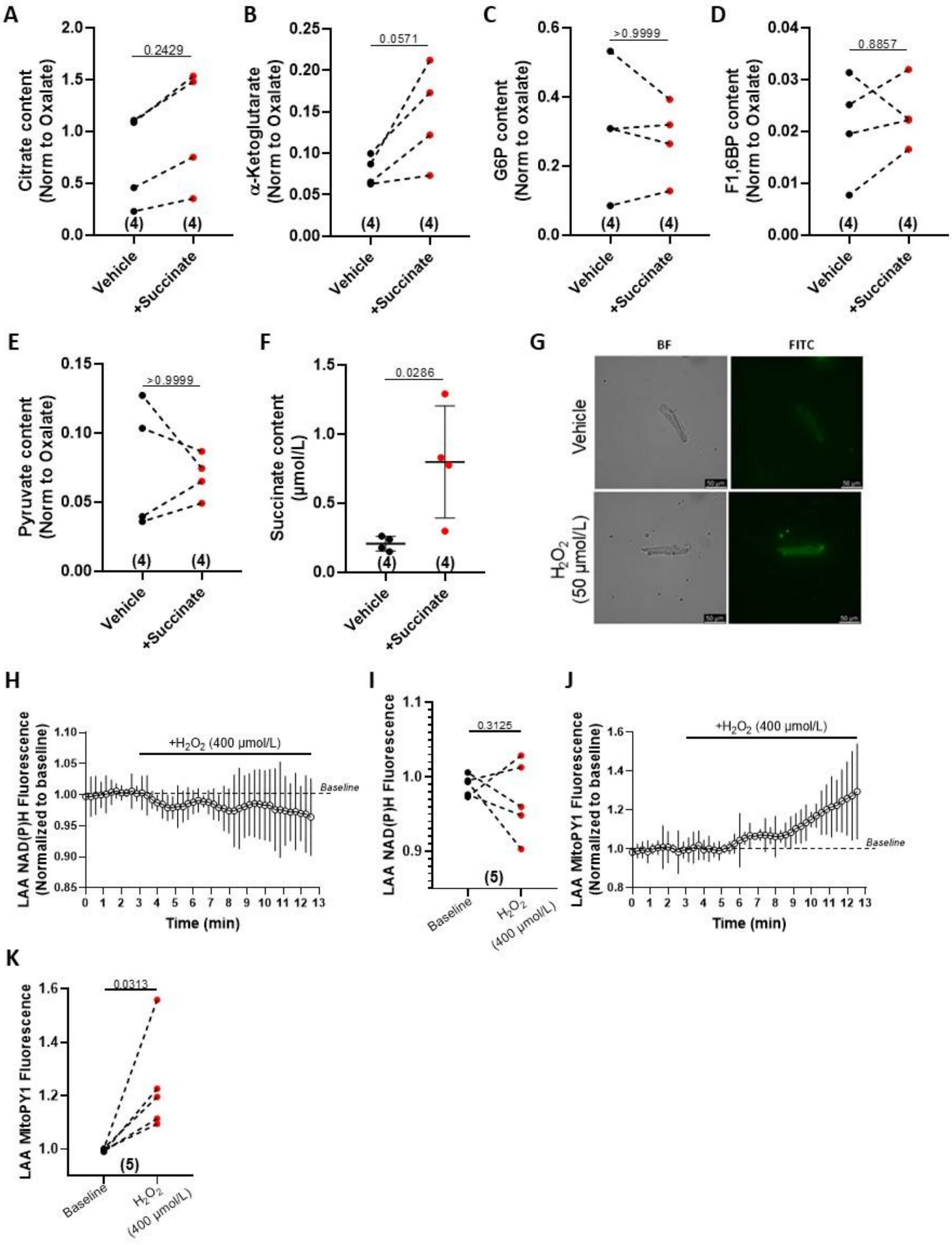


**<u>Supplemental Figure 9.</u> Effect of exogenous succinate on rat atrial cardiomyocytes metabolism**

**A-D:** Isolated rat atrial cardiomyocytes (RACMs) were incubated with 5 mmol/L of succinate and intracellular metabolite content was assessed by mass spectrometry. Signal correction-quantification and normalization were performed using deuterium-labelled succinic acid and oxalic acid, respectively (see Methods). Intracellular content of citrate (**A**), $\alpha$-ketoglutarate (**B**), Glucose-6-phosphate (G6P, **C**), Fructose-1,6-bisphosphate (F1,6BP, **D**), pyruvate (**E**), and succinate (**F**) is represented. **G:** Representative images illustrating ROS emission measurement by CellROX Green in RACMs (BF: bright field, FITC: Fluorescein). RACMs were incubated in presence of 50 µmol/L of $H_2O_2$. **H-K:** Evolution of LAA NAD(P)H time course (**H**), baseline and end-study quantification (**I**) and MitoPY1 ($H_2O_2$ sensitive fluorescent probe) time course (**J**), fluorescence baseline and end-study quantification (**K**) were measured in the *ex vivo* perfused rat heart, using an optical fiber positioned close the rat LAA, upon perfusion with 400 µmol/L of $H_2O_2$. **A-F:** Statistical analysis *via* Mann-Whitney, N=4 in each group Test. **I** and **K:** Statistical analysis *via* Wilcoxon test (N=5, paired samples).

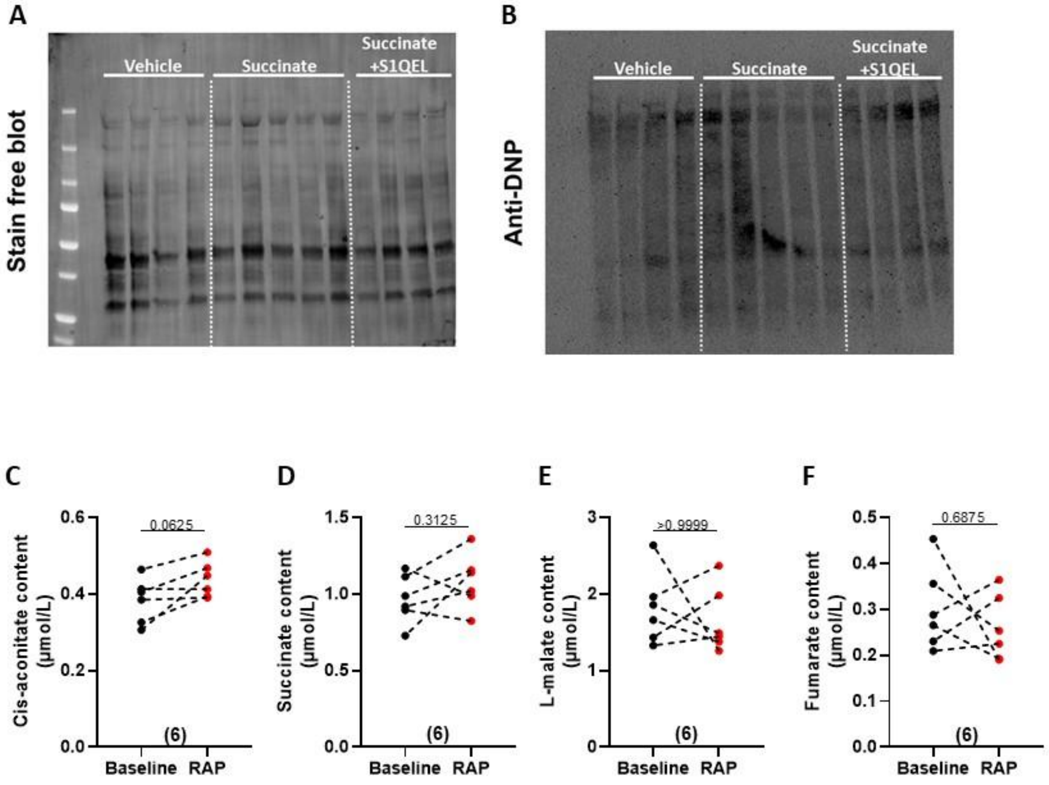


**<u>Supplemental Figure 10.</u> Carbonylation and blood metabolites in atrial tachystimulated rats.**

**A-B**: typical Oxyblot pictures obtained from rat heat treated with vehicle, succinate or succinate+S1QEL before the atrial tachystimulation challenge used to evaluate AF inducibility and stability, panel **A** stain free and panel **B** anti-DNP. **C-F**: Blood was harvested before (baseline) or after *in vivo* atrial tachystimulation by transesophageal burst-pacing (RAP), and cis aconitate (**C**), succinate (**D**), L-malate (**E**), and fumarate (**F**) were assessed by mass spectrometry (N=6). **C-F**: Statistical analysis by Wilcoxon test for paired samples. The number in brackets represents the number of individual experiments for each experimental condition.

### Supplementary References

(1) Martins, R. P.; Kaur, K.; Hwang, E.; Ramirez, R. J.; Filgueiras-Rama, D.; Ennis, S. R.; Takemoto, Y.; Ponce-Balbuena, D. Dominant Frequency Increase Rate Predicts Transition from Paroxysmal to Long-Term Persistent Atrial Fibrillation. *Circulation* **2014**, *129* (14), 1472–1482. https://doi.org/10.1161/CIRCULATIONAHA.113.004742.
(2) Schindelin, J.; Arganda-Carreras, I.; Frise, E.; Kaynig, V.; Longair, M.; Pietzsch, T.; Preibisch, S.; Rueden, C.; Saalfeld, S.; Schmid, B.; Tinevez, J.-Y.; White, D. J.; Hartenstein, V.; Eliceiri, K.; Tomancak, P.; Cardona, A. Fiji: An Open-Source Platform for Biological-Image Analysis. *Nat Methods* **2012**, *9* (7), 676–682. https://doi.org/10.1038/nmeth.2019.
(3) Pasdois, P.; Beauvoit, B.; Tariosse, L.; Vinassa, B.; Bonoron-Adèle, S.; Santos, P. D. Effect of Diazoxide on Flavoprotein Oxidation and Reactive Oxygen Species Generation during Ischemia-Reperfusion: A Study on Langendorff-Perfused Rat Hearts Using Optic Fibers. *American Journal of Physiology-Heart and Circulatory Physiology* **2008**, *294* (5), H2088–H2097. https://doi.org/10.1152/ajpheart.01345.2007.
(4) Nuutinen, E. M. Subcellular Origin of the Surface Fluorescence of Reduced Nicotinamide Nucleotides in the Isolated Perfused Rat Heart. *Basic Res Cardiol* **1984**, *79* (1), 49–58. https://doi.org/10.1007/BF01935806.
(5) Andrienko, T.; Pasdois, P.; Rossbach, A.; Halestrap, A. P. Real-Time Fluorescence Measurements of ROS and [Ca2+] in Ischemic / Reperfused Rat Hearts: Detectable Increases Occur Only after Mitochondrial Pore Opening and Are Attenuated by Ischemic Preconditioning. *PLoS ONE* **2016**, *11* (12), e0167300. https://doi.org/10.1371/journal.pone.0167300.
(6) Brandes, R.; Figueredo, V. M.; Camacho, S. A.; Baker, A. J.; Weiner, M. W. Quantitation of Cytosolic [Ca2+] in Whole Perfused Rat Hearts Using Indo-1 Fluorometry. *Biophysical Journal* **1993**, *65* (5), 1973–1982. https://doi.org/10.1016/S0006-3495(93)81274-8.
(7) Pasdois, P.; Parker, J. E.; Halestrap, A. P. Extent of Mitochondrial Hexokinase II Dissociation During Ischemia Correlates With Mitochondrial Cytochrome c Release, Reactive Oxygen Species Production, and Infarct Size on Reperfusion. *JAHA* **2013**, *2* (1), e005645. https://doi.org/10.1161/JAHA.112.005645.
(8) Rustin, P.; Chretien, D.; Bourgeron, T.; Gérard, B.; Rötig, A.; Saudubray, J. M.; Munnich, A. Biochemical and Molecular Investigations in Respiratory Chain Deficiencies. *Clinica Chimica Acta* **1994**, *228* (1), 35–51. https://doi.org/10.1016/0009-8981(94)90055-8.
(9) Pasdois, P.; Beauvoit, B.; Tariosse, L.; Vinassa, B.; Bonoron-Adèle, S.; Santos, P. D. MitoK ATP -Dependent Changes in Mitochondrial Volume and in Complex II Activity during Ischemic and Pharmacological Preconditioning of Langendorff-Perfused Rat Heart. *J Bioenerg Biomembr* **2006**, *38* (2), 101–112. https://doi.org/10.1007/s10863-006-9016-3.
(10) Hinman, L. M.; Blass, J. P. An NADH-Linked Spectrophotometric Assay for Pyruvate Dehydrogenase Complex in Crude Tissue Homogenates. *J Biol Chem* **1981**, *256* (13), 6583–6586.
(11) Pinson, B.; Moenner, M.; Saint-Marc, C.; Granger-Farbos, A.; Daignan-Fornier, B. On-Demand Utilization of Phosphoribosyl Pyrophosphate by Downstream Anabolic Pathways. *J Biol Chem* **2023**, *299* (8), 105011. https://doi.org/10.1016/j.jbc.2023.105011.
(12) Passonneau, J. V.; Lauderdale, V. R. A Comparison of Three Methods of Glycogen Measurement in Tissues. *Anal Biochem* **1974**, *60* (2), 405–412. https://doi.org/10.1016/0003-2697(74)90248-6.

(13) Bouchez, C. L.; Daubon, T.; Mourier, A. NADH-Independent Enzymatic Assay to Quantify Extracellular and Intracellular L-Lactate Levels. *STAR Protoc* **2022**, *3* (2), 101403. https://doi.org/10.1016/j.xpro.2022.101403.
(14) Galán, A.; Hernández, J.; Jimenez, O. Measurement of Blood Acetoacetate and Beta-Hydroxybutyrate in an Automatic Analyser. *J Autom Methods Manag Chem* **2001**, *23* (3), 69–76. https://doi.org/10.1155/S1463924601000086.
(15) Williamson, D. H.; Lund, P.; Krebs, H. A. The Redox State of Free Nicotinamide-Adenine Dinucleotide in the Cytoplasm and Mitochondria of Rat Liver. *Biochem J* **1967**, *103* (2), 514–527. https://doi.org/10.1042/bj1030514.
(16) Thomas, P. J.; Gaspers, L. D.; Pharr, C.; Thomas, J. A. Continuous Measurement of Mitochondrial pH Gradients in Isolated Hepatocytes by Difference Ratio Spectroscopy. *Arch Biochem Biophys* **1991**, *288* (1), 250–260. https://doi.org/10.1016/0003-9861(91)90192-l.
(17) Alano, C. C.; Tran, A.; Tao, R.; Ying, W.; Karliner, J. S.; Swanson, R. A. Differences among Cell Types in NAD(+) Compartmentalization: A Comparison of Neurons, Astrocytes, and Cardiac Myocytes. *J Neurosci Res* **2007**, *85* (15), 3378–3385. https://doi.org/10.1002/jnr.21479.
(18) Stein, L. R.; Imai, S. The Dynamic Regulation of NAD Metabolism in Mitochondria. *Trends Endocrinol Metab* **2012**, *23* (9), 420–428. https://doi.org/10.1016/j.tem.2012.06.005.
(19) Aliev, M. K.; Dos Santos, P.; Hoerter, J. A.; Soboll, S.; Tikhonov, A. N.; Saks, V. A. Water Content and Its Intracellular Distribution in Intact and Saline Perfused Rat Hearts Revisited. *Cardiovasc Res* **2002**, *53* (1), 48–58. https://doi.org/10.1016/s0008-6363(01)00474-6.
